# Dynamic Changes in Gene Expression Through Aging in *Drosophila melanogaster* Heads

**DOI:** 10.1101/2024.12.11.627977

**Authors:** Katherine M Hanson, Stuart J Macdonald

## Abstract

Work in many systems has shown large-scale changes in gene expression during aging. However, many studies employ just two, arbitrarily-chosen timepoints at which to measure expression, and can only observe an increase or a decrease in expression between “young” and “old” animals, failing to capture any dynamic, non-linear changes that occur throughout the aging process. We used RNA sequencing to measure expression in male head tissue at 15 timepoints through the lifespan of an inbred *Drosophila melanogaster* strain. We detected >6,000 significant, age-related genes, nearly all of which have been seen in previous fly aging expression studies, and which include several known to harbor lifespan-altering mutations. We grouped our gene set into 28 clusters via their temporal expression change, observing a diversity of trajectories; some clusters show a linear change over time, while others show more complex, non-linear patterns. Notably, re-analysis of our dataset comparing the earliest and latest timepoints – mimicking a two-timepoint design – revealed fewer differentially-expressed genes (around 4,500). Additionally, those genes exhibiting complex expression trajectories in our multi-timepoint analysis were most impacted in this re-analysis; Their identification, and the inferred change in gene expression with age, was often dependent on the timepoints chosen. Informed by our trajectory-based clusters, we executed a series of gene enrichment analyses, identifying enriched functions/pathways in all clusters, including the commonly seen increase in stress- and immune-related gene expression with age. Finally, we developed a pair of accessible shiny apps to enable exploration of our differential expression and gene enrichment results.

## INTRODUCTION

Aging is marked by both a decline in organismal function and an increased risk for disease. As complex traits, both environmental and genetic factors contribute to variation in lifespan and health at old age. Identifying these factors, and their impact on life- and healthspan, enables understanding of the negative effects of aging, which is increasingly important as populations throughout the world age. In humans, the estimated heritability of longevity is 7-30% (Herskind et al. 1996; Kaplanis et al. 2018; Mayer 1991; Ruby et al. 2018), and genome wide association studies (GWAS) have been used to study the genetic basis of aging (Broer et al. 2015; Deelen et al. 2011; 2014; Joshi et al. 2016; 2017; Melzer, Pilling, and Ferrucci 2020; Newman et al. 2010; Sebastiani et al. 2012; 2017; Pilling et al. 2016; 2017; Timmers et al. 2019; Zeng et al. 2016). GWAS often rely on examining sets of individuals who are >90 years old (Deelen et al. 2011; 2014; Newman et al. 2010; Sebastiani et al. 2012; 2017; Zeng et al. 2016), but this can constrain the sample size and power of such studies (Newman and Murabito 2013; Tan et al. 2008). Creative approaches have enabled markedly increased sample size – for instance, Joshi et al. (2016) associate offspring genotype with parental lifespan phenotype – yet human lifespan GWAS have only discovered a handful of replicable associations, for instance near APOE, FOXO3, and CHRNA3/5 (Broer et al. 2015; Deelen et al. 2011; Joshi et al. 2016; 2017; Pilling et al. 2016; Timmers et al. 2019).

Outside of a deficit of power in genetic studies, studying aging directly in humans can be difficult due to uncontrollable environmental variables, many of which can impact lifespan. For example, calorie restriction has been shown to increase lifespan and health in model organisms (Cypser, Kitzenberg, and Park 2013; Mattison et al. 2017; McCay et al. 1939; Pletcher et al. 2002), and in humans a two-year calorie restriction study showed a reduction in cardiometabolic risk factors including cholesterol and blood pressure (Kraus et al. 2019). However, effectively studying the impact of diet on lifespan in humans is challenging due to the difficulty accurately measuring calorie intake, the lack of adherence by participants, and several other concerns (F. B. Hu 2024). Ideally, studies investigating the genetic and molecular contributions to aging would limit all other sources of variation, something that is not feasible or practical in human studies.

Model organisms have proven useful conduits to study aging due to their shorter lifespans, ease of testing many individuals, and the ability to control both environmental and genetic variation. Both genetic and environmental contributors to aging have been successfully identified in vertebrate model systems (Cai, Wu, and Huang 2022; Gocmez et al. 2020; McCay et al. 1939), in invertebrates such as *Drosophila melanogaster* and *Caenorhabditis elegans* (Highfill et al. 2017; Kenyon 2010; McCarroll et al. 2004; Pletcher et al. 2002), and in the fungi *Saccharomyces cerevisiae* and *Podospora anserina* (Z. Hu et al. 2014; Philipp et al. 2013). These models have helped identify and understand lifespan-associated genes and systems that have relevance for human populations. For instance, FOXO3 – a member of the forkhead box transcription factor O (FOXO) family – has been implicated in human aging via GWAS (Broer et al. 2015; Deelen et al. 2014; Willcox et al. 2008), and studies in *C. elegans* and *D. melanogaster* have also identified and characterized FOXO family genes that impact lifespan (Alic et al. 2014; Giannakou et al. 2007; Kimura et al. 1997; Lin et al. 1997; Taormina et al. 2019).

Cellular processes do not remain static over an individual’s lifetime, and instead change, adapt, and sometimes break down over time. By measuring these changes, we can understand the cellular and physiological phenomena that are affected by aging. A common strategy to assess such age-related changes is with the use of gene expression analyses. These analyses often compare the genomewide gene expression profile between “young” and “old” individuals, and identify genes that are differentially expressed with age. This approach has been successfully employed in multiple systems (Bordet et al. 2021; Lu et al. 2004; X. Wang et al. 2022; Wilson et al. 2015), and has resulted in expression-based hallmarks of aging (Frenk and Houseley 2018), including increased expression of stress response and immunity genes (Bajgiran et al. 2021; Bordet et al. 2021; Lu et al. 2004; X. Wang et al. 2022), and decreased expression of genes associated with mitochondria and the electron transport chain (Bordet et al. 2021; Lu et al. 2004).

A challenge with two-timepoint, young versus old comparisons is that there is little consistency in the definitions of “young” and “old”, making comparisons across studies difficult. For instance, various studies in *D. melanogaster* have used “young” flies sampled between 1 and 10 days old, and “old” flies that were 35 to 61 days old (Bajgiran et al. 2021; Bordet et al. 2021; Carnes et al. 2015; Girardot et al. 2006; Landis et al. 2004). While in some cases the “old” timepoint is determined by the survivorship of the aging cohort, rather than by chronological age (Doroszuk et al. 2012; Highfill et al. 2017; Landis et al. 2004). Additionally, two-timepoint expression analyses are limited to making binary conclusions about expression change; expression either increases or decreases with age. Aging is a complex process and this simplification may often fail to yield an accurate picture of age-related expression for most genes. Indeed, work in several systems assaying various molecular phenotypes at multiple timepoints throughout lifespan have identified many genes and gene products with non-linear trajectories (Gheorghe et al. 2014; Haustead et al. 2016; Lund et al. 2002; Pletcher et al. 2002; Olecka et al. 2024; Remondini et al. 2010; Schaum et al. 2020; Shen et al. 2024; Xie et al. 2022).

Here we set out to robustly measure expression trajectories throughout the adult lifespan of an inbred *D. melanogaster* strain, focusing on head tissue from males. Flies were aged for 59 days, and sampled for RNAseq at 15 timepoints. We recorded the number of deaths in our aging cohort daily, allowing us to calculate a survival-based “physiological age” in addition to the “chronological age” for each time point, and endeavored to sample roughly evenly through both metrics. Subsequently, using multiple analyses we identified genes whose expression changed with aging, and clustered differentially expressed genes based on their expression trajectories. With the idea that genes with similar expression trajectories may share similar functional roles (e.g. Zhang et al. 2004), we did a series of enrichment analyses for each of our clusters, identifying numerous enriched pathways. To better understand the value of a multi-timepoint expression study, we subsequently re-analyzed our data in a two-timepoint, young versus old analysis framework. Finally, we compared the results of our work with a series of prior expression-based *D. melanogaster* aging studies.

## MATERIALS AND METHODS

### Fly rearing, maintenance, and aging

We employed a single inbred *D. melanogaster* strain, A4, which is one of the founders of the *Drosophila* Synthetic Population Resource (E. G. King et al. 2012; Chakraborty et al. 2018), a panel of recombinant strains enabling the genetic characterization of complex traits. Following several generations of expansion, we generated 200 replicate vials of the A4 strain, clearing adults to maintain roughly even egg/larval densities over vials. In the following generation we harvested 0-2 day old A4 males over CO_2_ anesthesia, collecting 132 vials of 40 males (for subsequent collection of aged animals for RNAseq), and 5 vials of 10 males (for collection of 3 day old flies for RNAseq). The aging cohort was maintained in vials, flies were tipped to new vials every 2-3 days without anesthesia, and periodically the entire cohort was anesthetized and re-arrayed into groups of 40 animals in vials. Dead animals were counted daily. Supplementary Table A provides further description of how the aging cohort was treated.

Flies were raised and maintained in standard narrow fly vials (Fisher Scientific, AS515) containing ∼10-ml of a cornmeal-yeast-molasses media (see Supplementary Text A), and kept in an incubator under the following environmental conditions: 25°C, 50% relative humidity, and a 12:12 Light:Dark cycle.

### Fly/tissue sampling

Multiple groups of flies were sampled from the population through the aging process, with animals for the first – Day 3 – timepoint coming from those vials initially holding 10 flies (see above). Flies were sampled without anesthesia via manual aspiration, and sampled flies were always given at least 24 hours between CO_2_ anesthesia and sampling. See Supplementary Table A for details on when flies were sampled.

On collection, groups of 10 male flies were moved into screw-top tubes, flash frozen in liquid nitrogen, and kept at −80°C. Subsequently – after the entire cohort had died and flies from all timepoints had been sampled – tubes were removed from the freezer into aluminum dry bath blocks held on dry ice. We then went through the tubes one by one, subjected each to liquid nitrogen, briefly vortexed to separate heads/bodies, poured the body parts into the flat surface of an aluminum dry bath block placed on dry ice, and used a paint brush to manually collect heads into a fresh screw-top tube. These destination tubes were pre-filled with 4-6 glass beads (BioSpec Products, 11079127) that had been previously washed in bleach and thoroughly rinsed with distilled water.

### RNA isolation, sequencing library preparation, and sequencing

RNA was isolated using the Zymo Direct-zol MicroPrep kit (Zymo, R2062), largely following the manufacturer’s protocol (see Supplementary Text B). Samples were isolated over 6 batches, replicate samples from a given timepoint were isolated in different batches, and RNA quantity was measured using a NanoDrop ND-1000.

Forty-eight RNA samples were used to generate mRNA sequencing libraries; 14 timepoints had 3 replicates each, while one (Day 59, the final timepoint) had 6 replicates. We used 200-500ng of total RNA from these samples to initiate half-reaction volume mRNA sequencing library construction (Illumina TruSeq stranded HT kit using dual indexing), generating libraries across 2 batches of 24 samples each. Libraries were quantified using a Qubit fluorometer, and a subset of 8 libraries from each batch were run on an Agilent TapeStation. Each of these libraries showed a single library peak, no evidence of adapter dimers, and estimated average fragment sizes of 277-294bp. See Supplementary Table B for details on RNA isolation / library preparation batching and quantification. Given the relatively uniform fragment sizes, equal quantities of all 48 libraries were pooled together. The final 48-plex pool had an average fragment size of 289bp, and was run over two Illumina NextSeq550 PE75 flowcells, yielding a total of over 635 million read pairs, with an average of 13.2 million per sample (range = 9.0 – 17.4 million).

### Quantifying expression level and identifying expression changes during aging

Reads from each of the 48 samples were processed using Salmon (Patro et al. 2017), employing release BDGP6.32 of the *Drosophila melanogaster* transcriptome/annotation from Ensembl. Salmon quantifications were then summarized to gene level using R/tximeta (Love et al. 2020). To identify genes whose expression significantly changed with aging (adjusted *p*-value < 0.05) we used R/DESeq2 (Love, Huber, and Anders 2014), executing three different analyses. First, we identified genes associated with chronological age (the “Day analysis”) by treating the age of the flies in each RNAseq sample as a continuous variable. Second, we identified genes associated with physiological age (the “Survival analysis”) by using the fraction of dead animals in the entire cohort at the point flies were sampled for RNAseq as a continuous variable (Supplementary Table C). Third, we identified genes showing expression variation through aging by considering the 15 sampling points as levels of a categorical variable (the “Sampling Point analysis”). This final analysis sought to identify genes with expression patterns not easily captured by the two continuous variables (day and survival).

### Clustering genes by their expression trajectories

Our three differential expression analyses (see above) collectively identified 6,142 unique genes, and we sought to cluster these genes into groups based on their expression trajectories through aging. To focus on expression trajectories, and avoid confounding with varying expression levels, we standardized the expression counts of each gene via *z*-scores. Briefly, for each gene we calculated the mean expression across replicates for each sampling point, along with the overall mean expression across all sampling points and replicates. We then subtracted the overall mean from the mean of each sampling point and divided by the overall standard deviation.

The calculated *z*-scores for the 6,142 genes were used to create a dissimilarity matrix using the Pearson correlation method in R/factoextra (Kassambara and Mundt 2020). We used this dissimilarity matrix for hierarchical clustering of our identified genes, and then “cut” the resulting dendrogram into a designated number of gene groups/clusters. This was done using the hclust and cutree functions from the R/stats package (R Core Team 2021).

There are a variety of ways to cut a gene dissimilarity matrix into clusters, and variation in the approach and parameters will yield different numbers and sizes of clusters. Our goal was to examine whether clusters of genes with similar expression trajectories were enriched for particular properties (e.g., gene ontology terms). To facilitate this, we sought to avoid clusters with either very small or very large numbers of genes, so targeted clusters with between 50 and 500 genes. After exploring several methods, we grouped our 6,142 differentially expressed genes into 28 clusters (see Supplementary Table D and Supplementary Figure 3 for more information).

### Summarizing and classifying cluster expression trajectories

For each of the 28 clusters we created a representative expression curve by smoothing the mean *z*-score from all genes in the cluster for each sampling point (Figure 3). The smoothing was executed using geom_smooth from R/ggplot2 (Wickham 2016) and spline modeling with rcs from R/rms (Harrell Jr 2023). The resulting 28 curves show a diversity of trajectories, with some being generally linear, while others show a more complex pattern. To classify the cluster trajectories, we ran a linear regression between the mean *z*-scores and the age of the sampled flies. A trajectory was designated as “Linear” if the *p*-value was less than 0.002 (0.05/28), or “Complex” otherwise. (Repeating this analysis using survival, or the numbered sampling points, 1-15, yields the same designations). Linear trajectories were further designated as “Up” (gene expression increases with aging) or “Down” (gene expression decreases with aging) based on the sign of the linear regression coefficient. To simplify subsequent discussion, clusters are named with these classifications (i.e., Complex, LinearUp or LinearDown), and given numeric codes based on the cluster position within the dendrogram (bottom to top in Figure 2 and left to right in Supplementary Figure 4). See Supplementary Table E for details on the trajectory-designating linear regression analyses.

### Cluster-specific enrichment analyses

To understand whether genes with similar expression trajectories share similar properties/functions, we used PANGEA (Version 1.1 beta December 2022) (Y. Hu et al. 2023), an online gene set enrichment tool that can perform Gene Ontology (GO) analysis, identify enrichment of particular gene groups or pathways, and – importantly for our needs – can execute analyses on multiple gene lists simultaneously. For each cluster we ran 6 separate enrichment analyses, examining 3 *Drosophila* GO Subsets (SLIM2 GO BP – biological process, SLIM2 GO CC – cellular component, SLIM2 GO MF – molecular function), 2 collections of gene groups (DRSC GLAD and FlyBase), and the REACTOME pathway set. For each analysis we used a custom background set of 13,303 genes that included only those with at least one mapped read in our dataset. We identified terms significantly enriched in each cluster using a Benjamini Hochberg false discovery rate (FDR) of 0.05. Each of the PANGEA tables are available as Supplementary Tables F-K and our enrichment analysis code is available at https://github.com/Hanson19/RNAseq-Aging.

### Comparing trajectory-based, multi-timepoint analysis results to analyses contrasting groups of young and old animals

Our design differs from some previous examinations of age-related expression in that we generated expression data from many points through the aging process. To examine what might be gained from our approach, we re-analyzed our data after dropping the bulk of the timepoints. The samples from Day 3 and Day 6 (N=6) collectively made up our “young” sample, while our “old” sample came from the last collection day, Day 59 (N=6). Using R/DESeq2 (Love, Huber, and Anders 2014) we identified genes whose expression changed significantly between these age groups, and determined if gene expression increased or decreased over time. We subsequently repeated this analysis, comparing the Day 3+6 “young” timepoint against every sequential pair of older timepoints (e.g., we compared Day 3+6 to Day 10+14, Day 3+6 to Day 14+17, and so on).

### Comparison with previous *Drosophila* aging genomewide expression studies

We compared our set of 6,142 multi-timepoint significant genes with those identified in 8 previously published, two-timepoint aging expression studies (Supplementary Table M). These papers vary in the strains/populations employed, the sex of the animals targeted, the tissue that was employed, and the actual timepoints during aging that were sampled. We validated all gene IDs using FlyBase (FB2024_01) (Jenkins et al. 2022), identified genes shared between our study and these previous works, and compared the expression change reported in the previous studies (up or down in expression with age) with the expression trajectories these genes were grouped into in our study (LinearUp, LinearDown, Complex; Supplementary Figure 14).

### Shiny apps to enable data exploration

Our analyses generated a considerable amount of data, and to make our results more accessible, we developed two interactive apps using R/shiny (version 1.8.0) (Chang et al. 2023). The Gene and Cluster app allows users to look up specific genes, receive information about the whether the gene was identified in our analysis, and if so in which cluster it was found, and what its expression trajectory is over time. The Cluster Enrichment app allows users to select specific clusters, or sets of clusters, and identify any enriched terms. For more information on how to run and use these apps see Supplementary Text C and D.

### Data availability

The A4 strain is available on request from the corresponding author. Raw FASTQ sequencing data is available from the NCBI SRA under BioProject accession number PRJNA1194574. All summary data and results are presented in supplementary files (available at WILL_INSERT_G3_FIGSHARE_SITE_URL_HERE), and all analysis code is available via GitHub (https://github.com/Hanson19/RNAseq-Aging).

## RESULTS AND DISCUSSION

### Over 6000 genes show expression change during aging

We aged a cohort of 5,330 *Drosophila melanogaster* males from a single inbred strain, recorded the number of fly deaths each day, and collected several replicates of 10 flies at 15 timepoints throughout the adult lifespan. Samples were collected every ∼4 days between day 3 (99.9% flies alive) and day 59 (1.92% alive) to roughly evenly sample flies throughout lifespan (see Kaplan-Meier survivorship curve in Supplementary Figure 1 and detail on the timing of the sampling in Supplementary Table C). RNA was isolated from the heads of collected flies, converted into RNAseq libraries and sequenced, and subsequently reads were assembled to the transcriptome to quantify gene expression.

Three separate analyses were used to identify genes with significant expression changes through aging. For two of the analyses, we associated gene expression with a continuous variable, defined as either the chronological age of the flies on the day samples were collected (Day analysis), or as the survivorship of the aging cohort upon sampling (Survival analysis). In the third analysis, we associated expression with a categorical variable with 15 levels, representing the sampling points when flies were collected for RNAseq (Sampling Point analysis). This third analysis has the potential to identify genes missed by the pair of continuous variable analyses, since expression variation of some genes may not follow a simple, continuous temporal pattern.

The three analyses collectively identified 6,142 unique genes that were differentially expressed through lifespan (Figure 1), representing a little under half of the genes with detectable expression in our study (N=13,303). The analyses individually identified 5,449 (Day), 5,264 (Survival), and 4,449 (Sampling Point) genes. Around 60% of the genes (3,706) were identified by all three analyses, and an additional 28% (1,432) were identified by both continuous variable analyses, Day and Survival. The large overlap of genes identified in both the Day and Survival analyses – well over 90% of the genes identified in each analysis are shared among the two – is expected since age in days and survivorship are strongly correlated (*r* = 0.98, *p* < 10^−9^; Supplementary Figure 2). The Sampling Point analysis identified the most genes unique to one analysis – 567 (9% of the total number of unique genes identified) – demonstrating its utility in capturing genes not easily found by either continuous variable analysis.

**Figure 1:**
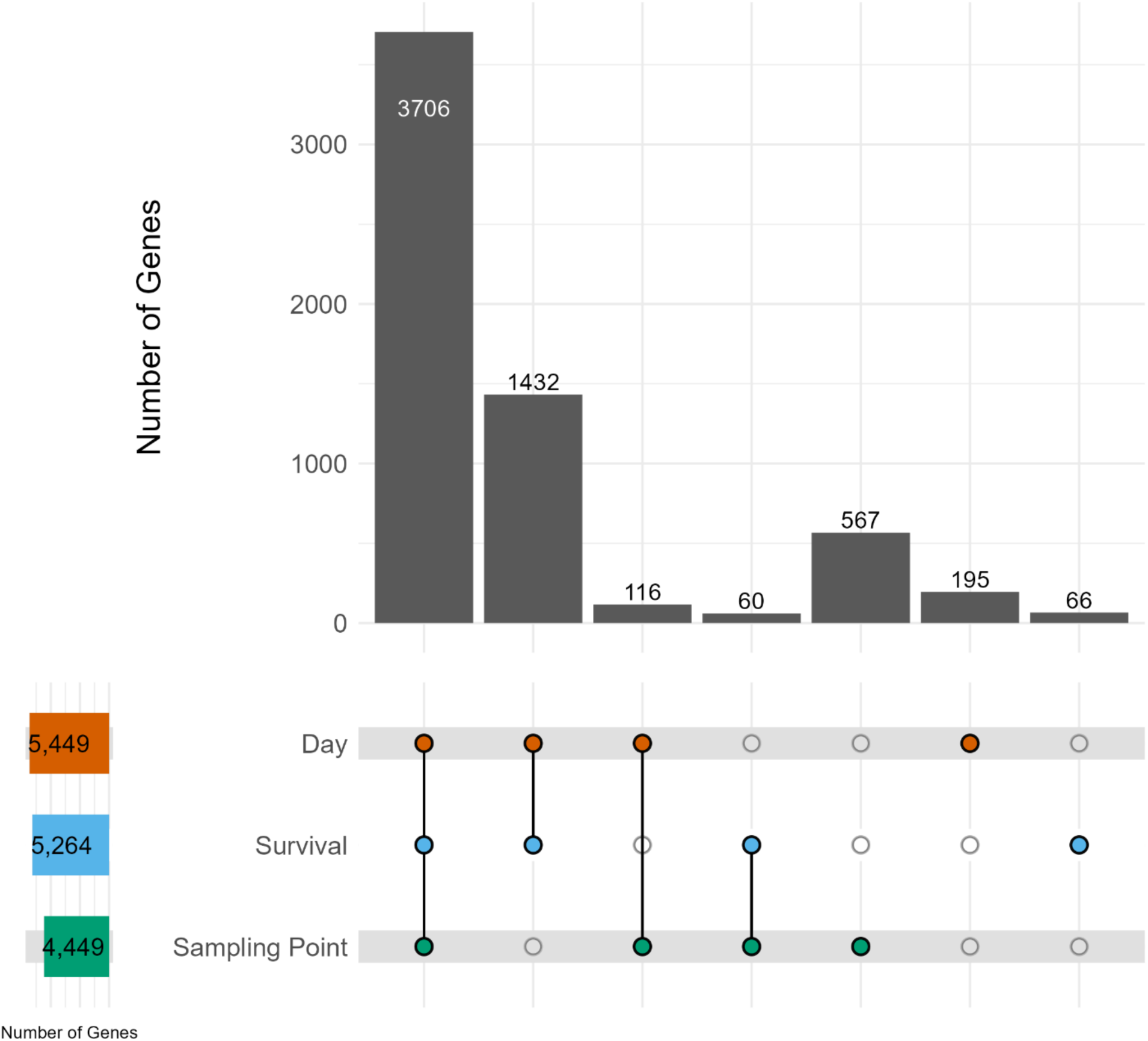
Identification of 6,142 genes with age-related gene expression. We identified genes whose expression was significantly associated with Day of life (N=5,449), with Survival (N=5,264), and with Sampling point (N=4,449). The upper bar chart shows the number of significant genes identified, with colored circles below showing which analysis the genes were identified in (3706 genes were identified in all three analyses, 1432 genes were identified in both the Day and Survival analyses, and so on).

### A diversity of gene expression trajectories through aging

Genes with similar functions, or that function in the same pathways, might be expected to share similar expression trends over time (e.g. Eisen et al. 1998; Zhang et al. 2004). To enable investigation of this, we normalized the expression trajectories of all 6,142 identified genes via *z*-scores and clustered them into 28 groups (Figure 2, Supplementary Figure 4). Based on the results of linear regressions of cluster-specific expression patterns (Figure 3) against time in Days, each cluster was assigned a trajectory designation of LinearUp or LinearDown (the mean cluster expression is significantly associated with time, and either goes up or down over time, respectively), or Complex (there is no significant association with time after correcting for multiple testing). See “Materials and Methods” for further details on this process and Supplementary Table E for statistical information. Over 80% of genes fall into the 6 LinearDown (2,596) and 8 LinearUp (2,536) clusters, with the remaining 1,010 genes split among 14 Complex clusters (Figure 3). In general, the Complex clusters harbor considerably fewer genes than the more linear clusters.

**Figure 2:**
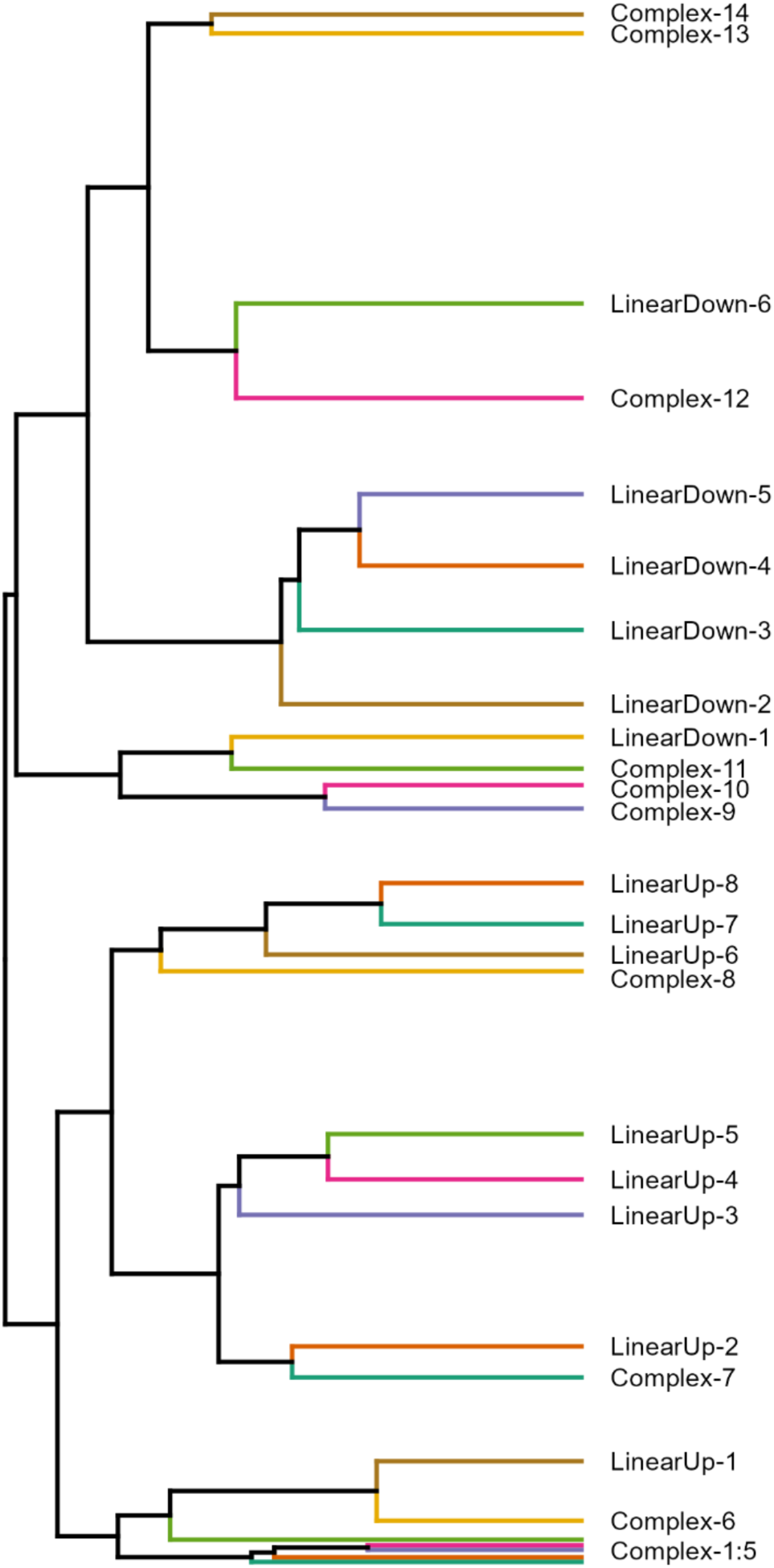
Clustering all 6,142 age-related genes into 28 clusters via their expression trajectories through lifespan. A simplified dendrogram representing the hierarchical clustering of our gene expression trajectory data. Each horizontal colored line represents all those genes in each of our 28 clusters; The closer clusters are to each other on the plot, the more similar their expression trajectories (see Figure 3). Each cluster is named based on their expression trajectory (LinearUp, LinearDown, Complex). Complex-1 to Complex-5 clusters are not individually labeled since they are very close together in the plot. Supplementary Figure 4 shows the full dendrogram highlighting the relationships among all 6,142 genes.

Clearly the LinearDown and LinearUp clusters do not show perfect linear patterns of expression; For instance, genes in LinearDown-1 (Figure 3) show a slight increase in expression in mid-life, before decreasing in expression towards late life. However, the curves for the linear clusters typically appear more linear than those of the Complex clusters (compare LinearUp-8 to Complex-1), and primarily show monotonic increases/decreases in expression over time (see LinearUp-7). Furthermore, among the Complex class of clusters we see a great deal of variation in expression trajectory; Many are decidedly non-linear, and show wave-like patterns (e.g., Complex-5), or are curved with the highest expression in mid-life (e.g., Complex-10). However, some Complex cluster are somewhat linear and exhibit patterns not dissimilar to those of LinearUp/Down clusters (e.g., compare Complex-6 with LinearUp-4). We recognize that our “linear” clusters are not perfectly linear, and do vary in their trajectories over time, and that our “complex” clusters exhibit a wide spectrum of trajectories. However, to simplify presentation, we elected to employ a straightforward trajectory-based naming scheme for the clusters we identify (i.e., LinearUp, LinearDown, Complex).

**Figure 3:**
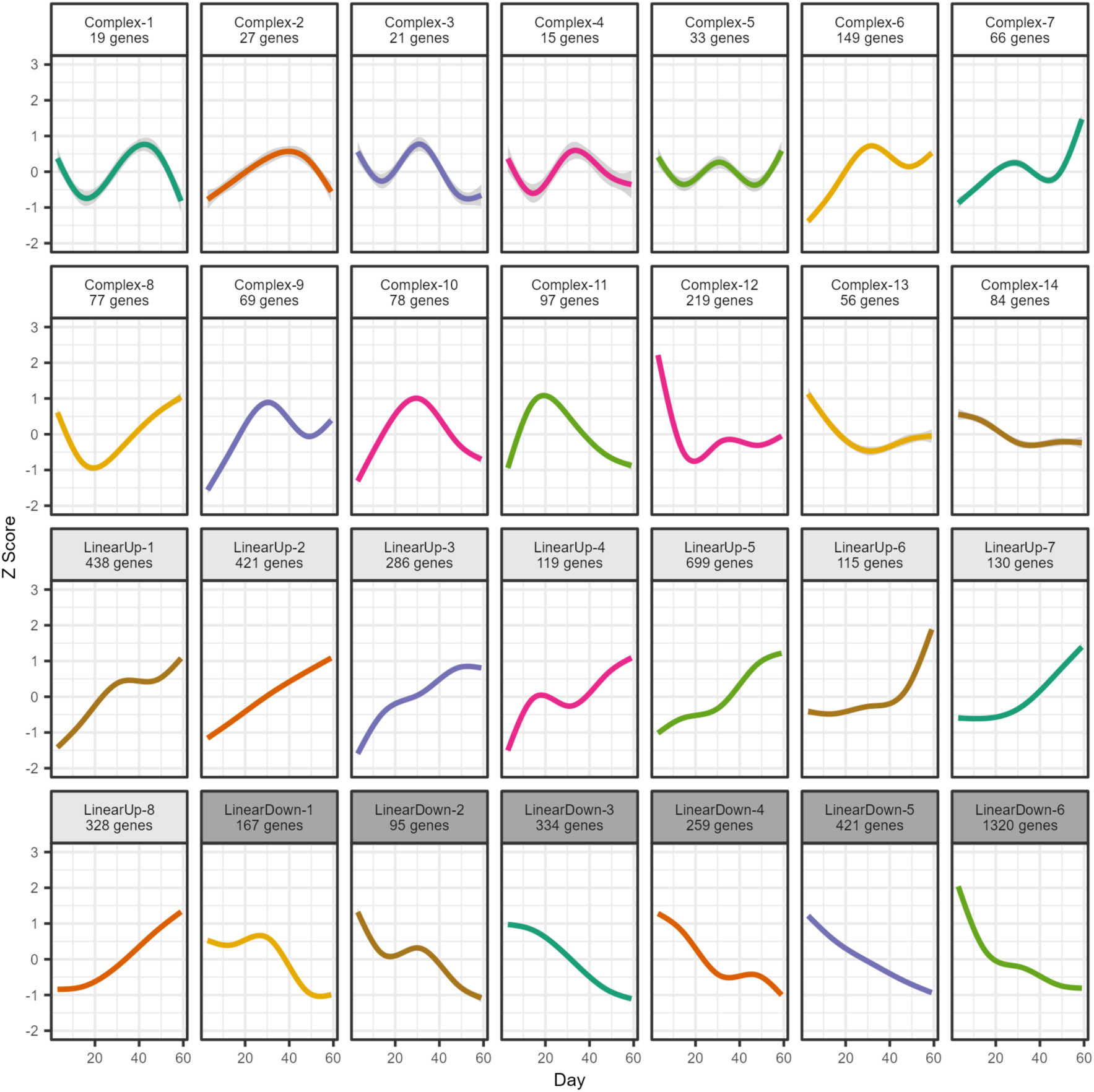
Representative expression trajectories for all 28 clusters. Within a cluster we calculated the average *z*-score over genes for each timepoint, and present a smoothed curve through those points highlighting the cluster-specific changes in gene expression over time. We determined whether each cluster-specific set of mean *z*-scores was statistically associated with age, and used this information to designate each cluster as LinearUp or LinearDown (we found a significant association, and expression either increases or decreases over time), or as Complex (there was no significant association between expression and age). This led to 14 Complex, 8 LinearUp, and 6 LinearDown clusters.

### Genes with complex expression trajectories are often identified via the sampling point analysis

We examined the relationship between the statistical analyses a gene was identified in (Day, Survival, Sampling Point), the cluster in which it resides, and the trajectory it was assigned (LinearUp, LinearDown, Complex). More than 75% of the genes identified solely in the continuous variable analyses (i.e., Day only, Survival only, or both) reside in the LinearUp/Down clusters, whereas ∼43% of those genes found in Complex clusters were uniquely identified in the Sampling Point analysis (Supplementary Figure 5). This result does not appear to be driven by specific clusters (Supplementary Figure 6). This again implies that the Sampling Point analysis has the potential to identify genes whose age-related changes in expression are difficult to capture with linear analyses based on chronological time or survivorship.

### Identification of known aging-relevant genes

Our analyses identified many genes previously associated with aging in *Drosophila*. We identified 107/176 genes associated with the Gene Ontology (GO) term “determination of adult lifespan” (GO:0008340), with at least one such gene being present in 24 of the 28 clusters. For instance, we identified *I’m not dead yet* (Complex-6, FBgn0036816), a transporter of Krebs cycle intermediates, which when mutated increases lifespan via a mechanism resembling the effect of caloric restriction (Rogina et al. 2000). We found insulin signaling genes, including *chico* (LinearUp-8, FBgn0024248) and *Insulin-like peptide 2* (Complex-11, FBgn0036046), for which loss-of-function mutations increase lifespan (Clancy et al. 2001; Grönke et al. 2010). We also identified the heat shock proteins *Hsp22* (LinearUp-6, FBgn0001223), *Hsp26* (LinearUp-8, FBgn0001225), and *Hsp68* (LinearUp-5, FBgn0001230) which when overexpressed can increase lifespan (Morrow et al. 2004; H.-D. Wang, Kazemi-Esfarjani, and Benzer 2004; M. C. Wang, Bohmann, and Jasper 2003).

### Exploring the biological functions of expression trajectory clusters

To understand if genes with similar expression trajectories share similar functions, we executed a series of enrichment analyses using the software PANGEA (Y. Hu et al. 2023), identifying enriched GO terms, gene groups and pathways within each of our 28 clusters. GO terms derive from the *Drosophila* GO subsets available within PANGEA (terms are indicated below by the “GO” stem), the gene groups are from DRSC GLAD (“GLAD”) and FlyBase (“FBgg”), and pathways come from Reactome information (“R-DME”). In total we identified 732 unique enriched terms, 595 of which are specific to just one of our expression trajectory designations (i.e., LinearUp, LinearDown, Complex), suggesting our three designations represent largely distinct sets of biological processes. Furthermore, most of the unique enrichment terms are found in just a single cluster, with only 67 terms being shared by multiple clusters. (The PANGEA output tables are available in Supplementary Tables F-K).

#### Confirming common aging-related gene expression patterns

In *Drosophila* aging expression studies it is commonly observed that both general stress response genes and immunity genes increase in expression with age (Bordet et al. 2021; Carlson et al. 2015; Girardot et al. 2006; Highfill et al. 2017; Landis et al. 2004; Pletcher et al. 2002; Zane et al. 2023), and we recapitulate this finding. Five of our clusters are enriched for genes that respond to stress (GO:0006950), all of which are designated LinearUp (LinearUp-2, 3, 5, 7, 8; Figure 3). Previous studies have seen heat shock proteins increase in expression with age (V. King and Tower 1999; Landis et al. 2004; Manière et al. 2014), and we see these genes (FBgg0000501) enriched in two LinearUp clusters (LinearUp-5, 7). Of the 5 clusters that are enriched for stress response genes, 3 are more specifically enriched for immune response genes (GO: 0006955; LinearUp-2, 5, 8). LinearUp-5 and 8 are enriched for antimicrobial peptides (FBgg0001101), including *Attacin-A* (LinearUp-8, FBgn0012042), *Listericin* (LinearUp-8, FBgn0033593), and *Drosocin* (LinearUp-5, FBgn0013088) which are commonly found to increase in expression with age in *Drosophila* expression studies (Bajgiran et al. 2021; Bordet et al. 2021; Carnes et al. 2015; Highfill et al. 2017; Lai et al. 2007; Landis et al. 2004; Zane et al. 2023).

A decrease in cognitive ability with advanced age has been reported in *D. melanogaster* (Tamura et al. 2003; Haddadi et al. 2014; Pacifico et al. 2018), mice (Lamberty and Gower 1990; Kubanis, Gobbel, and Zornetzer 1981; Healy et al. 2024), and rats (Rowe et al. 1998; Sagheddu et al. 2024). Similarly, age-related neurodegeneration is commonly observed, with decreased neurogenesis with aging being reported in both mice (Maslov et al. 2004) and rats (Kuhn, Dickinson-Anson, and Gage 1996), and synaptic deterioration seen in the motor neurons of aged *Caenorhabditis elegans* (J. Liu et al. 2013). Three of our clusters – LinearDown-3, 4 and 6 – are enriched for genes that are found in synapses (GO:0045202), are involved in synapse organization (GO:0050808), as well as those associated with cognition (GO:0050890) and nervous system development (GO:0007399). LinearDown-4 and 6 are also enriched for genes encoding voltage-gated potassium and sodium channel subunits (FBgg0000506, FBgg0000595). Notably, LinearDown-6 harbors *Atpα* (FBgn0002921), a gene encoding a subunit of the NA+/K+ exchanging ATPase pump, and which has been shown to affect lifespan (Palladino et al. 2003).

A decrease in the expression of genes involved with the electron transport chain (ETC) is a common finding in samples of aged individuals (Bordet et al. 2021; Girardot et al. 2006; Highfill et al. 2017; Landis et al. 2004; Pletcher et al. 2002; Zane et al. 2023). Such a decrease could lead to reduced ATP production, disruption of the NAD+/NADH ratio, and cellular senescence (Lenaz et al. 1997; Miwa et al. 2014; 2022). Our study supports a decrease in ETC-related gene expression; Genes in Complex-12 and LinearDown-6 show an overall decrease in expression across lifespan (Figure 3), and these clusters are adjacent to each other in the dendrogram resulting from hierarchical clustering (Figure 2 and Supplementary Figure 4). These two clusters are enriched for genes encoding mitochondrial ETC complex I and V (FBgg001836, FBgg0001849). LinearDown-6 is also enriched for genes that are part of mitochondrial complex III and IV (FBgg0001850, FBgg0001847). A series of ETC-related genes shown to influence lifespan are also present in these clusters; These include *ND-20* (Complex-12, FBgn0030718) and *ND-SGDH* (LinearDown-6, FBgn0011455) (Copeland et al. 2009), *levy* (LinearDown-6, FBgn0034877) (W. Liu et al. 2007), and *ATPsynD* (LinearDown-6, FBgn0016120) (Sun et al. 2014). Additionally, LinearDown-6 harbors *stress-sensitive B* (FBgn0003360), which is involved in the transport of ADP and ATP in and out of the mitochondrial matrix, and when mutated shortens lifespan (Celotto et al. 2006; Reynolds 2018).

#### A distinction between cytosolic and mitochondrial ribosomal gene expression responses

A decrease in the expression of ribosomal proteins and ribosomal biogenesis genes with age has been documented in a number of yeast studies (Choi et al. 2018; Janssens et al. 2015; Kamei et al. 2014; Philipp et al. 2013; Yiu et al. 2008), with three of these showing a reduction in cytosolic ribosome (GO: 0022626) gene expression over time (Choi et al. 2018; Philipp et al. 2013; Yiu et al. 2008). One fly study we identified (Doroszuk et al. 2012) observed a similar response, with enrichment of ribosome-related ontology terms in genes showing a reduction in expression with age in a given strain, including genes involved in ribosome biogenesis (GO:0042254) and mitochondrial ribosome (GO:0005761).

In the analyses of our dataset, we saw enrichment of ribosomal-related ontology terms in six clusters. Five of these clusters show a general increase in gene expression with age (LinearUp-1, 2, 4, 6, Complex-6; see Figure 3), and include genes associated with the terms ribosome biogenesis (GO:0042254), rRNA processing (R-DME-72312), cytoplasmic ribosomal proteins (FBgg0000141), and structural constituent of ribosomes (GO:0003735). The sixth (LinearDown-6) shows reduced expression with age, and shows enrichment of mitochondrial ribosomal proteins (FBgg0000059). Thus, in an apparent contrast with prior results, it appears that in our data many ribosome-associated genes increase in expression with age, while only mitochondrial ribosomal protein genes decrease with advanced age.

To examine this further we extracted from our complete set of 6,142 differentially-expressed genes all those affiliated with any ribosome-related term (N=270, See Supplementary Table L for list of terms). We plotted their age-related expression, separating out mitochondrial ribosomal proteins (N=36, FBgg0000059), and can clearly see that while all mitochondrial ribosomal protein genes go down in expression with age, nearly all other ribosomal genes (220/234) increase in expression with age (Supplementary Figure 8). Furthermore, just contrasting cytoplasmic (N=80, FBgg0000141) and mitochondrial (N=36, FBgg0000059) ribosomal proteins, we see the former all go up, and the latter all go down in expression with age (Supplementary Figure 7).

A *D. melanogaster* brain-specific, single-cell RNAseq study (Davie et al. 2018) appears to support our finding that – outside of mitochondrial ribosomes – ribosomal genes increase in expression with age. First, Davie et al. report that the cytosolic small ribosomal subunit (GO: 0022627) gene *stubarista* (FBgn0003517) is statistically upregulated during aging; We also identified this gene in our LinearUp-5 cluster. Second, while Davie et al. see a reduction in the overall levels of RNA and transcription through aging – a result seen previously (Tahoe, Mokhtarzadeh, and Curtsinger 2004), and which we also observed (Supplementary Table B and Supplementary Figure 9) – on average, ribosomal protein genes show a lower decline in expression than other genes (see Fig. 5C in Davie et al. 2018). In a bulk RNAseq analysis framework this result would be expected to translate to a relative increase in the expression of ribosomal proteins over time. Nonetheless, that we see a result that contrasts with some of the prior work on gene expression changes through aging is intriguing and worthy of future examination.

#### Various metabolic processes are enriched in some Complex clusters

Many of our Complex clusters have small gene counts, and relatively few enriched terms. However, 8/14 Complex clusters (2, 3, 4, 6, 7, 9, 10 and 12) are enriched for genes associated with metabolism (R-DME-1430728 and GLAD:24593). While this term is broad, when we focus on individual clusters more specific metabolic functions are evident. As described earlier, Complex-12 is enriched for genes involved with the ETC. Complex-2, 6, and 10 are specifically enriched for genes involved in the pentose phosphate pathway (R-DME-71336), a glucose catabolism pathway that produces NADPH and ribose sugars for nucleotide synthesis (Stincone et al. 2015). Notably, Complex-2 harbors *Pgd* (FBgn0004654) and *G6pd* (FBgn0004057) which are the two reducing enzymes involved in the pentose phosphate pathway (Gvozdev et al. 1976; Geer, Bowman, and Simmons 1974). Complex-3 is uniquely enriched for genes involved in galactose catabolism (R-DME-70370) and glycogen synthesis (R-DME-3322077), one of which – *Agbe* (FBgn0053138) – is involved in lifespan (Paik et al. 2012). That all of these metabolic pathways/genes emerged from our Complex clusters suggests that nonlinear changes in metabolic activity occur throughout lifespan, as has been suggested in a large, multi-omic study in humans (Shen et al. 2024).

#### Enrichment for protein folding, modification, and transport in the LinearUp-7 cluster

LinearUp-7 – which shows only limited change in expression for the first third of life, followed by increasing expression until end of life – is enriched for multiple terms involved with proper protein folding and modification, and transporting proteins from the endoplasmic reticulum to the golgi apparatus. We see enrichment for chaperones and co-chaperones (FBgg0001643), and heatshock proteins (see above), which can bind onto unfolded proteins (GO:0051082) and help correctly fold them (GO:0006457). Additionally, LinearUp-7 is enriched for genes involved in post-translation protein modification pathways (R-DME-597592). LinearUp-7 is the only cluster to be enriched for genes associated with both the endoplasmic reticulum (GO:0005783) and the golgi apparatus (GO:0005794). It includes genes that are involved in transport between these two organelles, and is enriched for both coat protein complex I (FBgg0000087) and II (FBgg0000116) genes, and genes involved in ER-to-Golgi anterograde transport (R-DME-199977), and Golgi-to-ER retrograde transport (R-DME-8856688). These enrichment patterns further demonstrate that genes with similar roles can have very similar temporal expression patterns.

### Multi-timepoint trajectory-based datasets offer more detail than two-timepoint studies

Often aging expression studies compare “young” and “old” samples, and we sought to re-analyze our data in this framework to discover what is gained from a multi-timepoint approach. Our “young” timepoint combined all the Day 3 and Day 6 samples (to achieve a sample size of 6), and our “old” timepoint used the six Day 59 samples. Contrasting these two sets of samples yielded 4,533 differentially-expressed genes. Of the 6,142 genes identified in our multi-timepoint analysis, 4,347 (∼71%) were re-identified in this two-timepoint analysis. The reduction in the number of genes is likely a combination of the switch in analytical approach, and a simple loss of power (since we have gone from 48 to 12 samples). Considering the assigned expression trajectories from our multi-timepoint analysis, the two-timepoint analysis recovered 77.5% of the LinearDown cluster genes, 72% of the LinearUp genes, but only 50.5% of the Complex genes (Figure 4). As anticipated, genes that do not exhibit a straightforward, monotonic increase/decrease in expression through lifespan are much less likely to be identified when sampling is restricted to very young and very old animals.

**Figure 4:**
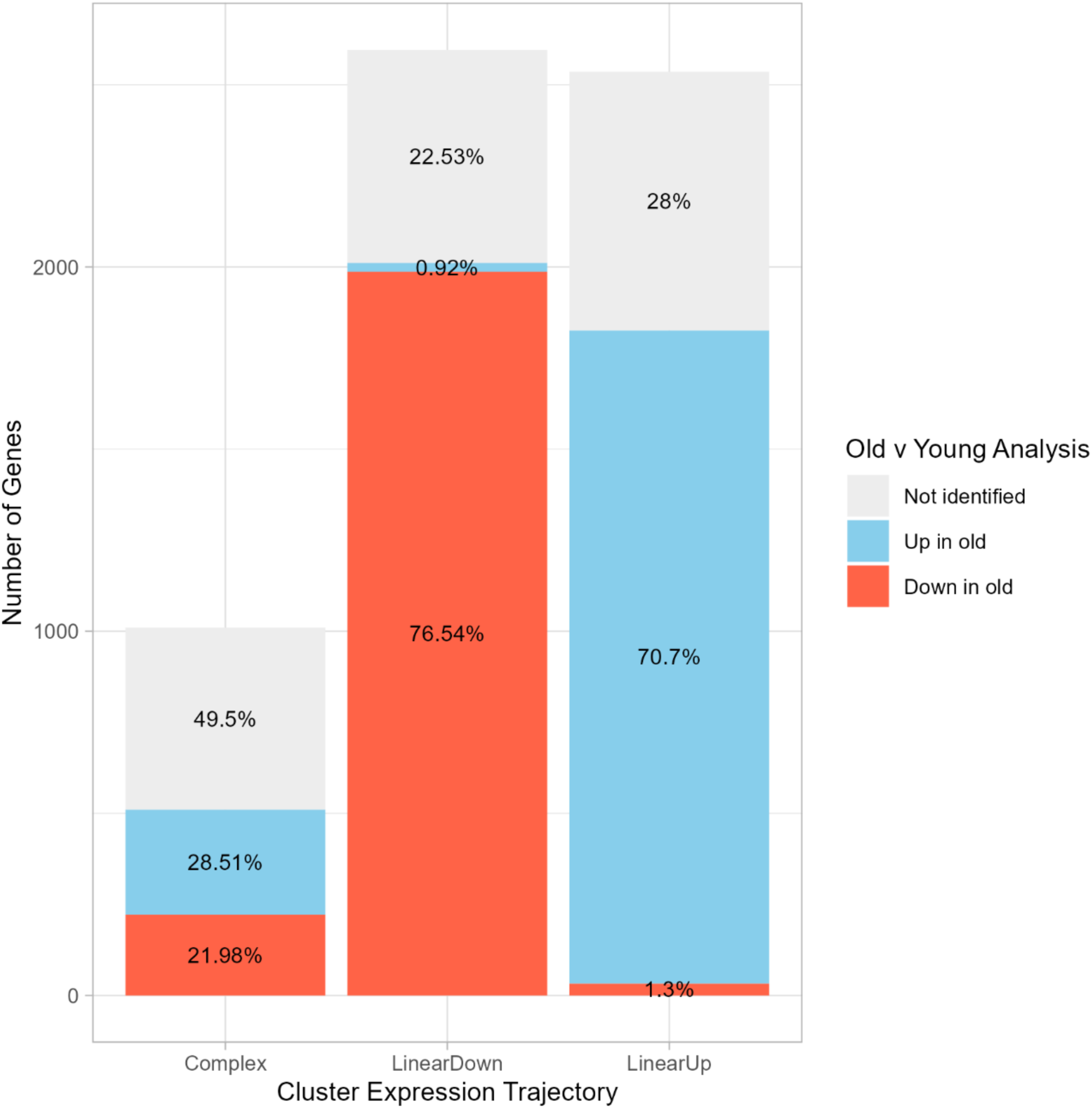
Differences between a multi-timepoint and a two-timepoint analysis: Our 3 trajectory-based analyses revealed > 6,000 differentially-expressed genes that were grouped into 3 trajectories (Complex, LinearDown, LinearUp). We compared these results to a re-analysis of a subset of the same data, where we directly contrasted expression between young (Day 3+6) and old (Day 59) samples. Each vertical bar depicts the fraction (in the figure) and the number (*y*-axis) of genes in each of our expression trajectories that are absent in the young versus old test (gray), are significantly up-regulated in old animals (blue) or are significantly down-regulated in old animals (red). Most genes with linear trajectories are re-identified, and ∼99% of those show the expected direction of change with age. However, only ∼50% of the Complex trajectory genes are re-identified in the two-timepoint analysis, and these are split between those that appear to increase or to decrease in expression with age.

A challenge with a two-timepoint analysis is that the only conclusion one can draw about the expression change is that it goes up or down with age. For those genes present in LinearUp/LinearDown clusters that were replicated in the two-timepoint analysis, the inferred direction of the expression change matched expectations 98-99% of the time (Figure 4), and this trend is consistent across the different LinearUp/LinearDown clusters (90%-100% matched) (Supplementary Figure 10). However, for the 50.5% of Complex genes that were re-identified in the two-timepoint analysis, 56% of them showed an increase in expression in old samples, and 44% showed a decrease (Figure 4). Examining the two-timepoint expression change direction calls across Complex clusters reveals cluster-to-cluster variation (see Supplementary Figure 10), and in most cases, re-identified genes within a given Complex cluster exhibit the same direction of expression change in the two-timepoint analysis (see Supplementary Figure 10). This likely reflects the particular, cluster-specific nature of each Complex expression trajectory, and what the precise expression levels were at the start and end of our experiment. For instance, Complex-7 shows a general increase in expression over time (Figure 3), and all genes re-identified in the two-timepoint analysis are marked as increasing in expression (Supplementary Figure 10).

In our initial analysis we chose Days 3+6 and Day 59 to represent our “young” and “old” samples. However, studies have used flies of quite different ages to represent young and old animals (Bajgiran et al. 2021; Bordet et al. 2021; Carnes et al. 2015; Girardot et al. 2006; Landis et al. 2004). To understand the impact of varying the “old” timepoint, we repeated the two-timepoint analysis several times. In each case, we fixed the young timepoint using our Days 3+6 data, and derived the old timepoint from two sequential sampling days (e.g., Days 10+14, Days 14+17, Days 17+23, and so on) such that each analysis compared two sets of 6 samples. We then compared the differentially-expressed genes that emerged from these analyses, along with the inferred expression change, with our initial Day 3+6 versus Day 59 analysis. As might be expected, using old sampling points that occur earlier in life results in fewer significant differentially expressed genes (Supplementary Figure 11); there has simply been less time for change. As the old timepoint moves later in life there is increasingly greater overlap with the Day 3+6 versus Day 59 analysis, and 97-99% of the shared genes show the same change in expression (Supplementary Figure 11). Nonetheless, every alternative analysis with a different old timepoint identified genes we initially identified in our multi-timepoint trajectory analysis, but that were missed in our initial two-timepoint analysis (Supplementary Figure 11). Particularly in those analyses using old timepoints that occur earlier in life (up to Day 36, a little over halfway through the lifespan of our cohort), such genes are often associated with our multi-timepoint Complex trajectory clusters (Supplementary Figure 12). Between 9-17% of Complex genes found using the earlier alternative old timepoints, were not found in our initial Day3+6 versus Day 59 test. It is likely that the observed change in the expression of these genes over time is highly dependent on the exact timepoints chosen. In general, our analyses clearly demonstrate that the age when individuals are sampled impacts the genes identified.

### Comparison with other aging expression studies in flies

We compared the outcome of our multi-timepoint analysis with 9 datasets from 8 previously published expression analyses in *D. melanogaster* (Bajgiran et al. 2021; Bordet et al. 2021; Carnes et al. 2015; Girardot et al. 2006; Highfill, Reeves, and Macdonald 2016; Lai et al. 2007; Landis et al. 2004; Zane et al. 2023). These were all two-timepoint studies that varied in the sex of the flies, the target tissue, and the sampling points employed (Supplementary Table M). Around 95% of the genes resulting from our multi-timepoint analysis were identified in at least 1 of these datasets, and the number of previous studies that identified a given gene was not clearly associated with the trajectory designation (LinearUp, LinearDown, Complex) we assigned (Supplementary Figure 13). The high rate of gene re-identification across studies is notable given the diversity of the study designs in terms of fly sex, the tissue targeted, the sampling points used (Supplementary Table M), and the likely many differences in the precise rearing/maintenance conditions employed. That the same sets of genes are regularly identified suggests some similarity across genotype/sex/tissue in the age-related expression profile (Izgi et al. 2022).

Similar to the comparisons between multi- and two-timepoint analyses within our own dataset (above), when we look at genes in our LinearUp/LinearDown clusters that were identified in prior studies, they broadly show the direction of expression change we would predict (Supplementary Figure 14). However, as might also be expected, the fraction of such genes showing the predicted expression change in a prior study is often lower than it is in our within-study methodological comparison (compare Figure 4 with Supplementary Figure 14). The highest fraction of LinearUp/LinearDown genes showing the expected expression change in a prior study – 95-98% – is with Highfill et al. (2016; Supplementary Figure 14), a study published by our group that used the same tissue type, employed a very similar fly maintenance environment, but targeted females rather than males. The other prior studies we examined varied more in their design, and these differences likely contribute to shared genes more often showing mis-matched expression changes (see Supplementary Figure 14).

### Benefits of a multi-timepoint, trajectory-based transcriptomics approach to exploring dynamic biological processes

It is increasingly clear that a complete view of the gene regulatory changes underlying a range of dynamic processes – from the response to infection (Schlamp et al. 2021) to cellular differentiation (Strober et al. 2019) to aging (Pletcher et al. 2002; Shen et al. 2024; Gheorghe et al. 2014) – requires a timecourse experimental design, interrogating samples taken throughout the process of interest. Here, we showed that a two-timepoint, young versus old analysis can successfully identify, and correctly infer the direction of expression change of many genes that have largely linear expression trajectories through aging. But we also showed that the set of genes identified depends on the pair of timepoints chosen. Furthermore, we showed that by limiting the sampling to just two timepoints, we would have failed to identify almost half of the genes with more complex, non-linear expression trajectories, and would not have captured the nature of the trajectories for genes that were identified. Even for those genes in the clusters we consider to be “linear” there is variation in rate of expression change with time, which can only be captured with a multi-timepoint approach (Gheorghe et al. 2014; Haustead et al. 2016; Lu et al. 2004; Schaum et al. 2020; Shen et al. 2024).

Another major benefit of being able to consider the trajectory of gene expression is that it enables the identification of genes with similar longitudinal expression patterns, and allows assessment of whether genes with similar patterns have similar functional/molecular properties. Our enrichment analyses support the contention that genes with similar expression trajectories share similar functions. Due to the multiple points we sampled throughout the aging process, around 70% of all significant, enriched gene ontology terms we identified were unique to a single cluster. This specificity allowed us to more precisely characterize the expression patterns of various sets of genes with related functional roles, rather than only – with a two-timepoint framework – being able to state that particular groups of genes are up- or down-regulated with age. Better understanding of the dynamic molecular changes underling aging will facilitate a deeper understanding of the cellular and physiological changes that occur as organisms age.

### Caveats

We recognize that the scope of our results may be somewhat limited due to the use of only males from one inbred strain, and the use of a single tissue type. Additionally, while our target tissue – the fly head – is enriched for brain/neuronal cells, since it also includes a mixture of other cells types, we lack true tissue specificity. This said, we were able to demonstrate some consistency over studies in the genes and expression patterns identified, despite these studies varying in multiple ways, making our results a useful resource for future aging investigations.

### Accessible data exploration via interactive shiny apps

Our expression, gene enrichment, and comparative analyses generated a significant amount of data. Above we have only discussed a subset of our observations. To make the results more accessible – in addition to sharing our raw data, summary data, and analytical code – we have developed two interactive R/shiny apps that enable individuals to explore our gene identification, clustering, and cluster enrichment results. The apps allow users to look up specific genes, examine their expression trajectory through time in our cohort of flies, the cluster they belong to, as well as to explore enriched terms within and across clusters. See the Materials and Methods, along with Supplementary Texts C and D for more information on the development of the apps, and how to access and use them.

## Supporting information

All Supplementary Materials

## ACKNOWLEDGEMENTS

We thank the KU Genome Sequencing Core (supported by NIH P30 GM145499) for library construction and sequencing, and the Kansas INBRE Data Science Core (supported by NIH P20 GM103418) for computational infrastructure. This work was supported by NIH R21 AG086734 and by NIH R01 OD034064.

## REFERENCES

Alic, Nazif, Maria E. Giannakou, Irene Papatheodorou, Matthew P. Hoddinott, T. Daniel Andrews, Ekin Bolukbasi, and Linda Partridge. 2014. “Interplay of DFOXO and Two ETS-Family Transcription Factors Determines Lifespan in Drosophila Melanogaster.” PLOS Genetics 10 (9): e1004619. 10.1371/journal.pgen.1004619.

Bajgiran, Morteza, Azali Azlan, Shaharum Shamsuddin, Ghows Azzam, and Mardani Abdul Halim. 2021. “Data on RNA-Seq Analysis of Drosophila Melanogaster during Ageing.” Data in Brief 38 (October):107413. 10.1016/j.dib.2021.107413.

Bordet, Guillaume, Niraj Lodhi, Andrew Kossenkov, and Alexei Tulin. 2021. “Age-Related Changes of Gene Expression Profiles in Drosophila.” Genes 12 (12): 1982. 10.3390/genes12121982.

Broer, Linda, Aron S. Buchman, Joris Deelen, Daniel S. Evans, Jessica D. Faul, Kathryn L. Lunetta, Paola Sebastiani, et al. 2015. “GWAS of Longevity in CHARGE Consortium Confirms APOE and FOXO3 Candidacy.” The Journals of Gerontology Series A: Biological Sciences and Medical Sciences 70 (1): 110–18. 10.1093/gerona/glu166.

Cai, Nanshuo, Yifan Wu, and Yan Huang. 2022. “Induction of Accelerated Aging in a Mouse Model.” Cells 11 (9): 1418. 10.3390/cells11091418.

Carlson, Kimberly A., Kylee Gardner, Anjeza Pashaj, Darby J. Carlson, Fang Yu, James D. Eudy, Chi Zhang, and Lawrence G. Harshman. 2015. “Genome-Wide Gene Expression in Relation to Age in Large Laboratory Cohorts of Drosophila Melanogaster.” Genetics Research International 2015:835624. 10.1155/2015/835624.

Carnes, Megan Ulmer, Terry Campbell, Wen Huang, Daniel G. Butler, Mary Anna Carbone, Laura H. Duncan, Sasha V. Harbajan, et al. 2015. “The Genomic Basis of Postponed Senescence in Drosophila Melanogaster.” PLoS ONE 10 (9): e0138569. 10.1371/journal.pone.0138569.

Celotto, Alicia M., Adam C. Frank, Steven W. McGrath, Tim Fergestad, Wayne A. Van Voorhies, Karolyn F. Buttle, Carmen A. Mannella, and Michael J. Palladino. 2006. “Mitochondrial Encephalomyopathy in Drosophila.” The Journal of Neuroscience: The OCicial Journal of the Society for Neuroscience 26 (3): 810–20. 10.1523/JNEUROSCI.4162-05.2006.

Chakraborty, Mahul, Nicholas W. VanKuren, Roy Zhao, Xinwen Zhang, Shannon Kalsow, and J. J. Emerson. 2018. “Hidden Genetic Variation Shapes the Structure of Functional Elements in Drosophila.” Nature Genetics 50 (1): 20–25. 10.1038/s41588-017-0010-y.

Chang, Winston, Joe Cheng, JJ Allaire, Carson Sievert, Barret Schloerke, Yihui Xie, Jeff Allen, Jonathan McPherson, Alan Dipert, and Barbara Borges. 2023. “Shiny: Web Application Framework for R.” R. https://CRAN.R-project.org/package=shiny.

Choi, Kyung-Mi, Seok-Jin Hong, Jan M van Deursen, Sooah Kim, Kyoung Heon Kim, and Cheol-Koo Lee. 2018. “Caloric Restriction and Rapamycin Differentially Alter Energy Metabolism in Yeast.” The Journals of Gerontology: Series A 73 (1): 29–38. 10.1093/gerona/glx024.

Clancy, D. J., D. Gems, L. G. Harshman, S. Oldham, H. Stocker, E. Hafen, S. J. Leevers, and L. Partridge. 2001. “Extension of Life-Span by Loss of CHICO, a Drosophila Insulin Receptor Substrate Protein.” *Science (New York*, N.Y.) 292 (5514): 104–6. 10.1126/science.1057991.

Copeland, Jeffrey M., Jaehyoung Cho, Thomas Lo, Jae H. Hur, Sepehr Bahadorani, Tagui Arabyan, Jason Rabie, Jennifer Soh, and David W. Walker. 2009. “Extension of Drosophila Life Span by RNAi of the Mitochondrial Respiratory Chain.” Current Biology: CB 19 (19): 1591–98. 10.1016/j.cub.2009.08.016.

Cypser, James R., David Kitzenberg, and Sang-Kyu Park. 2013. “Dietary Restriction in C. Elegans: Recent Advances.” Experimental Gerontology 48 (10): 1014–17. 10.1016/j.exger.2013.02.018.

Davie, Kristofer, Jasper Janssens, Duygu Koldere, Maxime De Waegeneer, Uli Pech, Łukasz Kreft, Sara Aibar, et al. 2018. “A Single-Cell Transcriptome Atlas of the Aging Drosophila Brain.” Cell 174 (4): 982–998.e20. 10.1016/j.cell.2018.05.057.

Deelen, Joris, Marian Beekman, Hae-Won Uh, Linda Broer, Kristin L. Ayers, Qihua Tan, Yoichiro Kamatani, et al. 2014. “Genome-Wide Association Meta-Analysis of Human Longevity Identifies a Novel Locus Conferring Survival beyond 90 Years of Age.” Human Molecular Genetics 23 (16): 4420–32. 10.1093/hmg/ddu139.

Deelen, Joris, Marian Beekman, Hae-Won Uh, Quinta Helmer, Maris Kuningas, Lene Christiansen, Dennis Kremer, et al. 2011. “Genome-Wide Association Study Identifies a Single Major Locus Contributing to Survival into Old Age; the APOE Locus Revisited.” Aging Cell 10 (4): 686–98. 10.1111/j.1474-9726.2011.00705.x.

Doroszuk, Agnieszka, Martijs J. Jonker, Nicolien Pul, Timo M. Breit, and Bas J. Zwaan. 2012. “Transcriptome Analysis of a Long-Lived Natural Drosophila Variant: A Prominent Role of Stress- and Reproduction-Genes in Lifespan Extension.” BMC Genomics 13 (1): 167. 10.1186/1471-2164-13-167.

Eisen, M. B., P. T. Spellman, P. O. Brown, and D. Botstein. 1998. “Cluster Analysis and Display of Genome-Wide Expression Patterns.” Proceedings of the National Academy of Sciences of the United States of America 95 (25): 14863–68. 10.1073/pnas.95.25.14863.

Frenk, Stephen, and Jonathan Houseley. 2018. “Gene Expression Hallmarks of Cellular Ageing.” Biogerontology 19 (6): 547–66. 10.1007/s10522-018-9750-z.

Geer, B. W., J. T. Bowman, and J. R. Simmons. 1974. “The Pentose Shunt in Wild-Type and Glucose-6-Phosphate Dehydrogenase Deficient Drosophila Melanogaster.” The Journal of Experimental Zoology 187 (1): 77–86. 10.1002/jez.1401870110.

Gheorghe, Marius, Marc Snoeck, Michael Emmerich, Thomas Bäck, Jelle J. Goeman, and Vered Raz. 2014. “Major Aging-Associated RNA Expressions Change at Two Distinct Age-Positions.” BMC Genomics 15 (1): 132. 10.1186/1471-2164-15-132.

Giannakou, Maria E., Martin Goss, Jake Jacobson, Giovanna Vinti, Sally J. Leevers, and Linda Partridge. 2007. “Dynamics of the Action of DFOXO on Adult Mortality in Drosophila.” Aging Cell 6 (4): 429–38. 10.1111/j.1474-9726.2007.00290.x.

Girardot, Fabrice, Christelle Lasbleiz, Véronique Monnier, and Hervé Tricoire. 2006. “Specific Age Related Signatures in Drosophila Body Parts Transcriptome.” BMC Genomics 7 (1): 69. 10.1186/1471-2164-7-69.

Gocmez, Semil Selcen, Yusufhan Yazir, Gulcin Gacar, Tuğçe Demirtaş Şahin, Sertan Arkan, Ayşe Karson, and Tijen Utkan. 2020. “Etanercept Improves Aging-Induced Cognitive Deficits by Reducing Inflammation and Vascular Dysfunction in Rats.” Physiology & Behavior 224 (October):113019. 10.1016/j.physbeh.2020.113019.

Grönke, Sebastian, David-Francis Clarke, Susan Broughton, T. Daniel Andrews, and Linda Partridge. 2010. “Molecular Evolution and Functional Characterization of Drosophila Insulin-like Peptides.” PLoS Genetics 6 (2): e1000857. 10.1371/journal.pgen.1000857.

Gvozdev, V. A., T. I. Gerasimova, G. L. Kogan, and null Braslavskaya OYu. 1976. “Role of the Pentose Phosphate Pathway in Metabolism of Drosophila Melanogaster Elucidated by Mutations Affecting Glucose 6-Phosphate and 6-Phosphogluconate Dehydrogenases.” FEBS Letters 64 (1): 85–88. 10.1016/0014-5793(76)80255-4.

Haddadi, Mohammad, Samaneh Reiszadeh Jahromi, B. K. Chandrasekhar Sagar, Rajashekhar K. Patil, T. Shivanandappa, and S. R. Ramesh. 2014. “Brain Aging, Memory Impairment and Oxidative Stress: A Study in Drosophila Melanogaster.” Behavioural Brain Research 259 (February):60–69. 10.1016/j.bbr.2013.10.036.

Harrell Jr, Frank E. 2023. “Rms: Regression Modeling Strategies.” R. https://CRAN.R-project.org/package=rms.

Haustead, Daniel J., Andrew Stevenson, Vishal Saxena, Fiona Marriage, Martin Firth, Robyn Silla, Lisa Martin, et al. 2016. “Transcriptome Analysis of Human Ageing in Male Skin Shows Mid-Life Period of Variability and Central Role of NF-ΚB.” Scientific Reports 6 (1): 26846. 10.1038/srep26846.

Healy, Dáire, Carol Murray, Ciara McAdams, Ruth Power, Pierre-Louis Hollier, Jessica Lambe, Lucas Tortorelli, Ana Belen Lopez-Rodriguez, and Colm Cunningham. 2024. “Susceptibility to Acute Cognitive Dysfunction in Aged Mice Is Underpinned by Reduced White Matter Integrity and Microgliosis.” Communications Biology 7 (1): 1–15. 10.1038/s42003-023-05662-9.

Herskind, A. M., M. McGue, N. V. Holm, T. I. Sørensen, B. Harvald, and J. W. Vaupel. 1996. “The Heritability of Human Longevity: A Population-Based Study of 2872 Danish Twin Pairs Born 1870-1900.” Human Genetics 97 (3): 319–23. 10.1007/BF02185763.

Highfill, Chad A., G. Adam Reeves, and Stuart J. Macdonald. 2016. “Genetic Analysis of Variation in Lifespan Using a Multiparental Advanced Intercross Drosophila Mapping Population.” BMC Genetics 17 (August):113. 10.1186/s12863-016-0419-9.

Highfill, Chad A., Jonathan H. Tran, Samantha K. T. Nguyen, Taylor R. Moldenhauer, Xiaofei Wang, and Stuart J. Macdonald. 2017. “Naturally Segregating Variation at Ugt86Dd Contributes to Nicotine Resistance in Drosophila Melanogaster.” Genetics 207 (1): 311–25. 10.1534/genetics.117.300058.

Hu, Frank B. 2024. “Diet Strategies for Promoting Healthy Aging and Longevity: An Epidemiological Perspective.” Journal of Internal Medicine 295 (4): 508–31. 10.1111/joim.13728.

Hu, Yanhui, Aram Comjean, Helen Attrill, Giulia Antonazzo, Jim Thurmond, Weihang Chen, Fangge Li, et al. 2023. “PANGEA: A New Gene Set Enrichment Tool for Drosophila and Common Research Organisms.” Nucleic Acids Research 51 (W1): W419–26. 10.1093/nar/gkad331.

Hu, Zheng, Kaifu Chen, Zheng Xia, Myrriah Chavez, Sangita Pal, Ja-Hwan Seol, Chin-Chuan Chen, Wei Li, and Jessica K. Tyler. 2014. “Nucleosome Loss Leads to Global Transcriptional Up-Regulation and Genomic Instability during Yeast Aging.” Genes & Development 28 (4): 396–408. 10.1101/gad.233221.113.

Izgi, Hamit, Dingding Han, Ulas Isildak, Shuyun Huang, Ece Kocabiyik, Philipp Khaitovich, Mehmet Somel, and Handan Melike Dönertaş. 2022. “Inter-Tissue Convergence of Gene Expression during Ageing Suggests Age-Related Loss of Tissue and Cellular Identity.” ELife 11 (January):e68048. 10.7554/eLife.68048.

Janssens, Georges E, Anne C Meinema, Javier González, Justina C Wolters, Alexander Schmidt, Victor Guryev, Rainer Bischoff, Ernst C Wit, Liesbeth M Veenhoff, and Matthias Heinemann. 2015. “Protein Biogenesis Machinery Is a Driver of Replicative Aging in Yeast.” Edited by Karsten Weis. ELife 4 (September):e08527. 10.7554/eLife.08527.

Joshi, Peter K., Krista Fischer, Katharina E. Schraut, Harry Campbell, Tõnu Esko, and James F. Wilson. 2016. “Variants near CHRNA3/5 and APOE Have Age- and Sex-Related Effects on Human Lifespan.” Nature Communications 7 (1): 11174. 10.1038/ncomms11174.

Joshi, Peter K., Nicola Pirastu, Katherine A. Kentistou, Krista Fischer, Edith Hofer, Katharina E. Schraut, David W. Clark, et al. 2017. “Genome-Wide Meta-Analysis Associates HLA-DQA1/DRB1 and LPA and Lifestyle Factors with Human Longevity.” Nature Communications 8 (1): 910. 10.1038/s41467-017-00934-5.

Kamei, Yuka, Yoshihiro Tamada, Yasumune Nakayama, Eiichiro Fukusaki, and Yukio Mukai. 2014. “Changes in Transcription and Metabolism During the Early Stage of Replicative Cellular Senescence in Budding Yeast*.” Journal of Biological Chemistry 289 (46): 32081–93. 10.1074/jbc.M114.600528.

Kaplanis, Joanna, Assaf Gordon, Tal Shor, Omer Weissbrod, Dan Geiger, Mary Wahl, Michael Gershovits, et al. 2018. “Quantitative Analysis of Population-Scale Family Trees with Millions of Relatives.” *Science (New York*, N.Y.) 360 (6385): 171–75. 10.1126/science.aam9309.

Kenyon, Cynthia J. 2010. “The Genetics of Ageing.” Nature 464 (7288): 504–12. 10.1038/nature08980.

Kimura, Koutarou D., Heidi A. Tissenbaum, Yanxia Liu, and Gary Ruvkun. 1997. “Daf-2, an Insulin Receptor-Like Gene That Regulates Longevity and Diapause in Caenorhabditis Elegans.” Science 277 (5328): 942–46. 10.1126/science.277.5328.942.

King, Elizabeth G., Chris M. Merkes, Casey L. McNeil, Steven R. Hoofer, Saunak Sen, Karl W. Broman, Anthony D. Long, and Stuart J. Macdonald. 2012. “Genetic Dissection of a Model Complex Trait Using the Drosophila Synthetic Population Resource.” Genome Research 22 (8): 1558–66. 10.1101/gr.134031.111.

King, V., and J. Tower. 1999. “Aging-Specific Expression of Drosophila Hsp22.” Developmental Biology 207 (1): 107–18. 10.1006/dbio.1998.9147.

Kraus, William E., Manjushri Bhapkar, Kim M. Huffman, Carl F. Pieper, Sai Krupa Das, Leanne M. Redman, Dennis T. Villareal, et al. 2019. “2 Years of Calorie Restriction and Cardiometabolic Risk (CALERIE): Exploratory Outcomes of a Multicentre, Phase 2, Randomised Controlled Trial.” The Lancet. Diabetes & Endocrinology 7 (9): 673–83. 10.1016/S2213-8587(19)30151-2.

Kubanis, P., G. Gobbel, and S. F. Zornetzer. 1981. “Age-Related Memory Deficits in Swiss Mice.” Behavioral and Neural Biology 32 (2): 241–47. 10.1016/S0163-1047(81)90555-0.

Kuhn, H. G., H. Dickinson-Anson, and F. H. Gage. 1996. “Neurogenesis in the Dentate Gyrus of the Adult Rat: Age-Related Decrease of Neuronal Progenitor Proliferation.” The Journal of Neuroscience: The OCicial Journal of the Society for Neuroscience 16 (6): 2027–33. 10.1523/JNEUROSCI.16-06-02027.1996.

Lai, Chao-Qiang, Laurence D. Parnell, Richard F. Lyman, Jose M. Ordovas, and Trudy F. C. Mackay. 2007. “Candidate Genes Affecting Drosophila Life Span Identified by Integrating Microarray Gene Expression Analysis and QTL Mapping.” Mechanisms of Ageing and Development 128 (3): 237–49. 10.1016/j.mad.2006.12.003.

Lamberty, Y., and A. J. Gower. 1990. “Age-Related Changes in Spontaneous Behavior and Learning in NMRI Mice from Maturity to Middle Age.” Physiology & Behavior 47 (6): 1137–44. 10.1016/0031-9384(90)90364-a.

Landis, Gary N., Diana Abdueva, Dmitriy Skvortsov, Junsheng Yang, Beth E. Rabin, James Carrick, Simon Tavaré, and John Tower. 2004. “Similar Gene Expression Patterns Characterize Aging and Oxidative Stress in Drosophila Melanogaster.” Proceedings of the National Academy of Sciences of the United States of America 101 (20): 7663–68. 10.1073/pnas.0307605101.

Lenaz, G., C. Bovina, C. Castelluccio, R. Fato, G. Formiggini, M. L. Genova, M. Marchetti, et al. 1997. “Mitochondrial Complex I Defects in Aging.” Molecular and Cellular Biochemistry 174 (1–2): 329–33.

Lin, K., J. B. Dorman, A. Rodan, and C. Kenyon. 1997. “Daf-16: An HNF-3/Forkhead Family Member That Can Function to Double the Life-Span of Caenorhabditis Elegans.” *Science (New York*, N.Y.) 278 (5341): 1319–22. 10.1126/science.278.5341.1319.

Liu, Jie, Bi Zhang, Haoyun Lei, Zhaoyang Feng, Jianfeng Liu, Ao-Lin Hsu, and X. Z. Shawn Xu. 2013. “Functional Aging in the Nervous System Contributes to Age-Dependent Motor Activity Decline in C. Elegans.” Cell Metabolism 18 (3): 392–402. 10.1016/j.cmet.2013.08.007.

Liu, Wensheng, Radhakrishnan Gnanasambandam, Jeffery Benjamin, Gunisha Kaur, Patricia B. Getman, Alan J. Siegel, Randall D. Shortridge, and Satpal Singh. 2007. “Mutations in Cytochrome c Oxidase Subunit VIa Cause Neurodegeneration and Motor Dysfunction in Drosophila.” Genetics 176 (2): 937–46. 10.1534/genetics.107.071688.

Lu, Tao, Ying Pan, Shyan-Yuan Kao, Cheng Li, Isaac Kohane, Jennifer Chan, and Bruce A. Yankner. 2004. “Gene Regulation and DNA Damage in the Ageing Human Brain.” Nature 429 (6994): 883–91. 10.1038/nature02661.

Lund, James, Patricia Tedesco, Kyle Duke, John Wang, Stuart K Kim, and Thomas E Johnson. 2002. “Transcriptional Profile of Aging in *C. Elegans*.” Current Biology 12 (18): 1566–73. 10.1016/S0960-9822(02)01146-6.

Manière, X., A. Krisko, F. X. Pellay, J.-M. Di Meglio, P. Hersen, and I. Matic. 2014. “High Transcript Levels of Heat-Shock Genes Are Associated with Shorter Lifespan of Caenorhabditis Elegans.” Experimental Gerontology 60 (December):12–17. 10.1016/j.exger.2014.09.005.

Maslov, Alexander Y., Tara A. Barone, Robert J. Plunkett, and Steven C. Pruitt. 2004. “Neural Stem Cell Detection, Characterization, and Age-Related Changes in the Subventricular Zone of Mice.” The Journal of Neuroscience: The OCicial Journal of the Society for Neuroscience 24 (7): 1726–33. 10.1523/JNEUROSCI.4608-03.2004.

Mattison, Julie A., Ricki J. Colman, T. Mark Beasley, David B. Allison, Joseph W. Kemnitz, George S. Roth, Donald K. Ingram, Richard Weindruch, Rafael de Cabo, and Rozalyn M. Anderson. 2017. “Caloric Restriction Improves Health and Survival of Rhesus Monkeys.” Nature Communications 8 (January):14063. 10.1038/ncomms14063.

Mayer, Peter J. 1991. “Inheritance of Longevity Evinces No Secular Trend among Members of Six New England Families Born 1650-1874.” American Journal of Human Biology: The OCicial Journal of the Human Biology Council 3 (1): 49–58. 10.1002/ajhb.1310030109.

McCarroll, Steven A., Coleen T. Murphy, Sige Zou, Scott D. Pletcher, Chen-Shan Chin, Yuh Nung Jan, Cynthia Kenyon, Cornelia I. Bargmann, and Hao Li. 2004. “Comparing Genomic Expression Patterns across Species Identifies Shared Transcriptional Profile in Aging.” Nature Genetics 36 (2): 197–204. 10.1038/ng1291.

McCay, C. M., L. A. Maynard, G. Sperling, and LeRoy L. Barnes. 1939. “Retarded Growth, Life Span, Ultimate Body Size and Age Changes in the Albino Rat after Feeding Diets Restricted in Calories: Four Figures.” The Journal of Nutrition 18 (1): 1–13. 10.1093/jn/18.1.1.

Melzer, David, Luke C. Pilling, and Luigi Ferrucci. 2020. “The Genetics of Human Ageing.” Nature Reviews. Genetics 21 (2): 88–101. 10.1038/s41576-019-0183-6.

Miwa, Satomi, Howsun Jow, Karen Baty, Amy Johnson, Rafal Czapiewski, Gabriele Saretzki, Achim Treumann, and Thomas von Zglinicki. 2014. “Low Abundance of the Matrix Arm of Complex I in Mitochondria Predicts Longevity in Mice.” Nature Communications 5 (1): 3837. 10.1038/ncomms4837.

Miwa, Satomi, Sonu Kashyap, Eduardo Chini, and Thomas von Zglinicki. 2022. “Mitochondrial Dysfunction in Cell Senescence and Aging.” The Journal of Clinical Investigation 132 (13): e158447. 10.1172/JCI158447.

Morrow, Geneviève, Sophie Battistini, Ping Zhang, and Robert M. Tanguay. 2004. “Decreased Lifespan in the Absence of Expression of the Mitochondrial Small Heat Shock Protein Hsp22 in Drosophila.” The Journal of Biological Chemistry 279 (42): 43382–85. 10.1074/jbc.C400357200.

Newman, Anne B., and Joanne M. Murabito. 2013. “The Epidemiology of Longevity and Exceptional Survival.” Epidemiologic Reviews 35 (1): 181–97. 10.1093/epirev/mxs013.

Newman, Anne B., Stefan Walter, Kathryn L. Lunetta, Melissa E. Garcia, P. Eline Slagboom, Kaare Christensen, Alice M. Arnold, et al. 2010. “A Meta-Analysis of Four Genome-Wide Association Studies of Survival to Age 90 Years or Older: The Cohorts for Heart and Aging Research in Genomic Epidemiology Consortium.” The Journals of Gerontology: Series A 65A (5): 478–87. 10.1093/gerona/glq028.

Olecka, Maja, Alena van Bömmel, Lena Best, Madlen Haase, Silke Foerste, Konstantin Riege, Thomas Dost, et al. 2024. “Nonlinear DNA Methylation Trajectories in Aging Male Mice.” Nature Communications 15 (1): 3074. 10.1038/s41467-024-47316-2.

Pacifico, Rodrigo, Courtney M. MacMullen, Erica Walkinshaw, Xiaofan Zhang, and Ronald L. Davis. 2018. “Brain Transcriptome Changes in the Aging Drosophila Melanogaster Accompany Olfactory Memory Performance Deficits.” PLoS ONE 13 (12): e0209405. 10.1371/journal.pone.0209405.

Paik, Donggi, Yeo Gil Jang, Young Eun Lee, Young Nam Lee, Rochelle Yamamoto, Heon Yung Gee, Seungmin Yoo, et al. 2012. “Misexpression Screen Delineates Novel Genes Controlling Drosophila Lifespan.” Mechanisms of Ageing and Development 133 (5): 234–45. 10.1016/j.mad.2012.02.001.

Palladino, Michael J., Jill E. Bower, Robert Kreber, and Barry Ganetzky. 2003. “Neural Dysfunction and Neurodegeneration in Drosophila Na+/K+ ATPase Alpha Subunit Mutants.” The Journal of Neuroscience: The OCicial Journal of the Society for Neuroscience 23 (4): 1276–86. 10.1523/JNEUROSCI.23-04-01276.2003.

Philipp, Oliver, Andrea Hamann, Jörg Servos, Alexandra Werner, Ina Koch, and Heinz D. Osiewacz. 2013. “A Genome-Wide Longitudinal Transcriptome Analysis of the Aging Model Podospora Anserine.” PLOS ONE 8 (12): e83109. 10.1371/journal.pone.0083109.

Pilling, Luke C., Janice L. Atkins, Kirsty Bowman, Samuel E. Jones, Jessica Tyrrell, Robin N. Beaumont, Katherine S. Ruth, et al. 2016. “Human Longevity Is Influenced by Many Genetic Variants: Evidence from 75,000 UK Biobank Participants.” Aging 8 (3): 547–60. 10.18632/aging.100930.

Pilling, Luke C., Chia-Ling Kuo, Kamil Sicinski, Jone Tamosauskaite, George A. Kuchel, Lorna W. Harries, Pamela Herd, Robert Wallace, Luigi Ferrucci, and David Melzer. 2017. “Human Longevity: 25 Genetic Loci Associated in 389,166 UK Biobank Participants.” Aging 9 (12): 2504–20. 10.18632/aging.101334.

Pletcher, Scott D., Stuart J. Macdonald, Richard Marguerie, Ulrich Certa, Stephen C. Stearns, David B. Goldstein, and Linda Partridge. 2002. “Genome-Wide Transcript Profiles in Aging and Calorically Restricted Drosophila Melanogaster.” Current Biology: CB 12 (9): 712–23. 10.1016/s0960-9822(02)00808-4.

Remondini, Daniel, Stefano Salvioli, Mirko Francesconi, Michela Pierini, Dawn J. Mazzatti, Jonathan R. Powell, Isabella Zironi, Ferdinando Bersani, Gastone Castellani, and Claudio Franceschi. 2010. “Complex Patterns of Gene Expression in Human T Cells during in Vivo Aging.” Molecular BioSystems 6 (10): 1983–92. 10.1039/C004635C.

Reynolds, Elaine R. 2018. “Shortened Lifespan and Other Age-Related Defects in Bang Sensitive Mutants of Drosophila Melanogaster.” G3: Genes|Genomes|Genetics 8 (12): 3953. 10.1534/g3.118.200610.

Rogina, B., R. A. Reenan, S. P. Nilsen, and S. L. Helfand. 2000. “Extended Life-Span Conferred by Cotransporter Gene Mutations in Drosophila.” *Science (New York*, N.Y.) 290 (5499): 2137–40. 10.1126/science.290.5499.2137.

Rowe, W. B., E. Spreekmeester, M. J. Meaney, R. Quirion, and J. Rochford. 1998. “Reactivity to Novelty in Cognitively-Impaired and Cognitively-Unimpaired Aged Rats and Young Rats.” Neuroscience 83 (3): 669–80. 10.1016/s0306-4522(97)00464-8.

Ruby, J. Graham, Kevin M. Wright, Kristin A. Rand, Amir Kermany, Keith Noto, Don Curtis, Neal Varner, et al. 2018. “Estimates of the Heritability of Human Longevity Are Substantially Inflated Due to Assortative Mating.” Genetics 210 (3): 1109–24. 10.1534/genetics.118.301613.

Sagheddu, Claudia, Tamara Stojanovic, Shima Kouhnavardi, Artem Savchenko, Ahmed M. Hussein, Marco Pistis, Francisco J. Monje, et al. 2024. “Cognitive Performance in Aged Rats Is Associated with Differences in Distinctive Neuronal Populations in the Ventral Tegmental Area and Altered Synaptic Plasticity in the Hippocampus.” Frontiers in Aging Neuroscience 16 (February). 10.3389/fnagi.2024.1357347.

Schaum, Nicholas, Benoit Lehallier, Oliver Hahn, Róbert Pálovics, Shayan Hosseinzadeh, Song E. Lee, Rene Sit, et al. 2020. “Aging Hallmarks Exhibit Organ-Specific Temporal Signatures.” Nature 583 (7817): 596–602. 10.1038/s41586-020-2499-y.

Schlamp, Florencia, Sofie Y. N. Delbare, Angela M. Early, Martin T. Wells, Sumanta Basu, and Andrew G. Clark. 2021. “Dense Time-Course Gene Expression Profiling of the Drosophila Melanogaster Innate Immune Response.” BMC Genomics 22 (April):304. 10.1186/s12864-021-07593-3.

Sebastiani, Paola, Anastasia Gurinovich, Harold Bae, Stacy Andersen, Alberto Malovini, Gil Atzmon, Francesco Villa, et al. 2017. “Four Genome-Wide Association Studies Identify New Extreme Longevity Variants.” The Journals of Gerontology: Series A 72 (11): 1453–64. 10.1093/gerona/glx027.

Sebastiani, Paola, Nadia Solovieff, Andrew T. DeWan, Kyle M. Walsh, Annibale Puca, Stephen W. Hartley, Efthymia Melista, et al. 2012. “Genetic Signatures of Exceptional Longevity in Humans.” PLOS ONE 7 (1): e29848. 10.1371/journal.pone.0029848.

Shen, Xiaotao, Chuchu Wang, Xin Zhou, Wenyu Zhou, Daniel Hornburg, Si Wu, and Michael P. Snyder. 2024. “Nonlinear Dynamics of Multi-Omics Profiles during Human Aging.” *Nature Aging*, August, 1–16. 10.1038/s43587-024-00692-2.

Stincone, Anna, Alessandro Prigione, Thorsten Cramer, Mirjam M. C. Wamelink, Kate Campbell, Eric Cheung, Viridiana Olin-Sandoval, et al. 2015. “The Return of Metabolism: Biochemistry and Physiology of the Pentose Phosphate Pathway.” Biological Reviews of the Cambridge Philosophical Society 90 (3): 927–63. 10.1111/brv.12140.

Strober, B.J., R. Elorbany, K. Rhodes, N. Krishnan, K. Tayeb, A. Battle, and Y. Gilad. 2019. “Dynamic Genetic Regulation of Gene Expression during Cellular Differentiation.” *Science (New York*, N.Y.) 364 (6447): 1287–90. 10.1126/science.aaw0040.

Sun, Xiaoping, Charles T. Wheeler, Jason Yolitz, Mara Laslo, Thomas Alberico, Yaning Sun, Qisheng Song, and Sige Zou. 2014. “A Mitochondrial ATP Synthase Subunit Interacts with TOR Signaling to Modulate Protein Homeostasis and Lifespan in Drosophila.” Cell Reports 8 (6): 1781–92. 10.1016/j.celrep.2014.08.022.

Tahoe, Nuzha M. A., Ali Mokhtarzadeh, and James W. Curtsinger. 2004. “Age-Related RNA Decline in Adult Drosophila Melanogaster.” The Journals of Gerontology: Series A 59 (9): B896–901. 10.1093/gerona/59.9.B896.

Tamura, Takuya, Ann-Shyn Chiang, Naomi Ito, Hsin-Ping Liu, Junjiro Horiuchi, Tim Tully, and Minoru Saitoe. 2003. “Aging Specifically Impairs Amnesiac-Dependent Memory in Drosophila.” Neuron 40 (5): 1003–11. 10.1016/s0896-6273(03)00732-3.

Tan, Qihua, Jing Hua Zhao, Dongfeng Zhang, Torben A. Kruse, and Kaare Christensen. 2008. “Power for Genetic Association Study of Human Longevity Using the Case-Control Design.” American Journal of Epidemiology 168 (8): 890–96. 10.1093/aje/kwn205.

Taormina, Giusi, Federica Ferrante, Salvatore Vieni, Nello Grassi, Antonio Russo, and Mario G. Mirisola. 2019. “Longevity: Lesson from Model Organisms.” Genes 10 (7): 518. 10.3390/genes10070518.

Timmers, Paul Rhj, Ninon Mounier, Kristi Lall, Krista Fischer, Zheng Ning, Xiao Feng, Andrew D. Bretherick, et al. 2019. “Genomics of 1 Million Parent Lifespans Implicates Novel Pathways and Common Diseases and Distinguishes Survival Chances.” ELife 8 (January):e39856. 10.7554/eLife.39856.

Wang, Horng-Dar, Parsa Kazemi-Esfarjani, and Seymour Benzer. 2004. “Multiple-Stress Analysis for Isolation of Drosophila Longevity Genes.” Proceedings of the National Academy of Sciences of the United States of America 101 (34): 12610–15. 10.1073/pnas.0404648101.

Wang, Meng C., Dirk Bohmann, and Heinrich Jasper. 2003. “JNK Signaling Confers Tolerance to Oxidative Stress and Extends Lifespan in Drosophila.” Developmental Cell 5 (5): 811–16. 10.1016/s1534-5807(03)00323-x.

Wang, Xueqing, Quanlong Jiang, Yuanyuan Song, Zhidong He, Hongdao Zhang, Mengjiao Song, Xiaona Zhang, et al. 2022. “Ageing Induces Tissue-specific Transcriptomic Changes in Caenorhabditis Elegans.” The EMBO Journal 41 (8): e109633. 10.15252/embj.2021109633.

Wickham, Hadley. 2016. “Ggplot2: Elegant Graphics for Data Analysis.” R.

Willcox, Bradley J., Timothy A. Donlon, Qimei He, Randi Chen, John S. Grove, Katsuhiko Yano, Kamal H. Masaki, D. Craig Willcox, Beatriz Rodriguez, and J. David Curb. 2008. “FOXO3A Genotype Is Strongly Associated with Human Longevity.” Proceedings of the National Academy of Sciences 105 (37): 13987–92. 10.1073/pnas.0801030105.

Wilson, C. H., S. Shalini, A. Filipovska, T. R. Richman, S. Davies, S. D. Martin, S. L. McGee, et al. 2015. “Age-Related Proteostasis and Metabolic Alterations in Caspase-2-Deficient Mice.” Cell Death & Disease 6 (1): e1615. 10.1038/cddis.2014.567.

Xie, Kan, Helmut Fuchs, Enzo Scifo, Dan Liu, Ahmad Aziz, Juan Antonio Aguilar-Pimentel, Oana Veronica Amarie, et al. 2022. “Deep Phenotyping and Lifetime Trajectories Reveal Limited Effects of Longevity Regulators on the Aging Process in C57BL/6J Mice.” Nature Communications 13 (1): 6830. 10.1038/s41467-022-34515-y.

Yiu, Gloria, Alejandra McCord, Alison Wise, Rishi Jindal, Jennifer Hardee, Allen Kuo, Michelle Yuen Shimogawa, et al. 2008. “Pathways Change in Expression During Replicative Aging in Saccharomyces Cerevisiae.” *The Journals of Gerontology. Series A*, Biological Sciences and Medical Sciences 63 (1): 21–34.

Zane, Flaminia, Hayet Bouzid, Sofia Sosa Marmol, Mira Brazane, Savandara Besse, Julia Lisa Molina, Céline Cansell, et al. 2023. “Smurfness-based Two-phase Model of Ageing Helps Deconvolve the Ageing Transcriptional Signature.” Aging Cell 22 (11): e13946. 10.1111/acel.13946.

Zeng, Yi, Chao Nie, Junxia Min, Xiaomin Liu, Mengmeng Li, Huashuai Chen, Hanshi Xu, et al. 2016. “Novel Loci and Pathways Significantly Associated with Longevity.” Scientific Reports 6 (1): 21243. 10.1038/srep21243.

Zhang, Wen, Quaid D. Morris, Richard Chang, Ofer Shai, Malina A. Bakowski, Nicholas Mitsakakis, Naveed Mohammad, et al. 2004. “The Functional Landscape of Mouse Gene Expression.” Journal of Biology 3 (5): 21. 10.1186/jbiol16.

