## Supplementary material for "Dynamic Changes in Gene Expression Through Aging in *Drosophila melanogaster* Heads": All Supplementary Materials: supp_mat_20241210.pdf

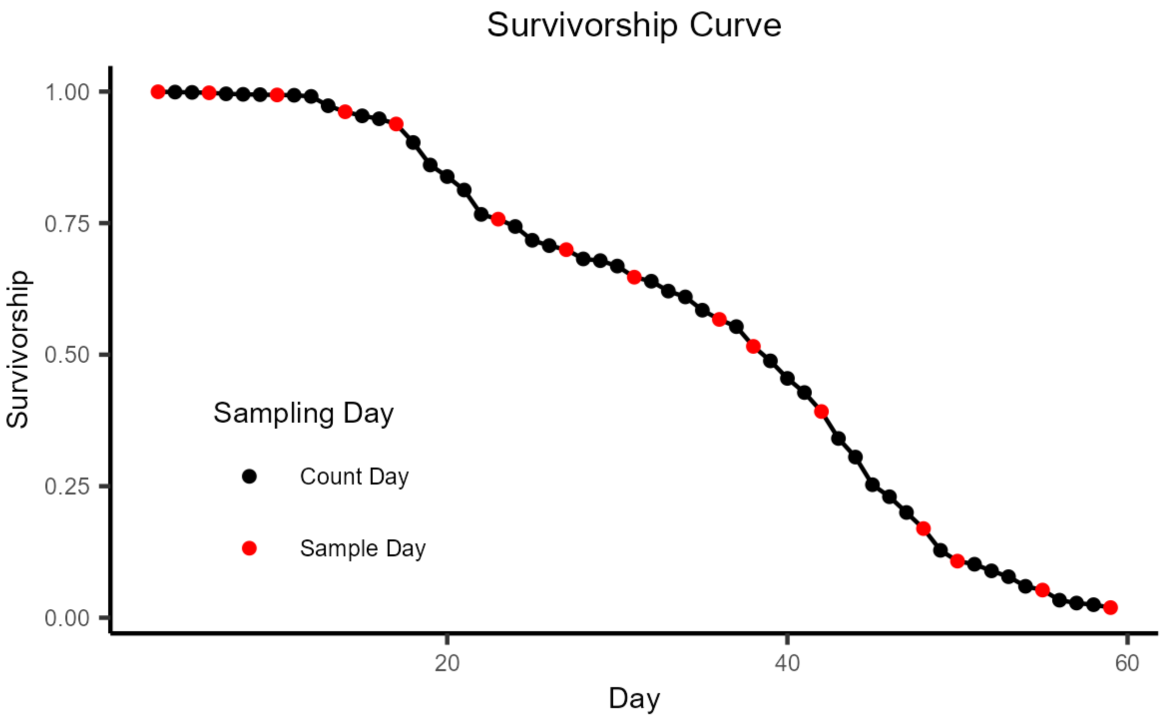


**Supplementary Figure 1:** Kaplan-Meier survivorship curve. Each point represents one day of the experiment, with black points denoting days when only the number of dead flies were counted, and red points those days when flies were also sampled. Sampling days were spaced out across the lifespan of the cohort, with an average of 4 days in between each sampling day. We began sampling on day 3 (with 99.9% of the flies remaining alive) and continued until day 59, when we sampled the final 1.92% of flies that remained alive.

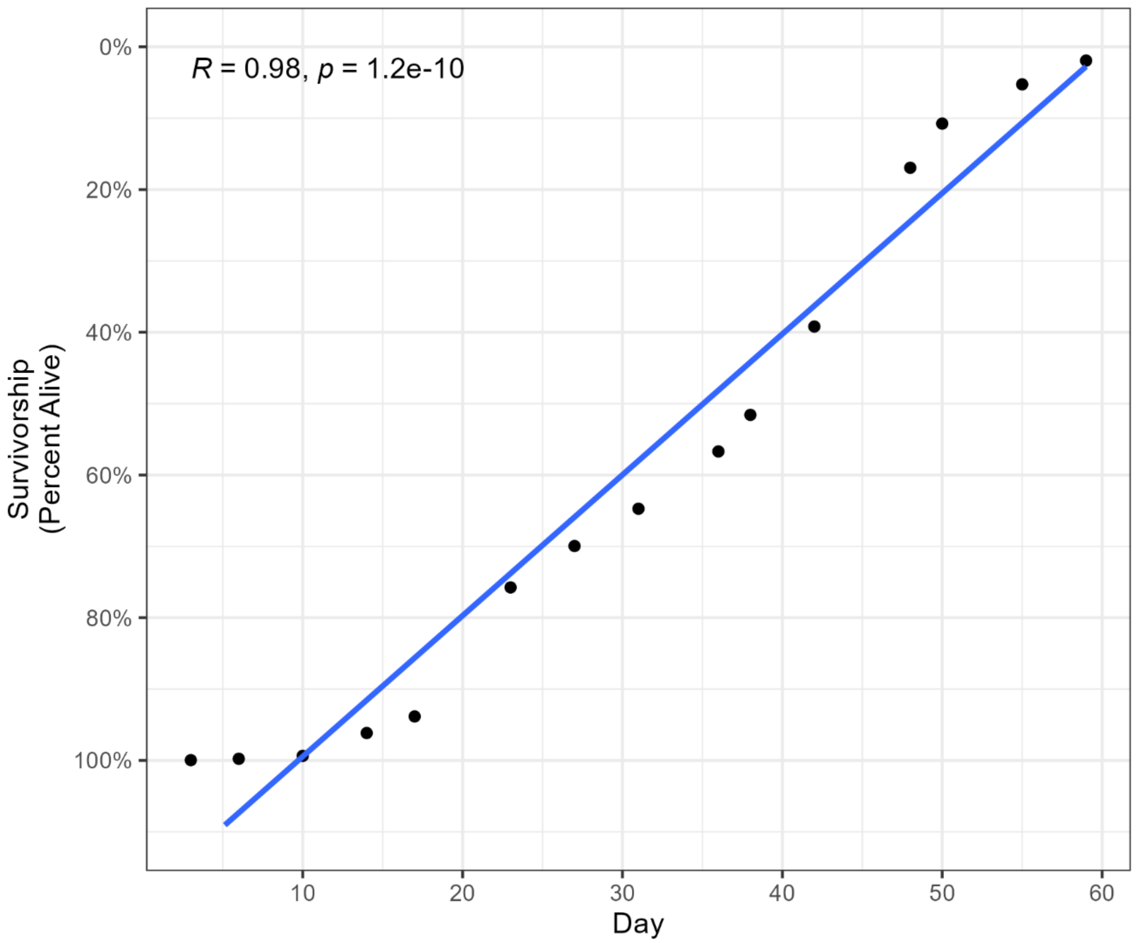


**Supplementary Figure 2:** Correlation between day of life and survivorship. The dots represent the 15 sampling points, which are plotted based on the day the sample was collected and the calculated survivorship of the fly population on that day (note that the survivorship *y*-axis is plotted in reverse). Day and survivorship have a Pearson's correlation coefficient of 0.98, which is highly significant ($p < 10^{-9}$).

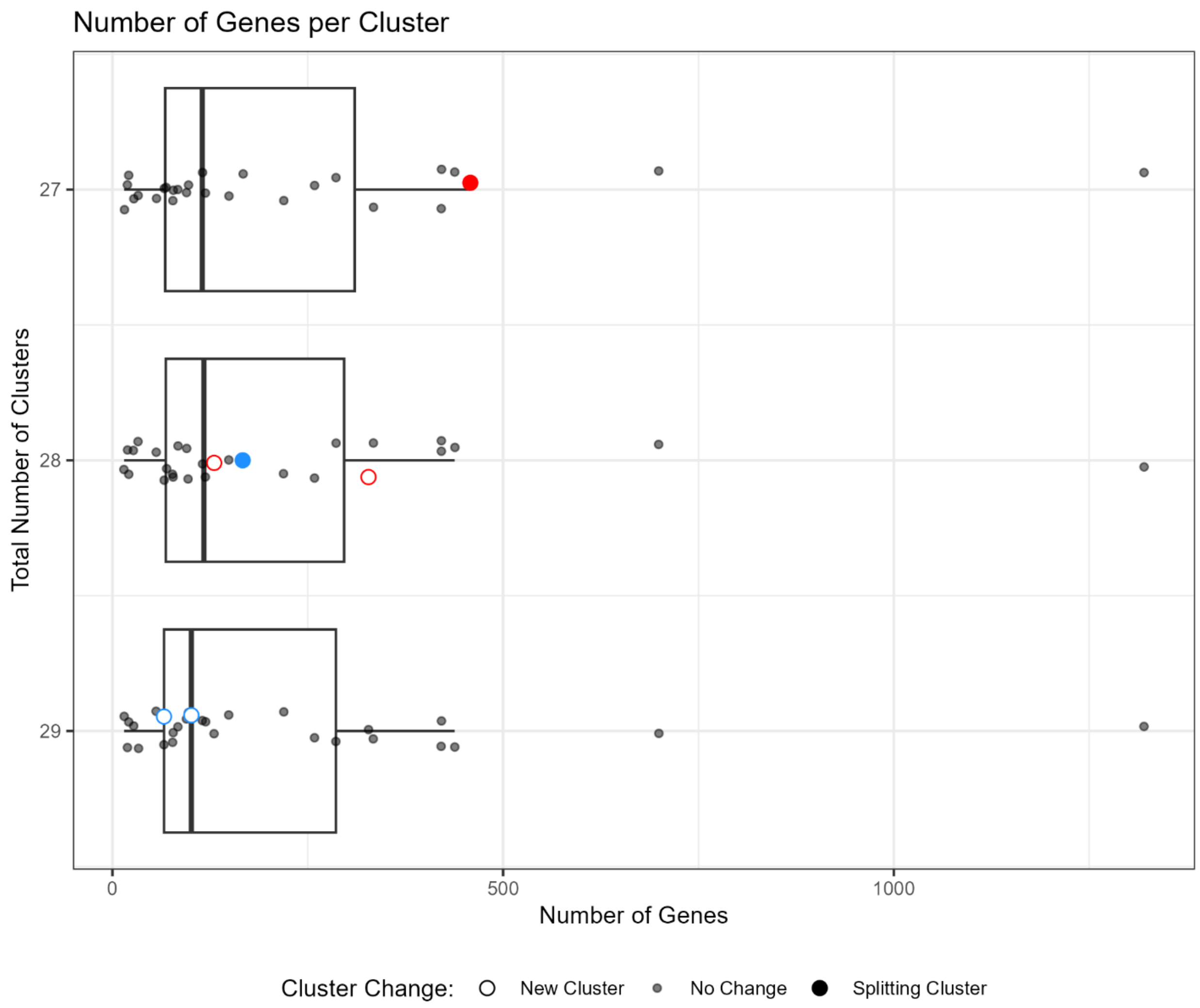


**Supplementary Figure 3:** Number of genes per cluster when varying the total number of clusters. We narrowed down our ideal cluster number to 27-29 clusters (see Supplementary Table D for further details), then for each of these values plotted the number of genes per cluster. Each point in the plot represents the gene count in 1 cluster. The solid red circle in “27” is a cluster (N=458 genes) that is split into two (open red circles) in “28”. Similarly, the solid blue circle in “28” is a cluster (N=167 genes) that splits into two (blue open circles) in “29”. Given that moving from 27 to 28 clusters splits the third largest cluster, but moving to 29 retains the two largest clusters with gene counts >500, we elected to divide our gene set into 28 clusters.

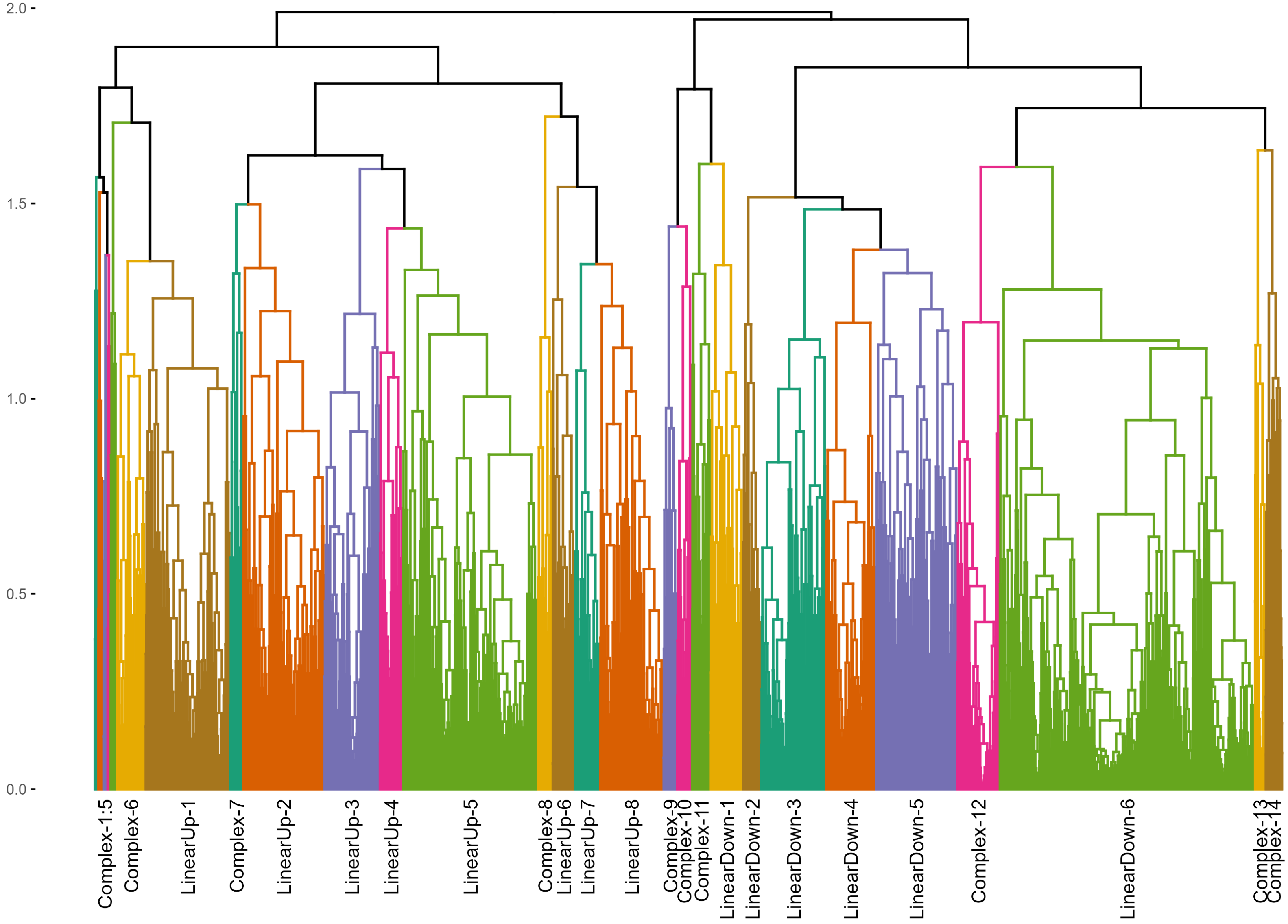


**Supplementary Figure 4:** Hierarchical clustering of 6,142 significant differentially expressed genes into 28 clusters based on their expression trajectories through aging (the dendrogram plots all genes as vertical lines, but individual lines cannot be discriminated in this figure due to the number of genes). Each colored group represents one of our 28 clusters, with labels along the *x*-axis identifying each cluster. The clusters range in size from LinearDown-6 (containing 1,320 genes) to Complex-4 (containing just 15 genes). Note that five of the clusters at the left side of the figure (Complex-1 to Complex-5) are not individually labeled since they are so close together on the plot.

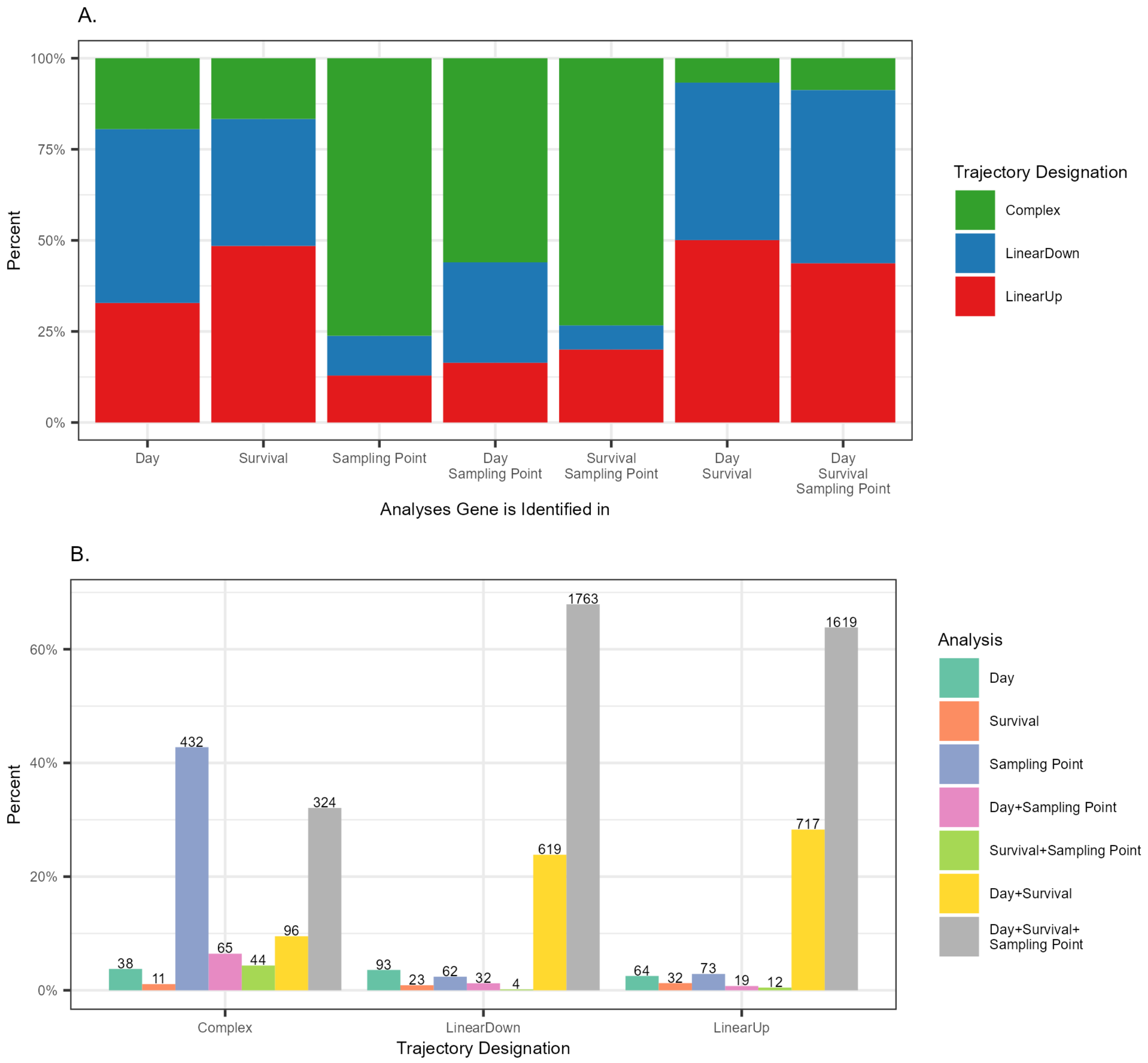


**Supplementary Figure 5:** Differentially expressed genes and their designated trajectories through aging. We executed three statistical analyses, identifying genes with expression changes associated with Day, with Survival, and across Sampling Points. Subsequently, all genes emerging from any of these analyses were designated as belonging to one of the three trajectory groups: LinearUp, LinearDown, or Complex. (See Material and Methods for additional detail). (A) Each column represents the set of genes emerging from at least one analysis, showing the fractions that fall into each of the three trajectory designations. (B) Each bar shows the fraction (*y*-axis) and number (above each bar) of genes within each trajectory designation (Complex, LinearDown, LinearUp) that were identified in each analysis or combination of analyses. Notably, most genes that show a straightforward increase/decrease in expression with age (i.e. LinearUp or LinearDown) were identified in both the Day and Survival analyses, or in all three analyses. However, > 40% of those genes with more complex expression trajectories were found only in the Sampling Point analysis.

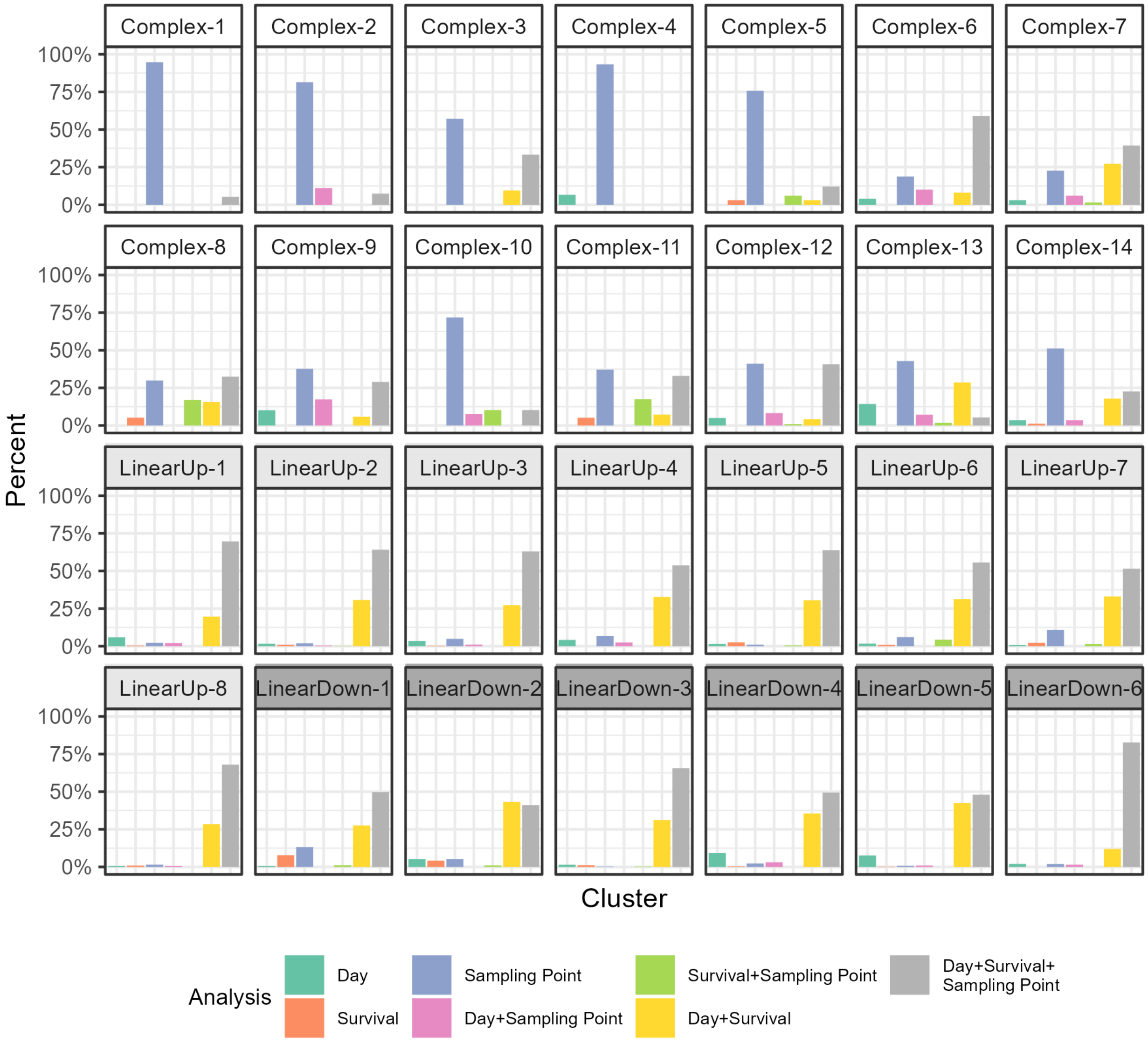


**Supplementary Figure 6:** Examination of the composition of genes in the 28 clusters based on which of the three statistical analyses they were identified in. (This figure is similar to Supplementary Figure 5B but breaks down the three trajectory designations – Complex, LinearUp, LinearDown – into the separate clusters). For the linear clusters (bottom 2 rows of barplots) almost all genes are identified in all three analyses or in the two analyses using continuous variables (Day and Survival). For the complex clusters (top 2 rows of barplots), many genes are found only by the Sampling Point analysis, although there is more variation in composition among the complex clusters than among the linear clusters.

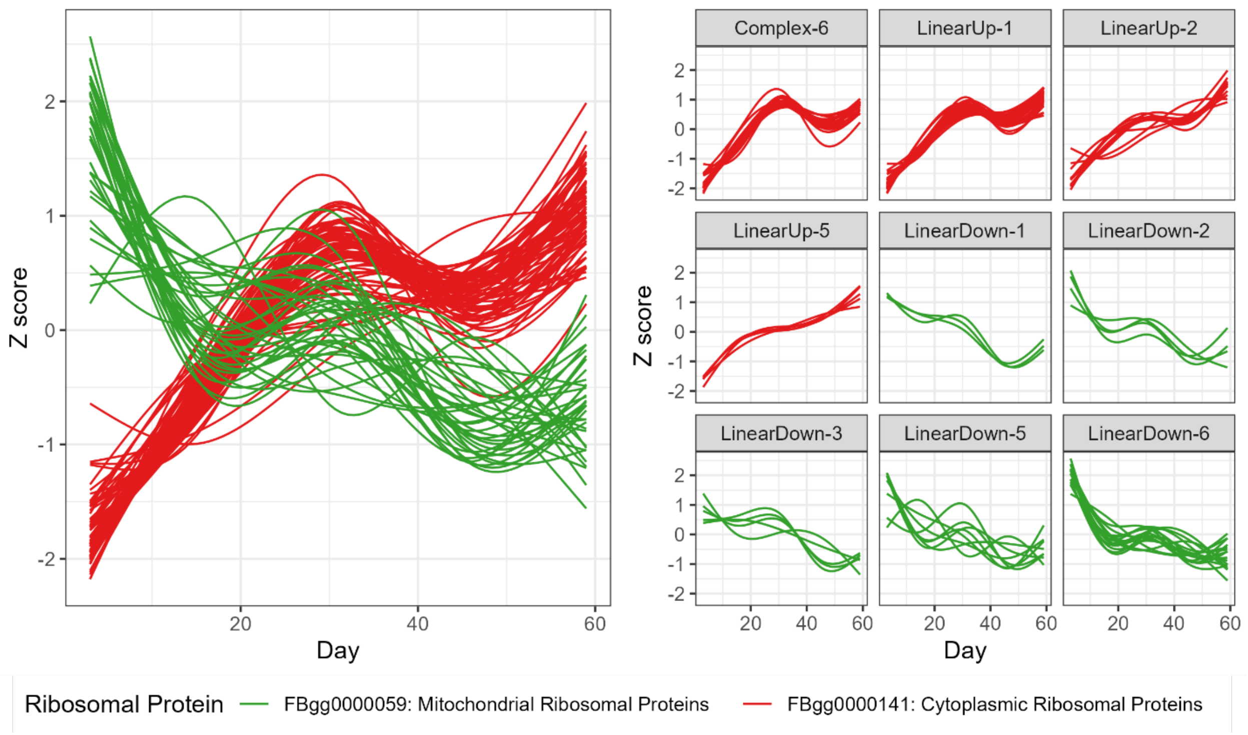


**Supplementary Figure 7:** Expression trajectories of genes encoding mitochondrial and cytoplasmic ribosomal genes. We determined which clusters harbor mitochondrial and cytoplasmic ribosomal protein genes; All mitochondrial ribosomal genes (N=36, green) are in LinearDown clusters, and all cytoplasmic ones (N=80, red) are found either in LinearUp clusters or in the Complex-6 cluster, which has an overall increase in expression.

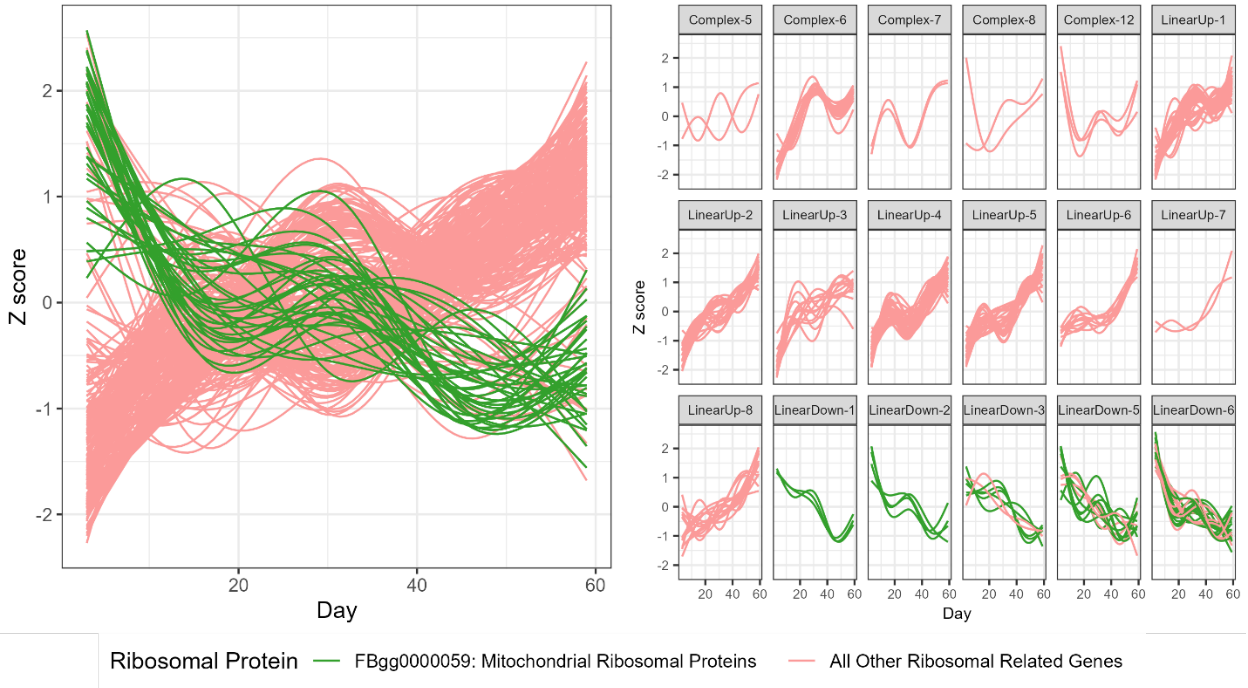


**Supplementary Figure 8:** Examination of the expression trajectories of ribosome-related genes. We compared the expression patterns of genes associated with mitochondrial ribosomal proteins (N=36, green) to the patterns of all other ribosomal-related genes across all clusters (N=234, pink). This latter group includes all those genes from Supplementary Figure 7, along with 154 genes associated with terms which included “ribosome”, “ribosomal”, “rRNA”, or “RNA Polymerase I”. See Supplementary Table L for complete list of terms. As we saw in Supplementary Figure 7, all mitochondrial ribosomal genes are found within LinearDown clusters, while 220/234 ribosome-related genes that are not associated with mitochondrial ribosomal proteins are found within LinearUp or Complex clusters.

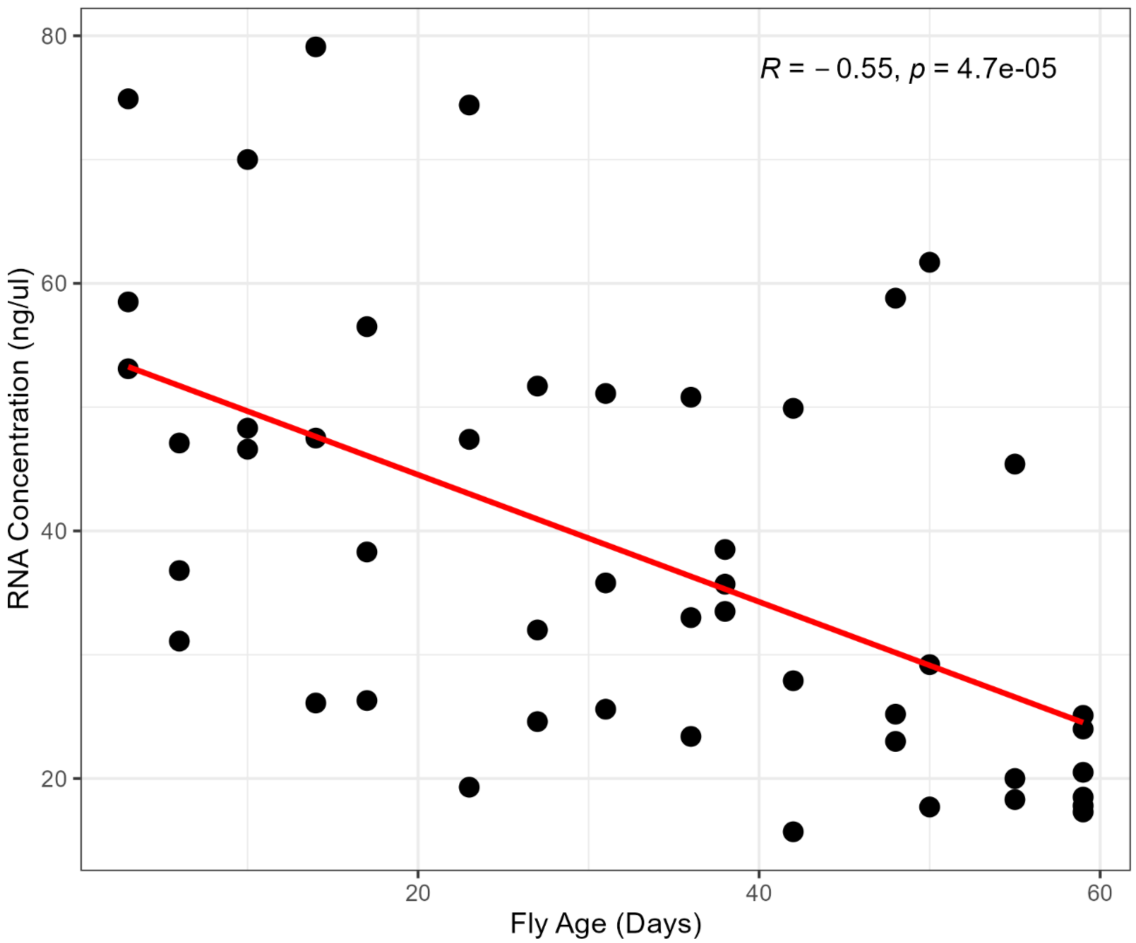


**Supplementary Figure 9:** Correlation between the age of flies and the concentration of the RNA isolated from the heads of sampled flies. The points represent the 48 RNA samples, and are plotted based on the day they were collected and the concentration of the isolated RNA. Note that RNA was isolated from 10 fly heads in all cases. Age and concentration have a Pearson's correlation coefficient of –0.55, which is significant ($p < 10^{-4}$).

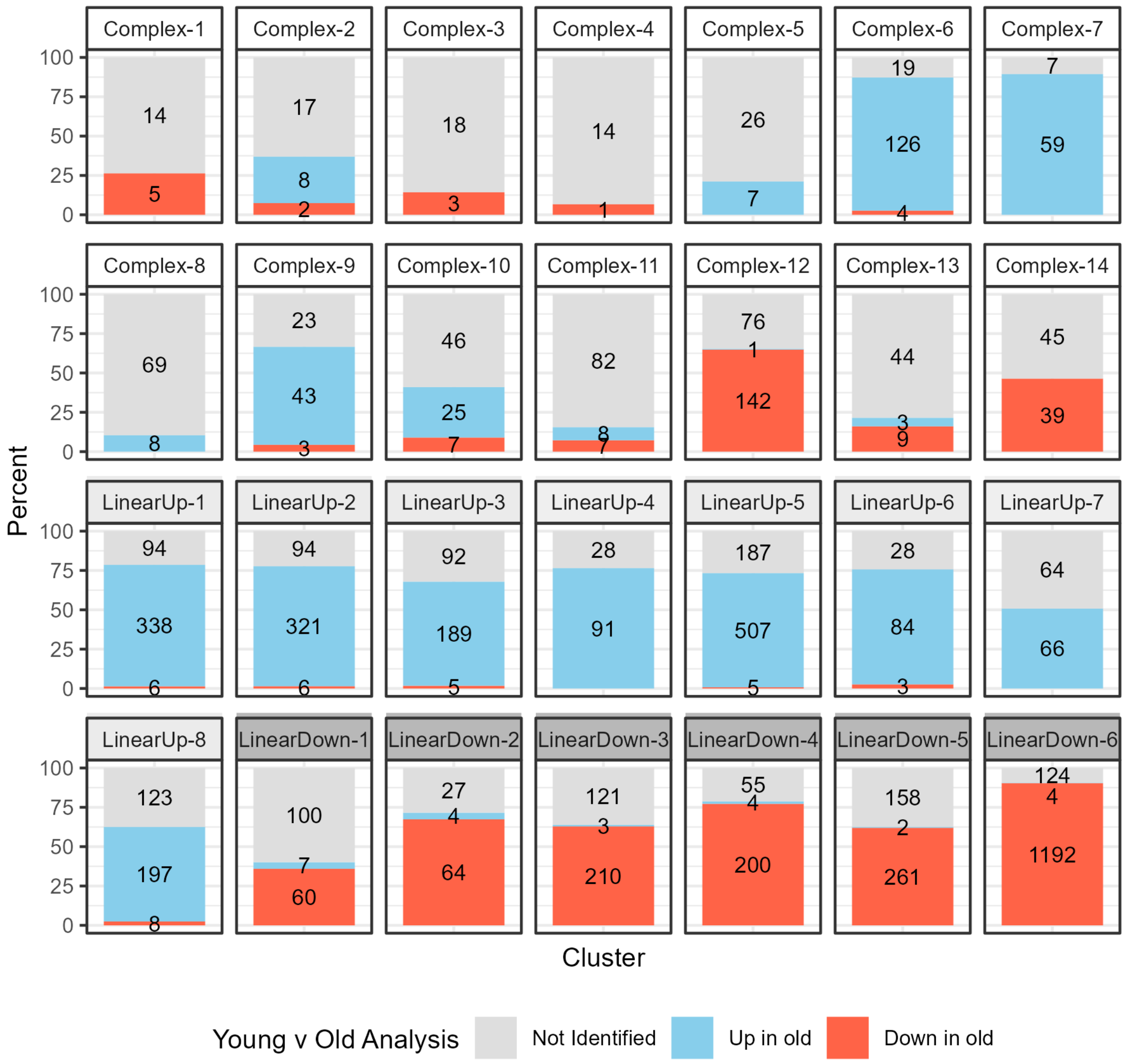


**Supplementary Figure 10:** The fraction of genes in each trajectory-based cluster that were identified in our young versus old (YvO) re-analysis (where we compare samples from Days 3+6 to samples from Day 59). Each bar shows the percent (*y*-axis) and number (within bars) of genes in the cluster that show an increase in expression in old animals in the YvO re-analyses (blue), that decrease in expression in old animals (red), or that were not identified in the re-analysis (gray). For all 8 LinearUp clusters, most of the genes were identified in the YvO analysis, and as anticipated were called as increasing in expression with age. Similarly, for the LinearDown clusters, most of the genes are re-identified in the YvO analysis and are called as decreasing in expression with age. LinearDown-1 is something of an exception since nearly 60% of genes in this cluster were not identified in the YvO re-analysis. Interestingly, for 10 of the 14 complex clusters, the majority of genes were not identified in the YvO analysis, and there is a tendency for those that are re-identified to predominantly be identified as changing in the same direction (either up or down) for a given cluster (e.g. for cluster Complex-12 nearly all identified genes go down with age).

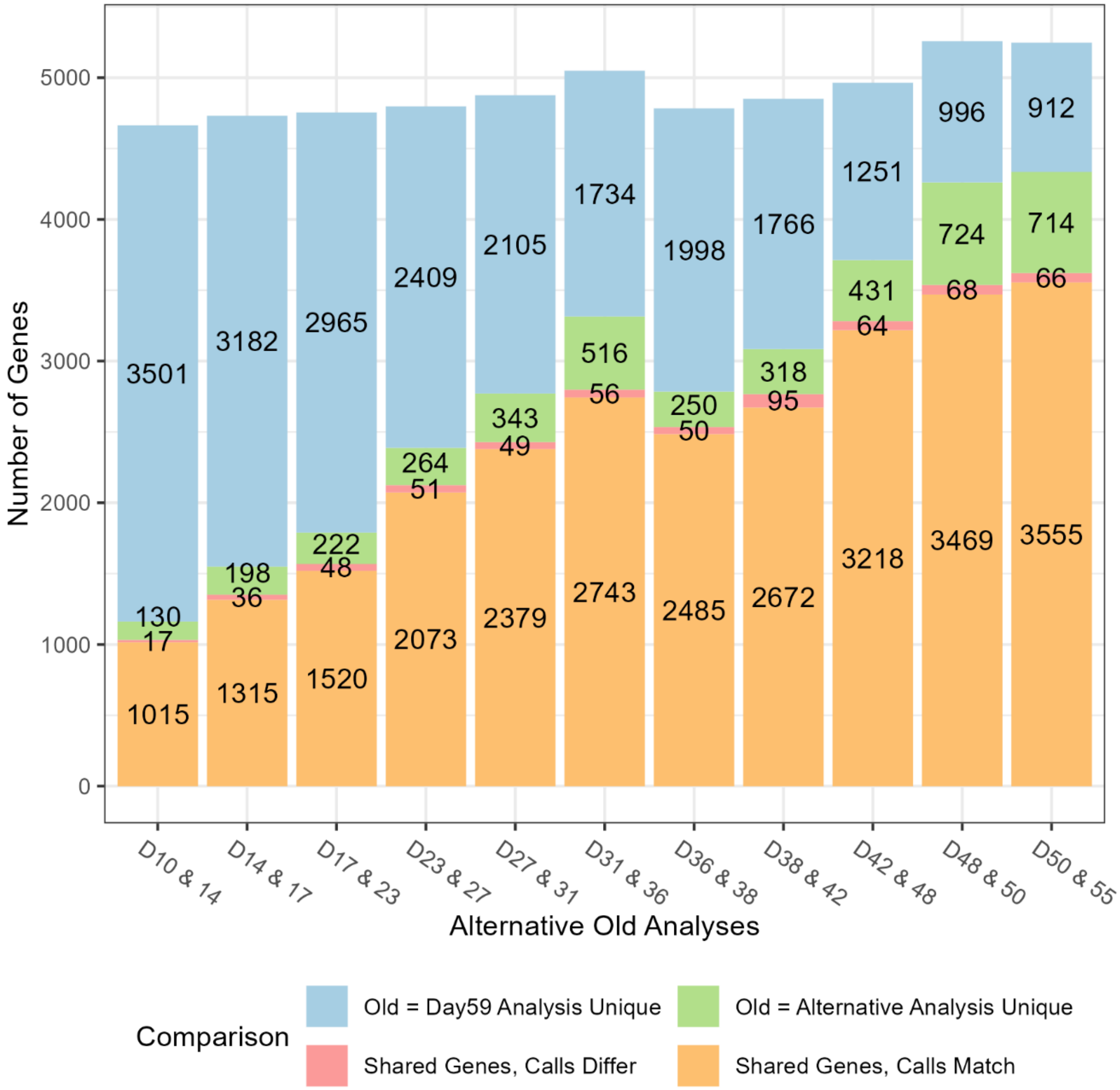


**Supplementary Figure 11:** Comparison of different young versus old analyses. Our initial young versus old analysis identified genes that increased or decreased in expression between Day 3+6 and Day 59. To determine the impact of the old time point used, we repeated this analysis but with alternative samples representing the "old" timepoint (e.g. in the first bar we are comparing Day 3+6 to Day 10+14, in the second bar comparing Day 3+6 to Day 14+17, and so on). Each bar shows the total number of genes identified in the initial Day 59 analysis and in the alternative old timepoint analysis. Genes identified in both analyses that share the same direction call (i.e. both up or both down in expression) are in orange. Genes identified in both analyses but with different direction calls are in red. Genes only identified in the alternative old analysis are in green and those only identified in the Day 59 old analysis are in blue. As the "old" sample becomes chronologically older there is an increase in the number of genes seen. Genes that are identified in both analyses tend to have matching direction calls (orange), with at most 3.4% of shared genes differing in calls (red). But the alternative analyses do often identify unique genes not identified in our original Day 3+6 versus Day 59 analysis (green).

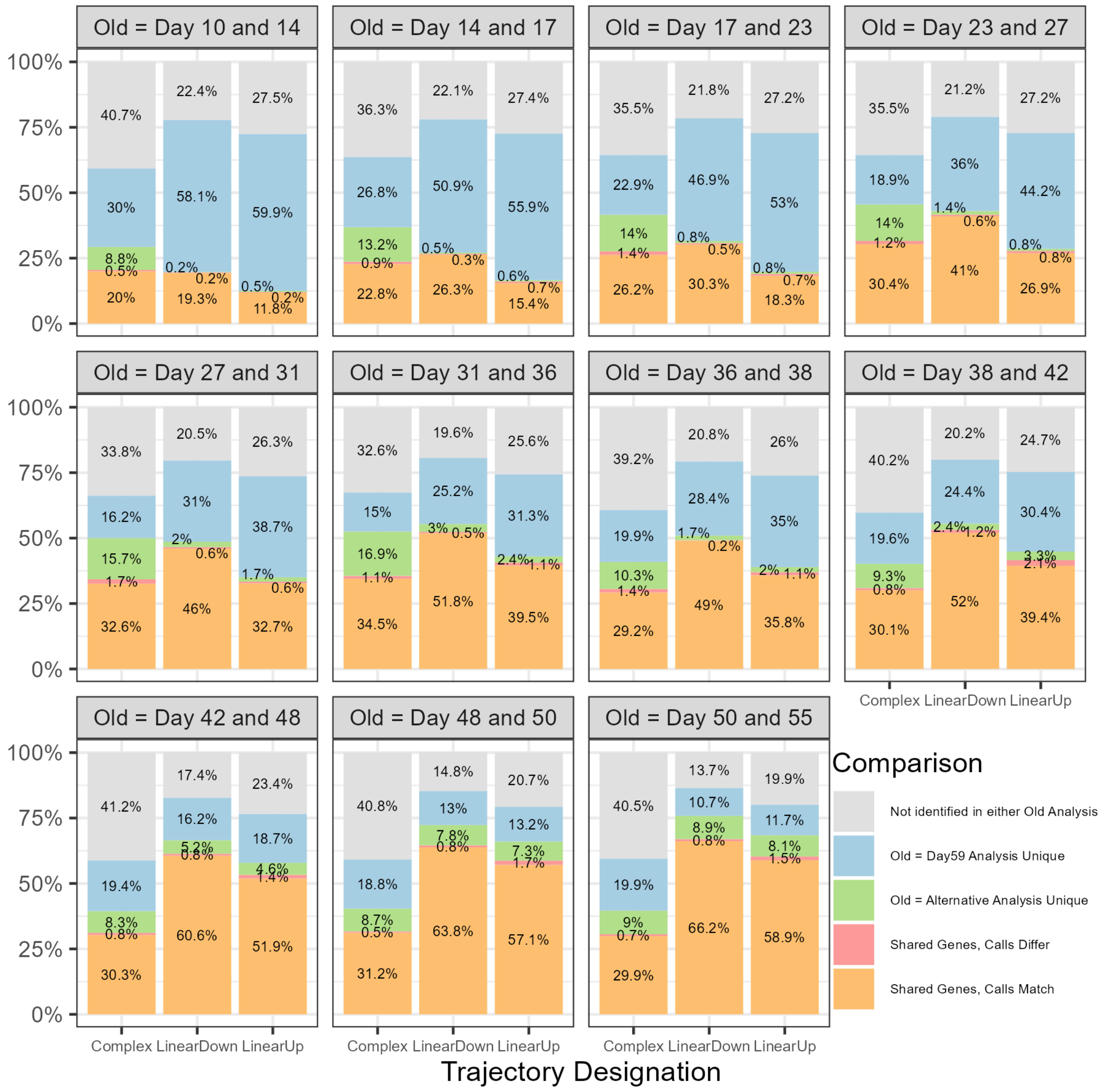


**Supplementary Figure 12:** Comparison of different young versus old time point analyses across cluster trajectory designations. Similar to Supplementary Figure 11 we wished to compare the results of our initial two-timepoint Day 3+6 versus Day 59 analysis with analyses that vary the "aged" timepoint, particularly in terms of the trajectory designation (Complex, LinearUp, LinearDown) of the genes identified in our multi-timepoint analyses. Each of the plots represents one alternative two-timepoint analysis with the chosen timepoints labeled at the top (e.g. "Old=Day10 and Day 14" means Day 3+6 versus Day 10+14). Orange = genes identified in both two-timepoint analyses and that share the same direction call. Red = genes identified in both but with different direction calls. Green = genes only identified in the alternative old analysis. Blue = genes only identified in the Day 59 old analysis. Gray = genes only identified in the multi-timepoint trajectory analysis (and in neither two-timepoint analyses). Almost all genes shared between the two young versus old analyses are called in the same direction regardless of trajectory designation. However, genes with a complex expression trajectory can often be unique to the alternative analysis using a different "aged" timepoint (see the relatively larger green segments of the "Complex" bar in each plot).

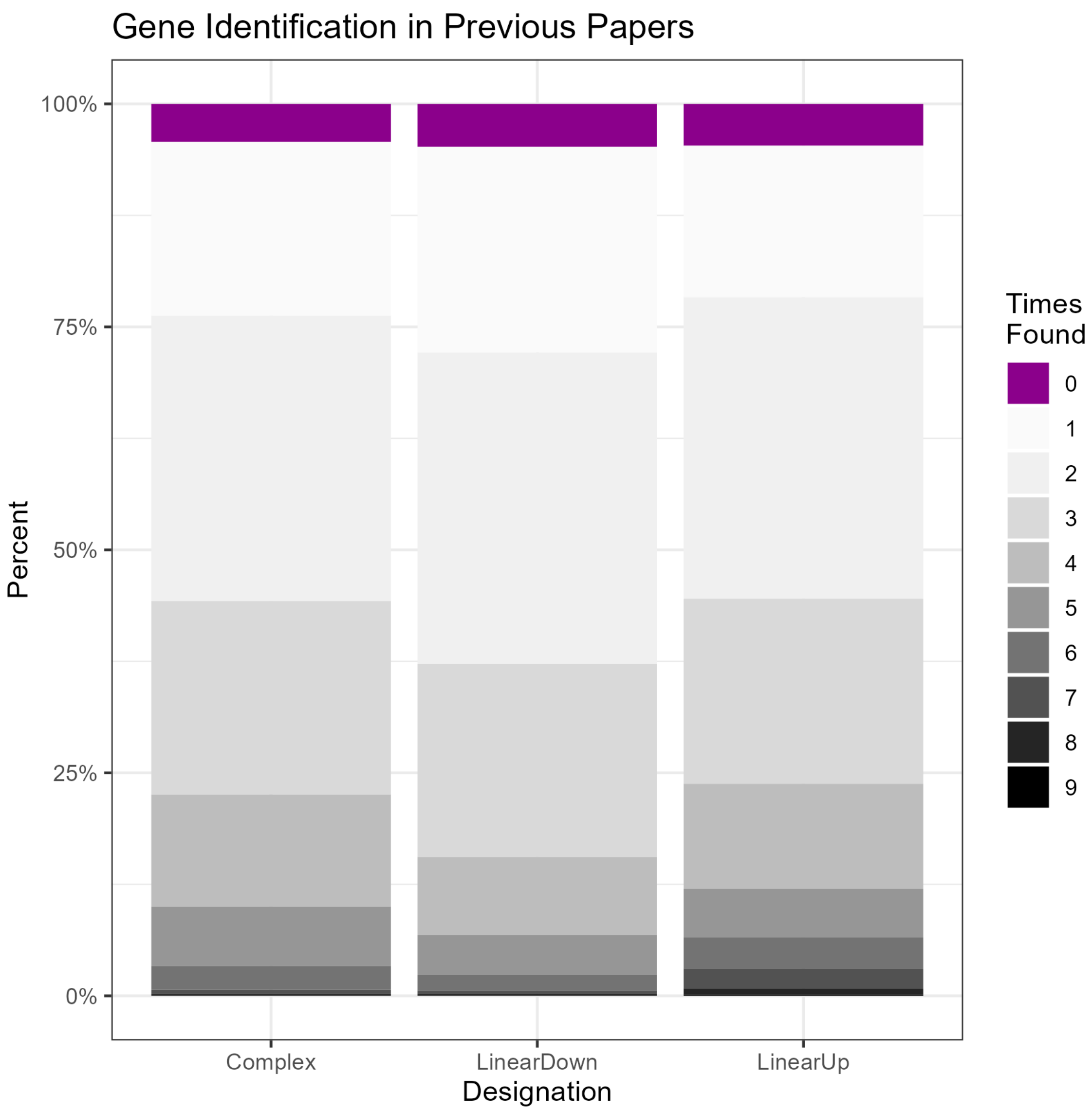


**Supplementary Figure 13:** We examined whether our gene expression trajectory classifications (Complex, LinearUp, LinearDown) are associated with the number of times a gene has been shown to change with age in previous studies. For each of the three trajectory designations we obtained the fraction of genes found in 9 comparison datasets. The fractions are similar across all three trajectories, and for each designation only a small fraction are unique to our analysis (purple).

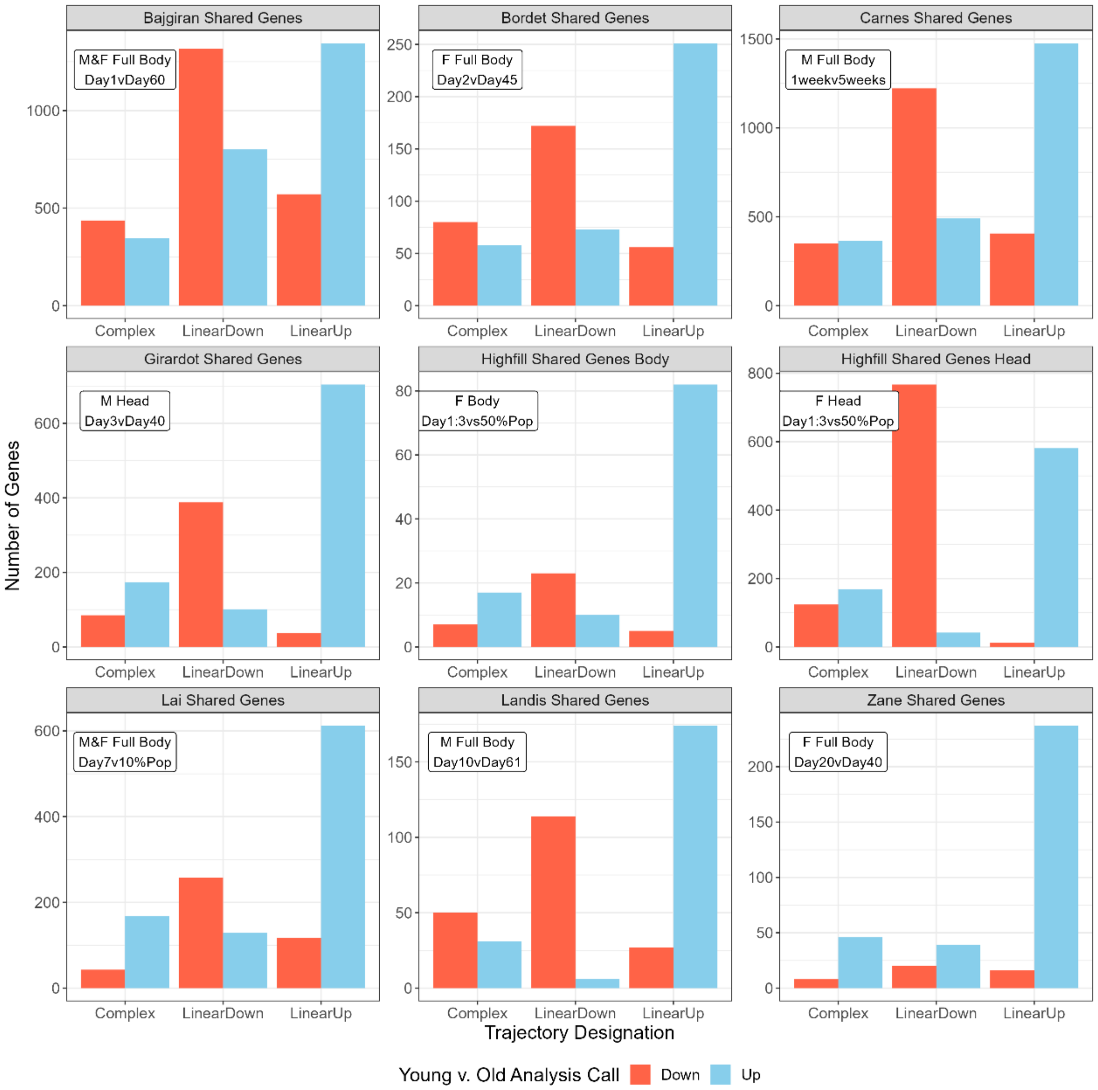


**Supplementary Figure 14**: Information on the genes we identify that are shared with previously published *D. melanogaster* aging expression datasets that compare "young" and "old" animals. Each of the 9 plots shows a single dataset, shared genes are split into our three expression trajectories (Complex, LinearDown, LinearUp), and are further color-coded based on the direction of expression called in the previous work (decreased expression with age = red, increased = blue). In the inset for each plot the sex (M = males, F = Females), tissue and ages of flies used for each published dataset are given. For those shared genes we placed in LinearDown/LinearUp clusters, other studies generally show the anticipated result; most genes are called Down or Up in expression, respectively. However, for those genes we designate as having a Complex age-related expression pattern, results from prior works are more mixed; For instance, for the Carnes study (top right), the Complex set of shared genes are approximately 50:50 Down:Up. We note that the Zane study (bottom right) is an exception, and the majority of all shared genes were called as being up-regulated during aging. We speculate that this discrepancy may be because the "young" group of flies used in that study are – based on a chronological age of 20 days – somewhat older than employed in the other studies.

**Supplementary Text A.** Ingredients and brief protocol for Macdonald lab cornmeal-yeast-molasses fly media.

28.5-liters water
318-g agar (Genesee Scientific; 66-111. This is based on a gel strength of 960 g/cm2, and will change depending on the batch)

- Add water to steam kettle, turn on electric mixer, and slowly add agar
- Bring mix to a boil

3,200-ml molasses (Genesee Scientific; 62-117)

- Reduce the kettle pressure to reduce the heat slightly
- Add molasses, and bring mix back to a boil

4-liters water
1,460-g inactive dry yeast (Genesee Scientific; 62-107)

- Mix in bucket using paint-stirring drill attachment

4-liters water
2,600-g yellow cornmeal (Genesee Scientific; 62-101)

- Mix in bucket using paint-stirring drill attachment
- Add both the water/yeast and water/cornmeal mixes to the steam kettle
- Bring mix back to boil, and simmer for ~15-min
- Release pressure from steam kettle, but continue to stir with electric mixer

330-ml water
259-ml propionic acid (ThermoFisher; A258-500)
31-ml phosphoric acid (85%; ThermoFisher; A242-500)

- Pour mix into kettle

400-ml 95% ethanol
1.5-g tegosept (Genesee Scientific; 20-258)

- Dissolve tegosept in ethanol Pour mix into kettle
- Fill vials/bottles

**Supplementary Text B.** Protocol for RNA isolation from 10 fly heads.

I. RNA Purification

1) Add 300µl cold TRIzol to a screw-top tube already containing 4-6 glass beads and 10 fly heads.
2) Use Mini-BeadBeater-96 (BioSpec Products) for 45sec to homogenize, then briefly/gently spin down each tube
3) Add 300µl of 100% EtOH.
4) Invert tube 10 times to mix.
5) Spin at maximum speed (14,000rpm) for 1min.
6) Transfer sample to spin column in collection tube (avoid beads/tissue)
7) Spin at maximum speed for 30sec. Discard flow through. Transfer column to new collection tube.

II. DNase Treatment

8) Add 400µl of RNA Wash Buffer to column.
9) Spin at maximum speed for 30sec.
10) Add 40µl of DNase mastermix** to column.
11) Incubate sample at RT for 15min.

III. Wash

12) Add 400µl of RNA Prewash to column.
13) Spin at maximum speed for 30 sec. Discard flow through.
14) Repeat steps 12 and 13.
15) Add 700µl of RNA Wash Buffer to column.
16) Spin at maximum speed for 2min.
17) Transfer column to final pre-labeled 1.7ml tube

IV. Elute

18) Add 15µl of DNase/RNase-free water to column.
19) Spin at maximum speed for 30sec.
20) Discard column and move forward to QC sample.

** DNase mastermix: Mix 35µl DNA Digestion Buffer and 5µl reconstituted DNase I for the appropriate number of samples (use the actual number + 1).

- TRIzol (LifeTech/Fisher, Cat Num 15596018)
- Tubes, spin columns, RNA Wash Buffer, DNase mastermix components, RNA Prewash, and DNase/RNase-free water (Zymo Direct-zol RNA MicroPrep kit, Cat Num R2062)

SUPPLEMENTARY TEXT LEGENDS

**Supplementary Text C:** R code to access our Gene and Cluster Shiny app and Cluster Enrichment Shiny app. Includes code to install the necessary packages for the apps. *Available as a text file.*

**Supplementary Text D:** A walkthrough describing how to access and use our shiny apps and their features. The guide includes screenshots of the apps, examples of the utility of the apps, and descriptions of their output. *Available as a PDF*.

SUPPLEMENTARY TABLE LEGENDS

**Supplementary Table A:** Details of how the A4 male aging cohort was maintained and sampled. Information in the columns is as follows: Date (in 2017), Fly_age (in days; since flies were allowed to emerge over a period of 48 hours, a fly age of T days should be interpreted as the interval T−2 to T days old), Num_deaths_per_day, Num_flies_sampled (note that not all sampled flies were used to generate RNAseq data), Sampling_time (incubator lights turned on at 8:00am, so flies were typically sampled 3-4 hours following lights on), Cohort_tipped (all flies were moved to new vials every 2-3 days, and sometimes – see "Notes" – flies were re-arrayed into groups following anesthesia), Num_flies_dead_cumulative, Num_flies_sampled_cumulative, Expected_num_flies_present (starting from the known number of animals collected – 5,330 – the expected number of flies remaining on any given day; that the number is not zero on the final day is explained by fly loss during tipping), and Notes (provides additional detail about how the cohort was treated). *Available as a tab-delimited text file*.

**Supplementary Table B:** Details of RNA isolation and mRNAseq library preparation and quality control. Information in the columns is as follows: RNA_isolation_batch, Date (when RNA was isolated), Fly_age (in days; since flies were allowed to emerge over a period of 48 hours, a fly age of T days should be interpreted as the interval T−2 to T days old), Replicate_ID (each of the replicate samples from a given timepoint were given a letter-based identifier), Used_for_library (only a subset of 48 RNA samples were used to produce mRNAseq libraries), Conc_RNA_ng_ul (measured via a NanoDrop ND-1000), Amount_starting_RNA_ng (the amount of total RNA used in the library prep), Volume_RNA_ul, Volume_Water_ul, Sample_Name, Plate_well_ID (following RNA isolation in tubes, RNA was moved to wells of a 96-well plate for library construction), Library_batch, Conc_library_ng_ul (measured via a Qubit), TapeStation_frag_size_bp. *Available as a tab-delimited text file*.

**Supplementary Table C:** For each of the 15 sampling points, records the day of collection, the calculated survivorship of the population, and how many samples were collected. *Available as a csv file*.

**Supplementary Table D:** Properties of gene clusters. We varied the total number of clusters from 15 to 40, and for each set recorded the minimum and maximum number of genes in any cluster, the first and third quartiles for cluster size, and the median and mean cluster sizes. We ideally wanted clusters with 50-500 genes, so we additionally recorded the number of clusters, and the number of genes in clusters that were either "low" (clusters < 50 genes), "in range" (cluster with 50 – 500 genes), or "high" (clusters > 500 genes). As we increased the number of clusters used, we also recorded how many genes moved to an "in range" cluster (from a "high" cluster) and how many genes moved out of an "in range" cluster (to a "low" cluster). Using these values, we narrowed down our ideal number to 27, 28, or 29 clusters due to a large gain of genes in our ideal range when increasing to 27 clusters, but losing genes in that range after splitting into 30 clusters. Supplementary Figure 3 shows detail of cluster gene sizes for 27, 28, and 29 clusters. *Available as a csv file*.

**Supplementary Table E:** Results of linear regressions to classify clusters into trajectories (Complex, LinearUp, LinearDown). A cluster was designated as linear if the *p*-value was less than 0.002 (0.05/28) and complex otherwise. Linear clusters were further designated as being up or down (expression increasing or decreasing with aging) based on the linear regression coefficient. Cluster designations were consistent across all three analyses performed. Cluster names are composed of its designation and a number based on its position in the dendrogram (see Supplementary Figure 4). *Available as a csv file*.

**Supplementary Tables F-K:** Downloaded raw PANGEA results for all 28 clusters from our 6 analyses. (**F**) SLIM2 GO BP, (**G**) SLIM2 GO CC, (**H**) SLIM2 GO MF, (**I**) DRSC GLAD Gene Groups, (**J**) Flybase Gene Groups, (**K**) REACTOME pathways. *Available as a csv file*.

**Supplementary Table L**: Ribosomal related terms from PANGEA analyses. Includes each term's set, ID, name, and the number of genes associated with the term. Terms were included if their name contained "ribosome", "ribosomal", "rRNA", or "RNA Polymerase I".

**Supplementary Table M**: Previously published aging expression datasets used. For each dataset we include identifying paper information such as first author, published year, and PMID. The number of genes identified in the original paper whose ID we were able to validate, and how many of those genes are shared with our identified genes, is also included. We additionally note information regarding sample collection (such as fly sex, the target tissue, and the time point comparison employed). We have also included where in the original paper the data we used can be found. *Available as a csv file*.
