## Supplementary material for "Dynamic Changes in Gene Expression Through Aging in *Drosophila melanogaster* Heads": All Supplementary Materials: Supplementary Text C.txt

#Paste the following into a R script to have acess to our interactive shiny apps.

#R Script to access Gene Identification and Cluster enrichment Shiny apps
#Created: 8/22/2024

#You will need the following packages:
#Install if needed
install.packages("shiny")
install.packages("bslib")
install.packages("tidyverse")
install.packages("rms")
install.packages("DT")

library(shiny)

#Run the following to have acesss to our Gene and Cluster app. 
#This allows users to look up specific genes, and receive information regarding their
#expression trajectory, cluster placement, and identification in young vs old analyses.
runGitHub("Gene_and_Cluster_Shiny", "Hanson19")


#Run the following to have access to our Cluster enrichment app.
#This allows users to view a cluster's enriched term, and have the ability to compare
#clusters and their enriched terms. You can select specific clusters you are interested in,
#and choose which types of terms you want shown. 
runGitHub("Enrichment_Shiny", "Hanson19")
