## Supplementary material for "Dynamic Changes in Gene Expression Through Aging in *Drosophila melanogaster* Heads": All Supplementary Materials: Supplementary Text D.pdf

### Visual Breakdown of Shiny Apps

- This document serves as a visual walkthrough/breakdown of the Gene and Cluster and Cluster Enrichment shiny apps.
- This document will walk through how to access the apps using the code available in Supplemental Text C, how to use the different app features available in the sidebars, and breakdown each app’s output.
- This guide is broken down into sections, with the first section walking through the Gene and Cluster app and the second walking through the Cluster Enrichment app.

### Gene and Cluster App Breakdown

To access shiny apps, paste code available in Supplementary Text C into R script, install necessary packages, and read in shiny package.

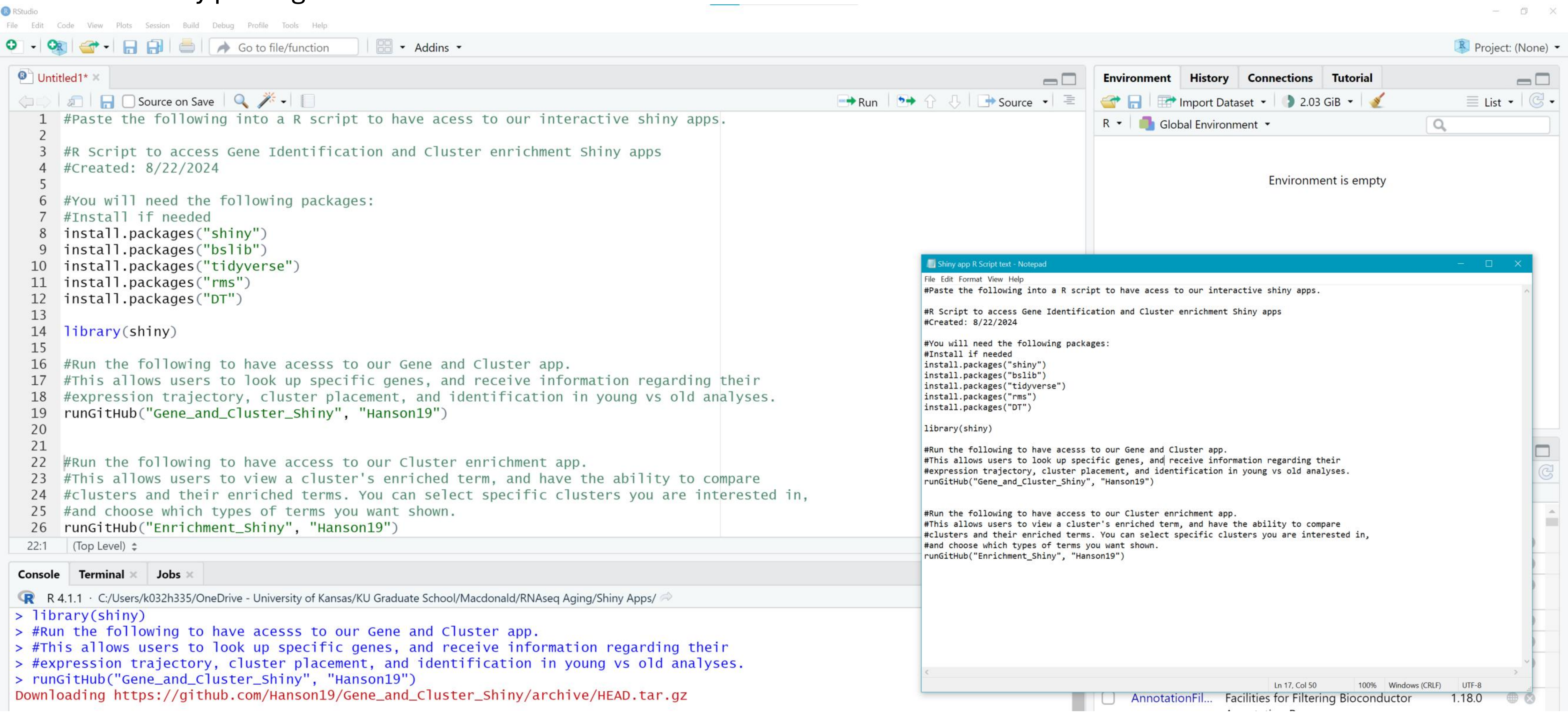


To access the Gene and Cluster app, run runGitHub(“Gene_and_Cluster_Shiny”, Hanson19) on line 19 of the screenshot.

A new window should pop up and look like this.
We have provided a brief description for each part of the app.

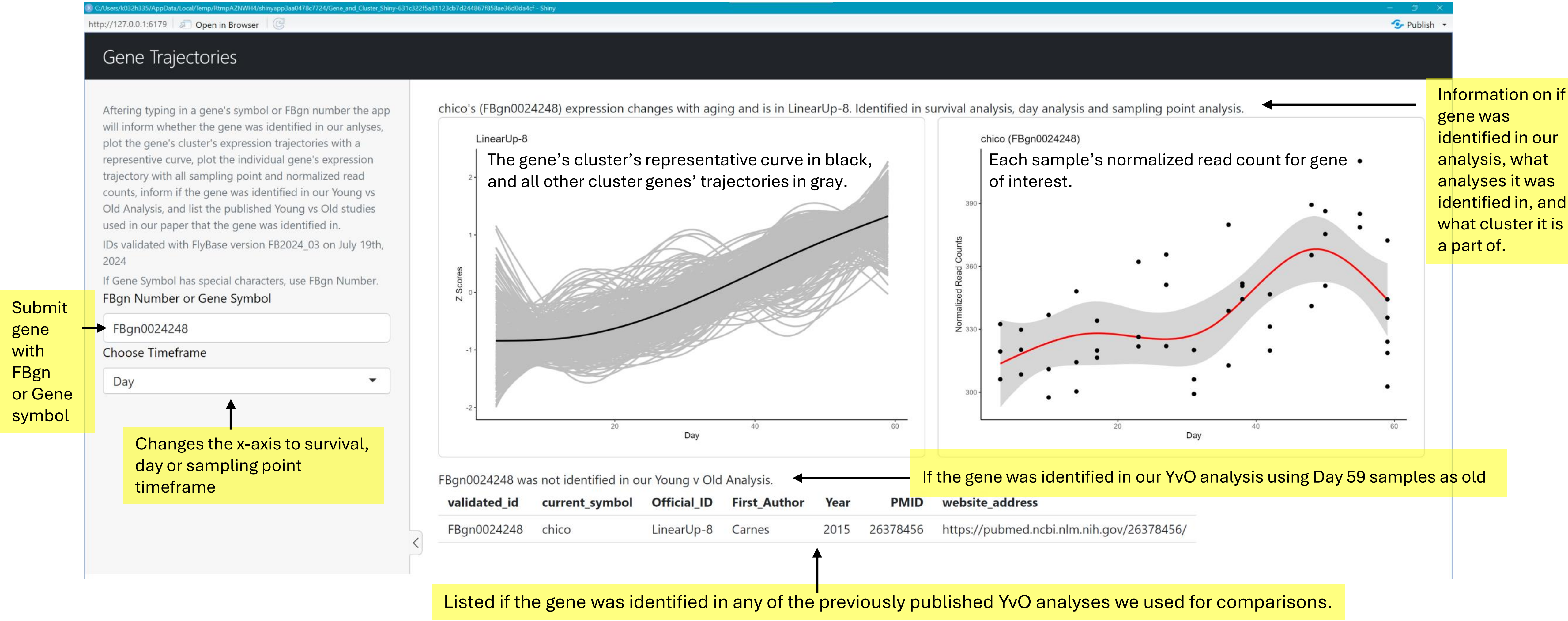

### Additional example now using Gene symbol and a different timeframe

C:/Users/k032h335/AppData/Local/Temp/Rtmp44cJhG/shinyapp1fa85b286050/Gene_and_Cluster_Shiny-631c322f5a81123cb7d244867f858ae36d0da4cf - Shiny

http://127.0.0.1:3104 | Open in Browser | Publish

#### Gene Trajectories

Aftering typing in a gene's symbol or FBgn number the app will inform whether the gene was identified in our anlyses, plot the gene's cluster's expression trajectories with a representive curve, plot the individual gene's expression trajectory with all sampling point and normalized read counts, inform if the gene was identified in our Young vs Old Analysis, and list the published Young vs Old studies used in our paper that the gene was identified in.

IDs validated with FlyBase version FB2024_03 on July 19th, 2024

If Gene Symbol has special characters, use FBgn Number.

FBgn Number or Gene Symbol

Indy

Choose Timeframe

Survival

Indy's (FBgn0036816) expression changes with aging and is in Complex-6. Identified in sampling point analysis.

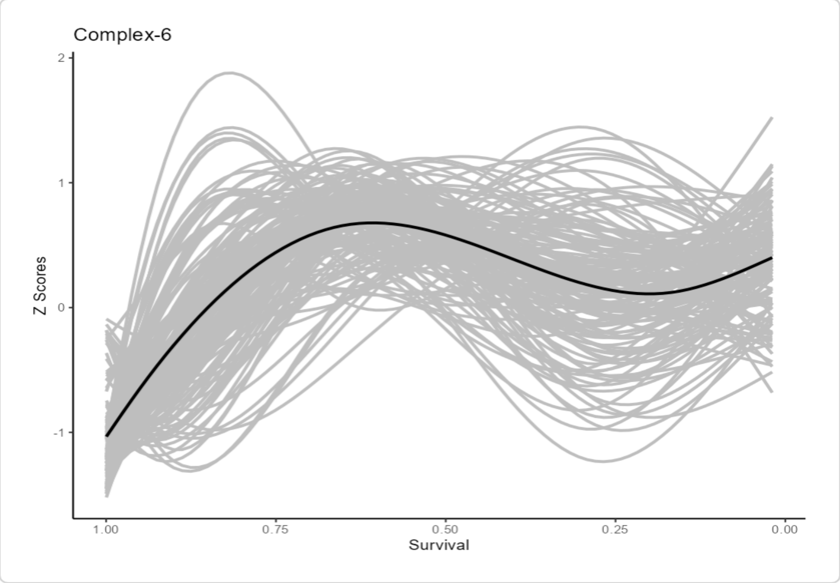


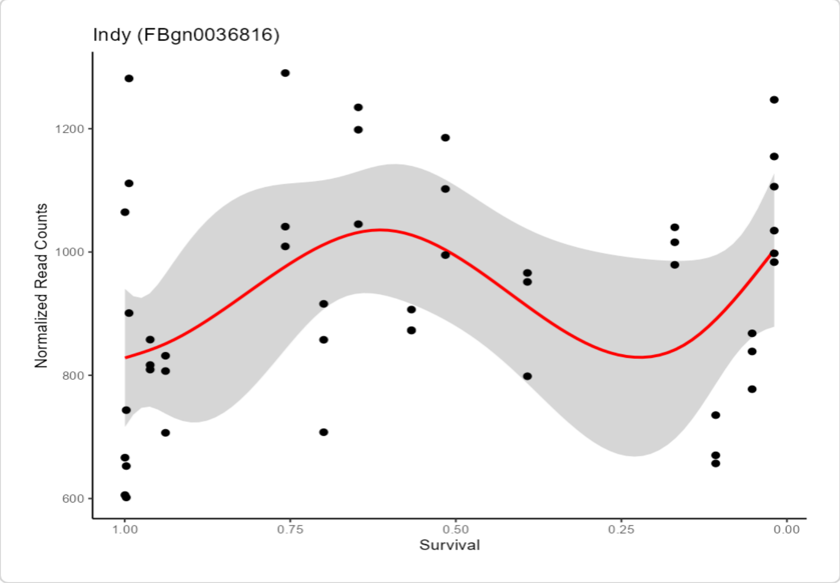


Indy (FBgn0036816) was identified in our Young v Old Analysis.

| validated_id | current_symbol | Official_ID | First_Author | Year | PMID | website_address |
|---|---|---|---|---|---|---|
| FBgn0036816 | Indy | Complex-6 | Carnes | 2015 | 26378456 | https://pubmed.ncbi.nlm.nih.gov/26378456/ |
| FBgn0036816 | Indy | Complex-6 | Giradot | 2006 | 16584578 | https://pubmed.ncbi.nlm.nih.gov/16584578/ |
| FBgn0036816 | Indy | Complex-6 | Highfill | 2016 | 27485207 | https://pubmed.ncbi.nlm.nih.gov/27485207/ |
| FBgn0036816 | Indy | Complex-6 | Lai | 2007 | 17196240 | https://pubmed.ncbi.nlm.nih.gov/17196240/ |

If a gene’s expression does not significantly change with aging, the app will still plot the gene’s expression trajectory.

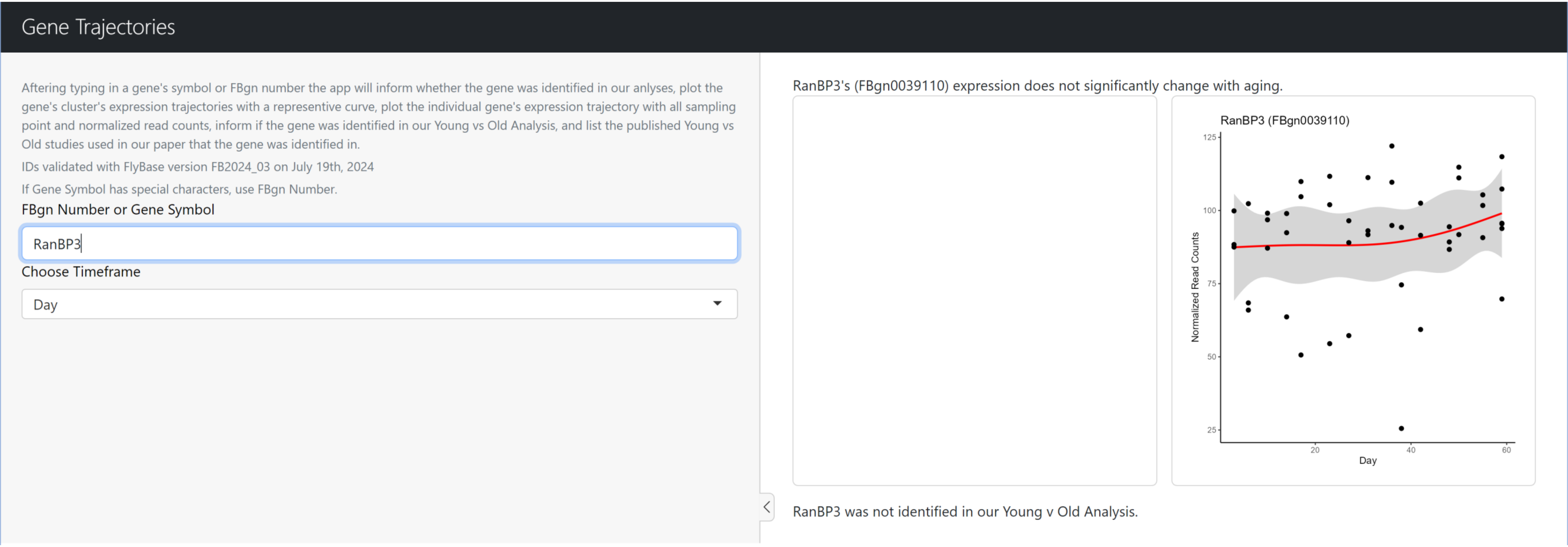

If the submitted gene symbol or FBgn is not found within our data, the app will state that it was not identified.
We do suggest to double check the spelling of the submitted ID to be sure. If using a gene symbol, we also suggest to try the gene's FBgn in case the gene symbol has changed.
As a reminder IDs were validated with FlyBase version FB2024_03 on July 19th, 2024.

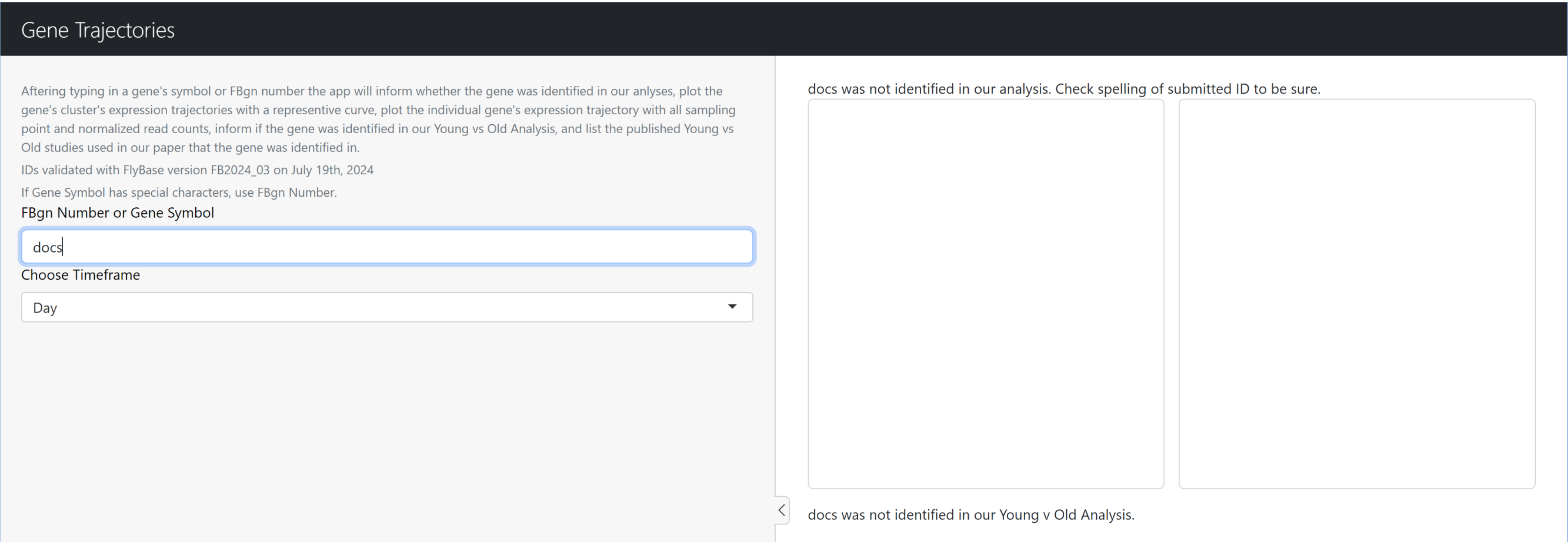

### Cluster Enrichment App Breakdown

To access the cluster enrichment app , run runGitHub(“Enrichment_Shiny”, Hanson19) on line 26 of the screenshot.

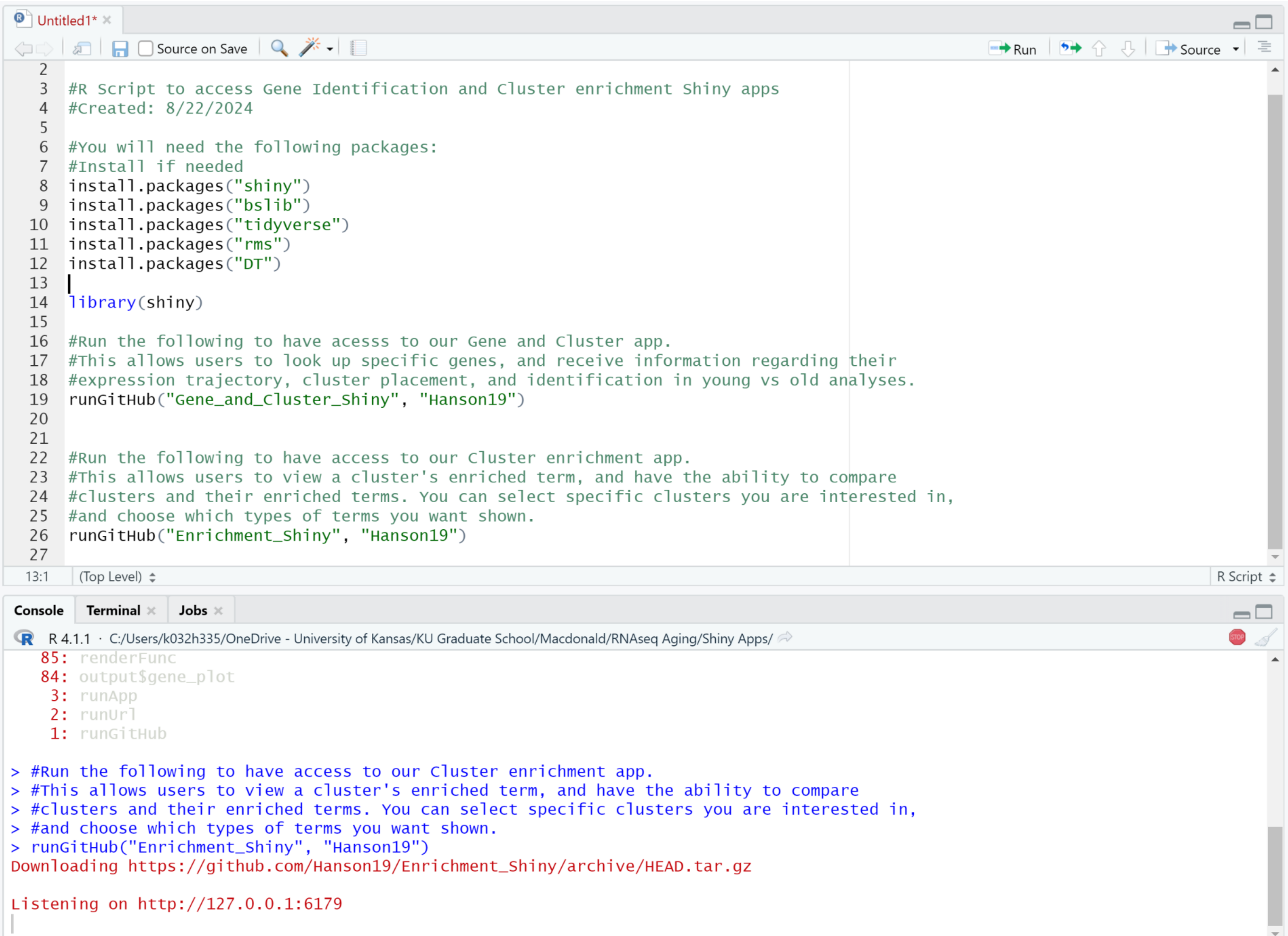

A window that looks like this should pop up. The output of this app is a lot bigger than that of the gene and cluster app so it may not all fit on one page and require scrolling.

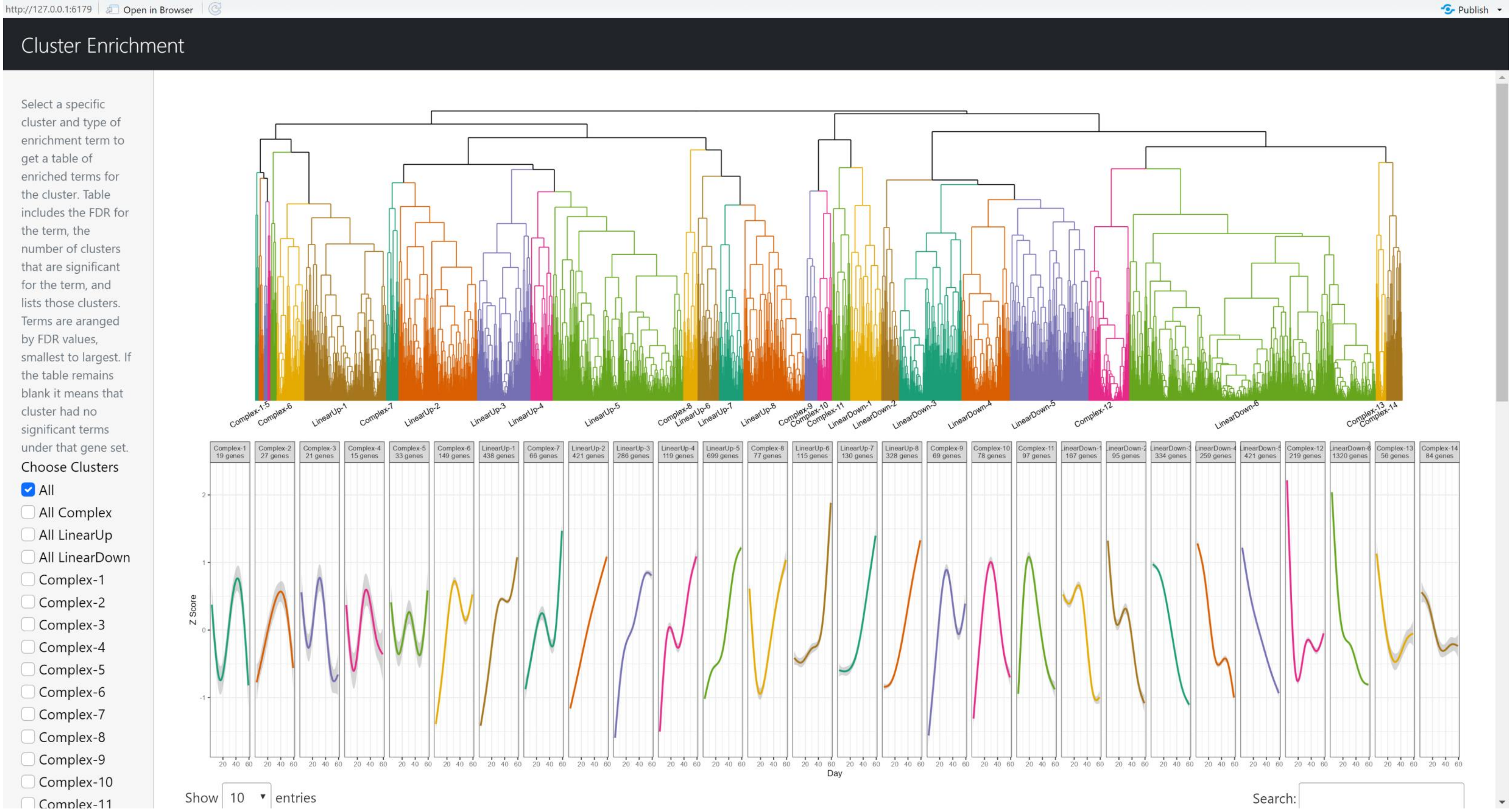

### Sidebar features:

Select which clusters you want to see enriched terms for. You can choose to look at multiple clusters at the same time. App defaults to “All”.

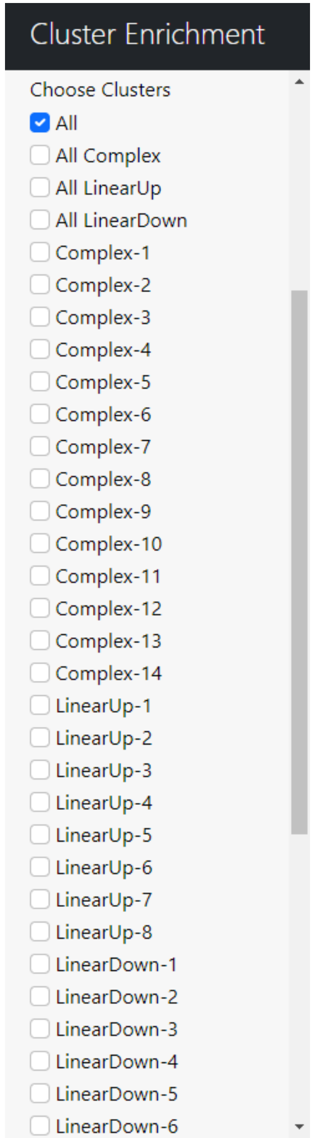


Select what type of terms you want to see. Can look at multiple types at the same time.

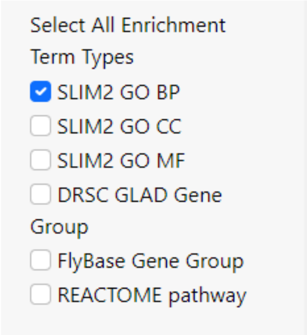


If multiple clusters are selected, you can choose to show all terms the clusters are enriched. You can also compare clusters by either only showing terms that are shared between selected clusters or terms that are unique to one of the selected clusters.

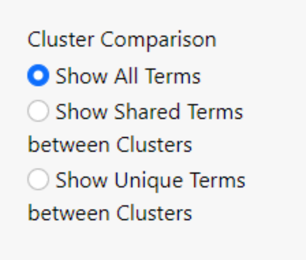

Dendrogram in the top panel, and representative cluster expression curves in the second panel.
The dendrogram will highlight clusters that are selected, and only the representative curves of selected clusters will be plotted.

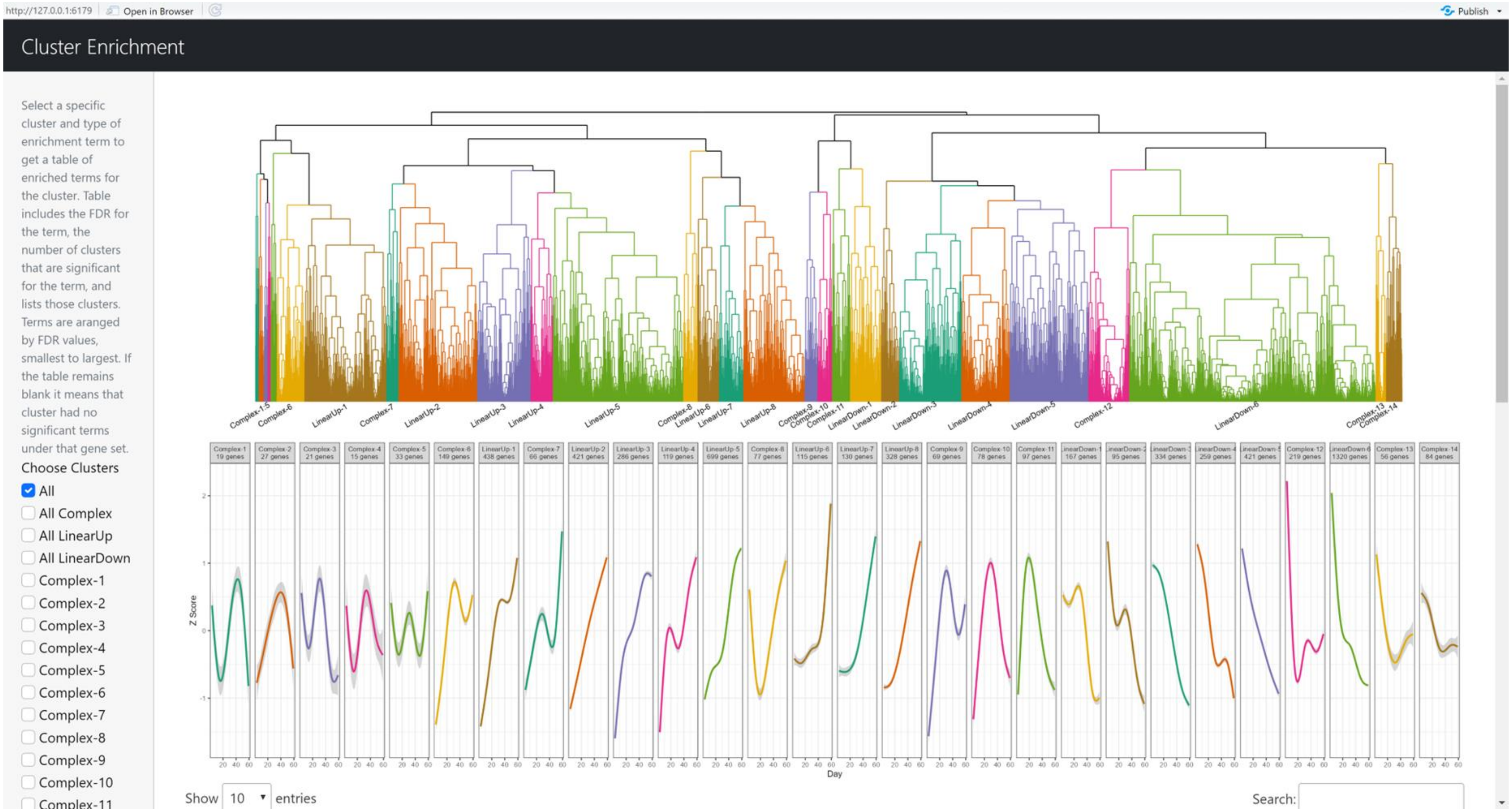

Only Complex clusters are selected in this example.

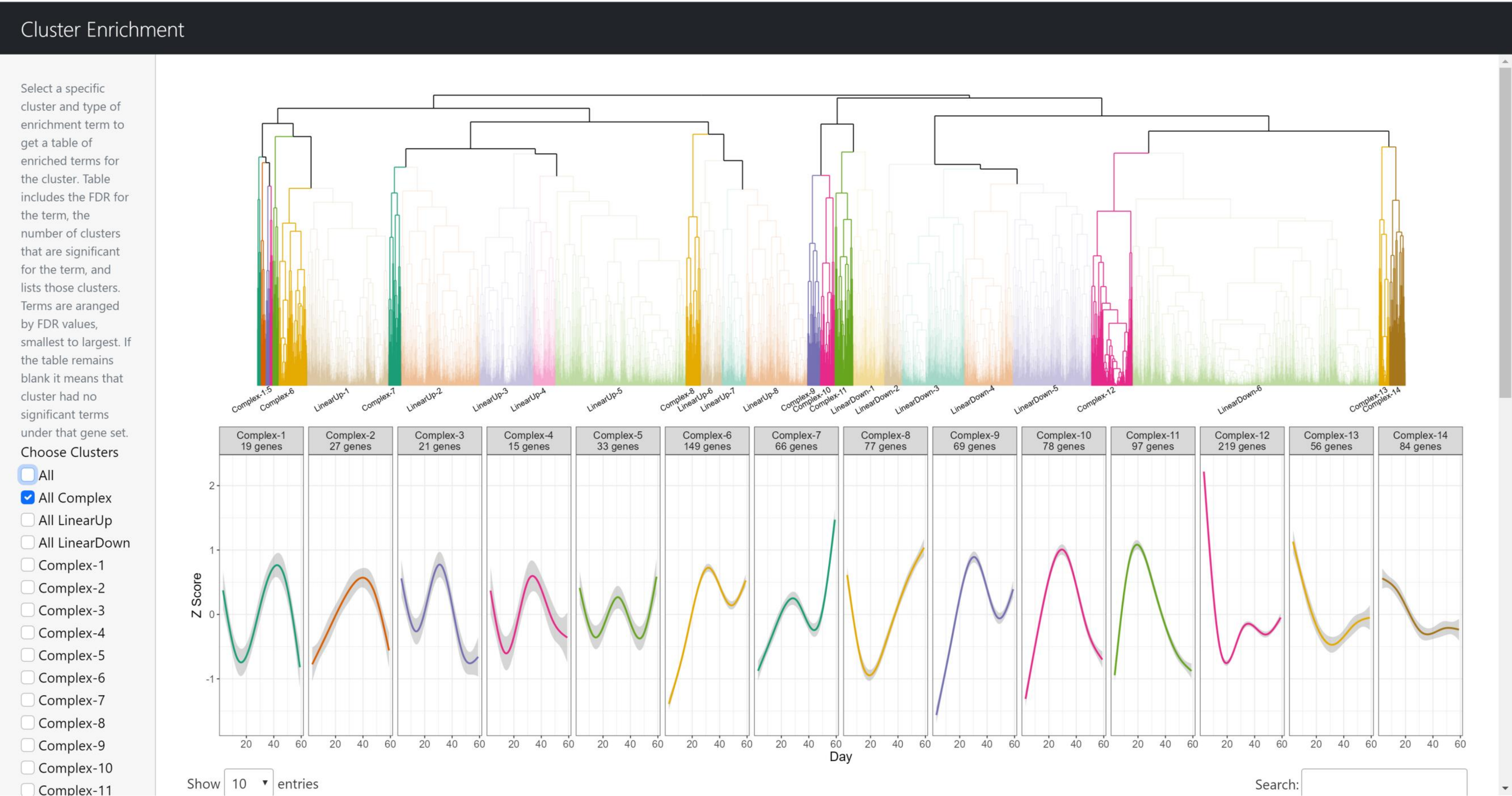

Only Complex-6 and LinearUp-1 are selected here.

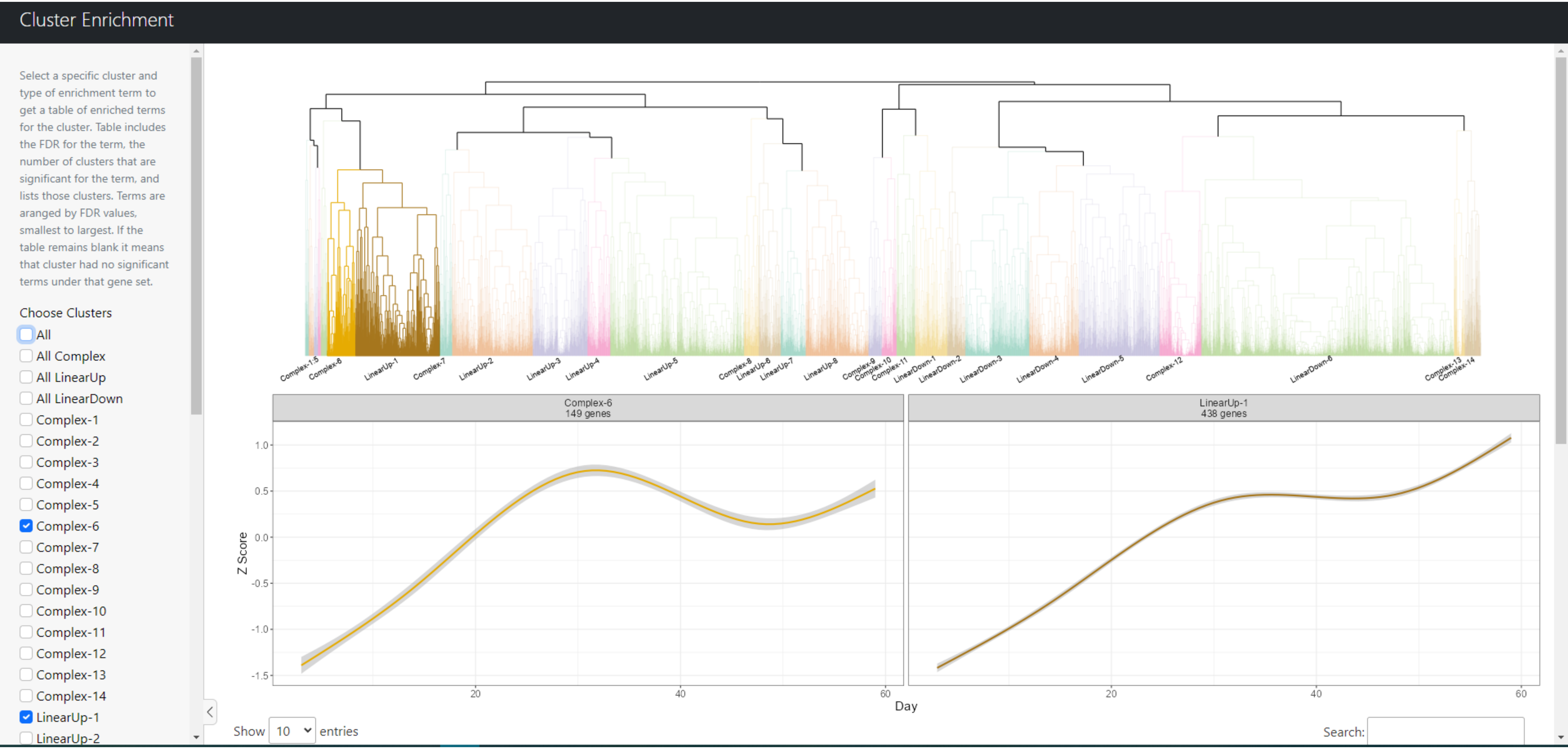

The last panel is a table of the selected cluster's enriched terms. Here we are listing all the enriched SLIM2 GO BP terms in all the clusters based on selections in our sidebar.

#### Cluster Enrichment

- [ ] Complex-13
- [ ] Complex-14
- [ ] LinearUp-1
- [ ] LinearUp-2
- [ ] LinearUp-3
- [ ] LinearUp-4
- [ ] LinearUp-5
- [ ] LinearUp-6
- [ ] LinearUp-7
- [ ] LinearUp-8
- [ ] LinearDown-1
- [ ] LinearDown-2
- [ ] LinearDown-3
- [ ] LinearDown-4
- [ ] LinearDown-5
- [ ] LinearDown-6

Select All Enrichment Term Types

- [x] SLIM2 GO BP
- [ ] SLIM2 GO CC
- [ ] SLIM2 GO MF
- [ ] DRSC GLAD Gene Group
- [ ] FlyBase Gene Group
- [ ] REACTOME pathway

Cluster Comparison

- (•) Show All Terms
- ( ) Show Shared Terms between Clusters
- ( ) Show Unique Terms between Clusters

Show 10 entries

Search:

| | Gene.Set | Gene.Set.ID | Gene.Set.Name | Cluster | FDR | Num.Cluster.Genes | Gene.Set.Size | Total.Num.Cluster.Genes | Num.Sig.Clusters | All.Sig.Clusters |
|---|---|---|---|---|---|---|---|---|---|---|
| 1 | SLIM2 GO BP | GO:0042254 | ribosome biogenesis | LinearUp-4 | 5.5e-24 | 34 | 206 | 119 | 4 | LinearUp-1, LinearUp-2, LinearUp-4, LinearUp-6 |
| 2 | SLIM2 GO BP | GO:0002181 | cytoplasmic translation | LinearUp-1 | 8e-24 | 42 | 128 | 438 | 3 | Complex-6, LinearUp-1, LinearUp-2 |
| 3 | SLIM2 GO BP | GO:0006811 | ion transport | LinearDown-6 | 8.1e-24 | 136 | 514 | 1320 | 2 | Complex-12, LinearDown-6 |
| 4 | SLIM2 GO BP | GO:0002181 | cytoplasmic translation | Complex-6 | 3.07e-21 | 24 | 128 | 149 | 3 | Complex-6, LinearUp-1, LinearUp-2 |
| 5 | SLIM2 GO BP | GO:0016072 | rRNA metabolic process | LinearUp-4 | 6.13e-19 | 21 | 138 | 119 | 3 | LinearUp-1, LinearUp-4, LinearUp-6 |
| 6 | SLIM2 GO BP | GO:0050808 | synapse organization | LinearDown-6 | 1.02e-11 | 72 | 295 | 1320 | 3 | LinearDown-3, LinearDown-4, LinearDown-6 |
| 7 | SLIM2 GO BP | GO:0006091 | generation of precursor metabolites and energy | LinearDown-6 | 1.11e-10 | 57 | 218 | 1320 | 3 | Complex-12, Complex-3, LinearDown-6 |
| 8 | SLIM2 GO BP | GO:0016192 | vesicle-mediated transport | LinearUp-7 | 1.32e-09 | 25 | 510 | 130 | 5 | LinearDown-4, LinearDown-6, LinearUp-5, LinearUp-7, LinearUp-8 |
| 9 | SLIM2 GO BP | GO:0007399 | nervous system development | LinearDown-4 | 1.6e-09 | 56 | 1112 | 259 | 5 | LinearDown-3, LinearDown-4, LinearDown-6, LinearUp-2, LinearUp-5 |
| 10 | SLIM2 GO BP | GO:0006281 | DNA repair | LinearUp-3 | 2.82e-09 | 22 | 179 | 286 | 2 | LinearUp-2, LinearUp-3 |

Showing 1 to 10 of 141 entries

Previous 1 2 3 4 5 ... 15 Next

### Breakdown of enriched terms table:

- Term identification information → Gene.Set, Gene.Set.ID, Gene.Set.Name
- Enriched cluster → Cluster
- Benjamini Hochberg false discovery rate → FDR
- Number of genes in cluster associated with term → Num.Cluster.Genes
- Total number of genes associated with term → Gene.Set.Size
- Total number of genes in the cluster → Total.Num.Cluster.Genes
- Total number of enriched clusters → Num.Sig.Clusters
- List of all enriched clusters → All.Sig.Clusters

| | Gene.Set | Gene.Set.ID | Gene.Set.Name | Cluster | FDR | Num.Cluster.Genes | Gene.Set.Size | Total.Num.Cluster.Genes | Num.Sig.Clusters | All.Sig.Clusters |
|---|---|---|---|---|---|---|---|---|---|---|
| 1 | SLIM2 GO BP | GO:0042254 | ribosome biogenesis | LinearUp-4 | 5.5e-24 | 34 | 206 | 119 | 4 | LinearUp-1, LinearUp-2, LinearUp-4, LinearUp-6 |
| 2 | SLIM2 GO BP | GO:0002181 | cytoplasmic translation | LinearUp-1 | 8e-24 | 42 | 128 | 438 | 3 | Complex-6, LinearUp-1, LinearUp-2 |
| 3 | SLIM2 GO BP | GO:0006811 | ion transport | LinearDown-6 | 8.1e-24 | 136 | 514 | 1320 | 2 | Complex-12, LinearDown-6 |
| 4 | SLIM2 GO BP | GO:0002181 | cytoplasmic translation | Complex-6 | 3.07e-21 | 24 | 128 | 149 | 3 | Complex-6, LinearUp-1, LinearUp-2 |

Can use search bar to lookup terms containing certain words.
Here we have pulled out enriched terms across all clusters that are part of SLIM2 GO BP and MF sets that have "ribosome" within their name.

Cluster Enrichment

- [ ] Complex-13
- [ ] Complex-14
- [ ] LinearUp-1
- [ ] LinearUp-2
- [ ] LinearUp-3
- [ ] LinearUp-4
- [ ] LinearUp-5
- [ ] LinearUp-6
- [ ] LinearUp-7
- [ ] LinearUp-8
- [ ] LinearDown-1
- [ ] LinearDown-2
- [ ] LinearDown-3
- [ ] LinearDown-4
- [ ] LinearDown-5
- [ ] LinearDown-6

Select All Enrichment Term Types

- [x] SLIM2 GO BP
- [ ] SLIM2 GO CC
- [x] SLIM2 GO MF
- [ ] DRSC GLAD Gene Group
- [ ] FlyBase Gene Group
- [ ] REACTOME pathway

Cluster Comparison

- (•) Show All Terms
- ( ) Show Shared Terms between Clusters
- ( ) Show Unique Terms between Clusters

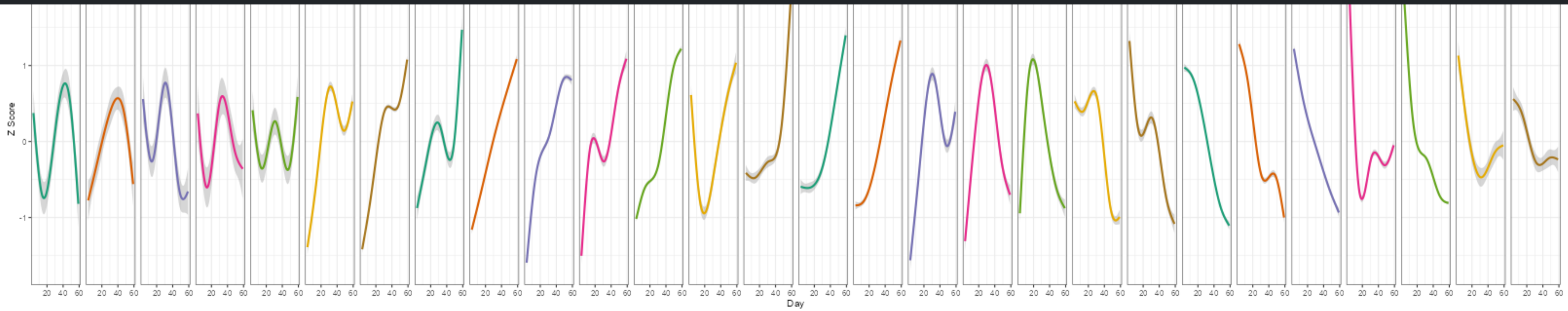


Show 10 entries

Search: ribosome

| | Gene.Set | Gene.Set.ID | Gene.Set.Name | Cluster | FDR | Num.Cluster.Genes | Gene.Set.Size | Total.Num.Cluster.Genes | Num.Sig.Clusters | All.Sig.Clusters |
|---|---|---|---|---|---|---|---|---|---|---|
| 1 | SLIM2 GO BP | GO:0042254 | ribosome biogenesis | LinearUp-4 | 5.5e-24 | 34 | 206 | 119 | 4 | LinearUp-1, LinearUp-2, LinearUp-4, LinearUp-6 |
| 4 | SLIM2 GO MF | GO:0003735 | structural constituent of ribosome | LinearUp-1 | 1.34e-21 | 40 | 171 | 438 | 2 | Complex-6, LinearUp-1 |
| 8 | SLIM2 GO MF | GO:0003735 | structural constituent of ribosome | Complex-6 | 4.01e-17 | 23 | 171 | 149 | 2 | Complex-6, LinearUp-1 |
| 26 | SLIM2 GO BP | GO:0042254 | ribosome biogenesis | LinearUp-1 | 7.12e-07 | 25 | 206 | 438 | 4 | LinearUp-1, LinearUp-2, LinearUp-4, LinearUp-6 |
| 81 | SLIM2 GO BP | GO:0042254 | ribosome biogenesis | LinearUp-6 | 0.00201 | 9 | 206 | 115 | 4 | LinearUp-1, LinearUp-2, LinearUp-4, LinearUp-6 |
| 115 | SLIM2 GO BP | GO:0042254 | ribosome biogenesis | LinearUp-2 | 0.0105 | 15 | 206 | 421 | 4 | LinearUp-1, LinearUp-2, LinearUp-4, LinearUp-6 |

Showing 1 to 6 of 6 entries (filtered from 182 total entries)

Previous 1 Next

We can also compare clusters based on the selected cluster comparison option on the sidebar.
Here we are showing terms that are shared by LinearUp-2, LinearUp-5, and LinearUp-8 across all term sets.

#### Cluster Enrichment

- [ ] Complex-13
- [ ] Complex-14
- [ ] LinearUp-1
- [x] LinearUp-2
- [ ] LinearUp-3
- [ ] LinearUp-4
- [x] LinearUp-5
- [ ] LinearUp-6
- [ ] LinearUp-7
- [x] LinearUp-8
- [ ] LinearDown-1
- [ ] LinearDown-2
- [ ] LinearDown-3
- [ ] LinearDown-4
- [ ] LinearDown-5
- [ ] LinearDown-6

Select All Enrichment Term Types

- [x] SLIM2 GO BP
- [x] SLIM2 GO CC
- [x] SLIM2 GO MF
- [x] DRSC GLAD Gene Group
- [x] FlyBase Gene Group
- [x] REACTOME pathway

Cluster Comparison

- ( ) Show All Terms
- (•) Show Shared Terms between Clusters
- ( ) Show Unique Terms between Clusters

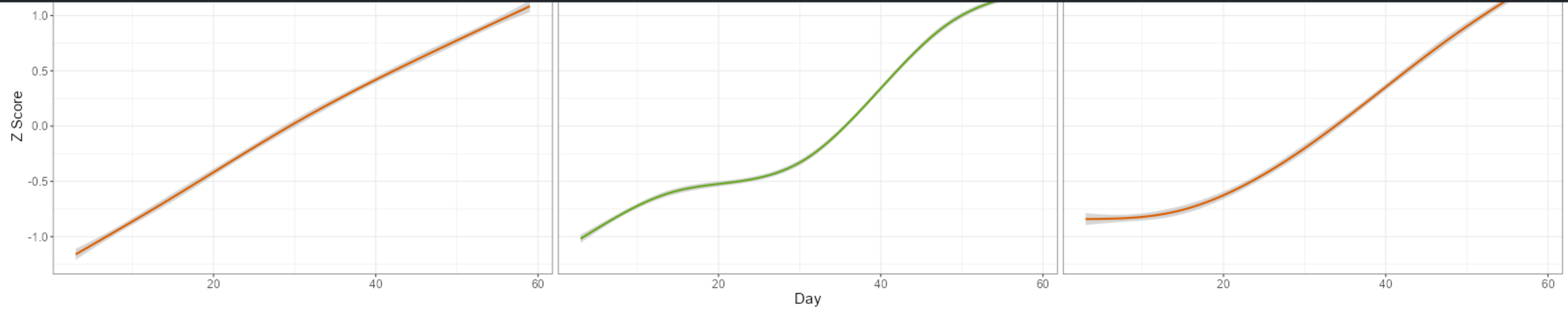


Show 10 entries

Search:

| | Gene.Set | Gene.Set.ID | Gene.Set.Name | Cluster | FDR | Num.Cluster.Genes | Gene.Set.Size | Total.Num.Cluster.Genes | Num.Sig.Clusters | All.Sig.Clusters |
|---|---|---|---|---|---|---|---|---|---|---|
| 1 | SLIM2 GO BP | GO:0006950 | response to stress | LinearUp-2 | 5.37e-09 | 80 | 1197 | 421 | 5 | LinearUp-2, LinearUp-3, LinearUp-5, LinearUp-7, LinearUp-8 |
| 2 | SLIM2 GO BP | GO:0006950 | response to stress | LinearUp-5 | 3.05e-09 | 115 | 1197 | 699 | 5 | LinearUp-2, LinearUp-3, LinearUp-5, LinearUp-7, LinearUp-8 |
| 3 | SLIM2 GO BP | GO:0006950 | response to stress | LinearUp-8 | 0.000351 | 54 | 1197 | 328 | 5 | LinearUp-2, LinearUp-3, LinearUp-5, LinearUp-7, LinearUp-8 |
| 4 | SLIM2 GO BP | GO:0006955 | immune response | LinearUp-2 | 0.00105 | 22 | 287 | 421 | 3 | LinearUp-2, LinearUp-5, LinearUp-8 |
| 5 | SLIM2 GO BP | GO:0006955 | immune response | LinearUp-5 | 1.85e-06 | 38 | 287 | 699 | 3 | LinearUp-2, LinearUp-5, LinearUp-8 |
| 6 | SLIM2 GO BP | GO:0006955 | immune response | LinearUp-8 | 2e-04 | 22 | 287 | 328 | 3 | LinearUp-2, LinearUp-5, LinearUp-8 |

Showing 1 to 6 of 6 entries

Previous 1 Next

Now showing enriched terms in LinearUp-2, 5 and 8 that are not shared with each other.
The term doesn't have to be unique to that cluster, it just can't be found in the other two, as seen in rows 4-7 of the table.

Cluster Enrichment

- [ ] Complex-13
- [ ] Complex-14
- [ ] LinearUp-1
- [x] LinearUp-2
- [ ] LinearUp-3
- [ ] LinearUp-4
- [x] LinearUp-5
- [ ] LinearUp-6
- [ ] LinearUp-7
- [x] LinearUp-8
- [ ] LinearDown-1
- [ ] LinearDown-2
- [ ] LinearDown-3
- [ ] LinearDown-4
- [ ] LinearDown-5
- [ ] LinearDown-6

Select All Enrichment Term Types

- [x] SLIM2 GO BP
- [x] SLIM2 GO CC
- [x] SLIM2 GO MF
- [x] DRSC GLAD Gene Group
- [x] FlyBase Gene Group
- [x] REACTOME pathway

Cluster Comparison

- ( ) Show All Terms
- ( ) Show Shared Terms between Clusters
- (•) Show Unique Terms between Clusters

20 40 60 20 40 60 20 40 60
Day

Show 10 entries Search:

| | Gene.Set | Gene.Set.ID | Gene.Set.Name | Cluster | FDR | Num.Cluster.Genes | Gene.Set.Size | Total.Num.Cluster.Genes | Num.Sig.Clusters | All.Sig.Clusters |
|---|---|---|---|---|---|---|---|---|---|---|
| 1 | FlyBase Gene Group | FBgg0000721 | BOMANINS | LinearUp-5 | 1.58e-07 | 9 | 12 | 699 | 1 | LinearUp-5 |
| 2 | DRSC GLAD Gene Group | GLAD:24590 | Major signaling pathways | LinearUp-5 | 1.72e-05 | 40 | 330 | 699 | 1 | LinearUp-5 |
| 3 | SLIM2 GO BP | GO:0060541 | respiratory system development | LinearUp-5 | 3.89e-05 | 32 | 252 | 699 | 1 | LinearUp-5 |
| 4 | FlyBase Gene Group | FBgg0000215 | ENDOSOMAL SORTING COMPLEXES REQUIRED FOR TRANSPORT | LinearUp-5 | 5.5e-05 | 9 | 20 | 699 | 2 | LinearUp-1, LinearUp-5 |
| 5 | SLIM2 GO BP | GO:0051301 | cell division | LinearUp-2 | 0.000127 | 24 | 269 | 421 | 3 | LinearUp-2, LinearUp-3, LinearUp-7 |
| 6 | SLIM2 GO BP | GO:0051321 | meiotic cell cycle | LinearUp-2 | 0.000129 | 21 | 221 | 421 | 2 | LinearUp-2, LinearUp-3 |
| 7 | SLIM2 GO BP | GO:0000278 | mitotic cell cycle | LinearUp-2 | 0.000173 | 34 | 487 | 421 | 2 | LinearUp-2, LinearUp-3 |
| 8 | SLIM2 GO BP | GO:0006260 | DNA replication | LinearUp-2 | 0.000241 | 13 | 103 | 421 | 1 | LinearUp-2 |
| 9 | FlyBase Gene Group | FBgg0000390 | CYTOPLASMIC AMINOACYL-TRNA SYNTHETASES | LinearUp-5 | 0.000925 | 7 | 16 | 699 | 1 | LinearUp-5 |
| 10 | FlyBase Gene Group | FBgg0001102 | UNCLASSIFIED ANTIMICROBIAL PEPTIDES | LinearUp-5 | 0.000925 | 5 | 7 | 699 | 1 | LinearUp-5 |

Showing 1 to 10 of 80 entries

Previous 1 2 3 4 5 ... 8 Next
